# Balancing the scales: Impact of irrigation and pathogen burden on potato blackleg disease and soil microbial communities

**DOI:** 10.1101/2023.07.13.548922

**Authors:** Ciara Keating, Elizabeth Kilbride, Mark A. Stalham, Charlotte Nellist, Joel Milner, Sonia Humphris, Ian Toth, Barbara K. Mable, Umer Zeeshan Ijaz

## Abstract

Understanding the interaction between environmental conditions, crop yields, and soil health is crucial for sustainable agriculture in a changing climate. Management practices to limit disease are a balancing act. For example, in potato production, dry conditions favour common scab (*Streptomyces* spp.) and wet conditions favour blackleg disease (*Pectobacterium* spp.). The exact mechanisms involved and how these link to changes in the soil microbiome is unclear. Our objectives were to test how irrigation management and bacterial pathogen load in potato seed stocks impact: i) crop yields; ii) disease development (blackleg/common scab); and iii) soil microbial community dynamics. We used stocks of seed potatoes with varying *Pectobacterium* levels (Jelly [high load], Jelly [low load], and Estima [Zero – no *Pectobacterium*]). Stocks were grown under four irrigation regimes that differed in the timing and level of watering. The soil microbial communities were profiled using amplicon sequencing at 50% plant emergence and harvest and advanced bioinformatic analyses were used to correlate microbes to treatments and disease symptoms. Irrigation increased blackleg symptoms in the plots planted with stocks with low and high levels of *Pectobacterium* (22-34%) but not in the zero stock (2-6%). Not irrigating increased common scab symptoms (2-5%) and reduced crop yields. Irrigation did not impact the composition of the soil microbiome, but planting stock with a high *Pectobacterium* burden resulted in an increased abundance of *Planctomycetota*, *Anaerolinea*, and *Acidobacteria* species within the microbiome. Ensemble quotient analysis highlighted *Anaerolinea* as highly associated with blackleg symptoms in the field. We conclude that *Pectobacterium* pathogen load within seed stocks could have more substantial effects on soil communities than irrigation regimes.

## Introduction

To meet the growing global demand for food, it has been estimated that agricultural crop production needs to more than double by 2050 (Ray et al., 2012). Production rates are predicted to decrease due to the impact of climate change and requirements for environmentally sustainable production (Ray et al., 2015). Potatoes (*Solanum tuberosum*) are one of the most important food crops, with over 359 million metric tons produced globally (Statista, 2020). However, production yields have stagnated over the past decade and land use for potatoes has decreased (Griffin et al., 2022; Potato production worldwide, 2020). Potatoes are highly susceptible to infection from a plethora of pests and pathogens. These include for example; oomycetes (late blight; *Phytophthora infestans* - Fry, 2020; Yuen, 2021), bacteria (blackleg; *Pectobacterium* spp., - Van Gijsegem et al. 2021, common scab; *Streptomyces* spp. - Braun et al. 2017), potato cyst nematodes (*Globodera* spp. - Price et al. 2021) and viruses (Gray and Power, 2018). These diseases reduce plant growth, the quality of tubers and overall potato production yields. Changing environmental conditions are anticipated to exacerbate disease prevalence and future potato crop production (Haverkort and Verhagen, 2008; Luck et al., 2011; Dahal et al., 2019; Divya et al., 2021). For example, even within the UK, there is extensive variation in sustainable management solutions, which need to be customised to the local environmental conditions. Growers must also make trade-offs between crop value (based on consumer choice and supply chains) and pathogen management. While cultivar selection is largely driven by crop value, there are known to be both climate-induced and cultivar-induced differences in disease susceptibility (Garrett et al., 2009; Ardanov et al., 2016; Velásquez et al., 2018). However, this effect is highly variable and not clearly understood. Thus, a deeper understanding of the impact of management practices, potato stock selection and pathogen load on plant susceptibility and soil diversity would be invaluable.

Soil moisture content is a key parameter in the development of many potato crop diseases. In dry conditions, crops are typically more prone to common scab, *Streptomyces* species (Lapwood and Lewis, 1967; Wale and Sutton, 2004). In contrast, wet conditions promote the development of blackleg disease, *Pectobacterium* spp. (Shock et al., 2007), while a combination of wet, warm and dry conditions favours late blight, *Phytophthora infestans* (Shock et al., 2007). In this work, we will focus on two of these diseases: common scab and blackleg disease.

Common scab disease results in scab-like lesions on potato tubers. Timing is crucial for disease development, with potatoes being the most vulnerable up to six weeks post tuber initiation (i.e. tuber development; Lapwood and Lewis, 1967). Irrigation applied during the period when plants are most susceptible to common scab infection can reduce disease and increase crop yields (Lapwood et al., 1970; Lapwood et al., 1973; Wilson et al., 2001; Djaman et al., 2021), but some authors have noted that this is not a reliable means of control (Dees and Wanner, 2012). Indeed, the mechanisms of control via irrigation or other means (soil pH amendment) are not clearly understood. Moreover, disease control is challenging as many *Streptomyces* spp. are naturally present in the soil environment (Lapwood, 1972).

Blackleg disease of potato stems and soft rot disease of tubers is typically caused by *Pectobacterium atrosepticum* in the UK but may be caused by other *Pectobacterium* and *Dickeya* species elsewhere (Van Gijsegem et al., 2021). A high soil moisture content induces blackleg infection in two ways. Firstly, anaerobic conditions from water logging cause the suppression of plant oxidative stress response mediated defence mechanisms and increase plant cell permeability (Burton and Wigginton, 1970; Pérombelon, 1992; Pérombelon, 2002). Secondly, *P. atrosepticum* is a facultative anaerobe and thus anoxic conditions enable it to proliferate and take advantage of the previously mentioned weaknesses in the plant host (Pérombelon, 2002). *Pectobacterium* species are thought to have limited survival in soil, with disease primarily spread through contaminated seed tubers. There are no chemical control options for blackleg and control of the disease has primarily been through stringent seed lot certification. For example, in the UK there is zero tolerance in the first-generations of progeny seed [pre-basic seed grade] (APHA, 2020; SASA 2023). In terms of management practice, tubers are typically only grown for 5-6 years due to the build-up of *P. atrosepticum* (Pérombelon and Hyman, 1992; Toth et al., 2003a). However, other factors such as irrigation, aerosols and contaminated seed handling equipment may also play a role (de Werra et al., 2020; van der Wolf et al., 2021). Additional management practices during the growing season are not well defined and are less evidenced in the literature, but some practices include rogueing fields to remove crops showing characteristic above-ground wilting associated with blackleg disease (Czajkowski et al., 2011). Blackleg disease development is heavily influenced by climate; thus, management must be modified to local conditions (Pérombelon, 1992). For example, Scotland produces most UK potato seed stock, which are mainly grown without irrigation to avoid blackleg disease, whilst in England irrigation is used to manage common scab, avoid plant desiccation and increase yield in what are mainly ware crops (except in the case of ware crops). This example nicely highlights the balancing act placed on potato growers, where they must trade off management practices with the environmental niches of multiple pathogens.

Management practices to control crop disease could also affect soil fertility and health, which in turn could impact on crop productivity. For example, fertilisation can lead to soil degradation, while mechanical tillage can lead to soil erosion (Carretta et al., 2021). Soil microbial communities are also influenced by agricultural practices such as crop rotations (Chamberlain et al., 2020), soil compaction (Longepierre et al., 2021), tillage and crop choice (Li et al., 2021; Hills et al., 2020). This is important, as the microorganisms in soil are part of a wider interactive web involving animals and plants whose activities contribute to soil fertility, diversity and plant health. These factors are essential for global food production (Schratzberger et al., 2019), but the exact mechanisms of these interactions and how these intricate systems respond to management practices and changing environmental conditions remain unclear (Dubey et al., 2019). Nevertheless, regulating, predicting and manipulating these responses is crucial for healthy soil and global food security in a changing global environment.

We are beginning to recognise that the soil microbiome is intrinsic to ecosystem health, services, and plant, animal and human health (Banerjee and van der Heijden, 2023). Agricultural management practices can disrupt or alter the native soil microbiomes (Bünemann et al., 2006; Tsiafouli et al., 2015). However, the ramifications of such impacts are not clearly understood. Soil and plant microbiome networks are complex and shaped by interactions with surrounding fauna and responses to the abiotic and biotic environment. Despite this complexity, it has been observed that some plants show a bespoke local microbiome that is distinct from the surrounding soil microbiome (DiLegge et al., 2022). In some cases, this microbiome can be modulated using the plant’s defence system for protection against pathogens (Berendsen et al., 2018). The soil microbiome can also play a role in defence against diseases, e.g. in disease-suppressive soils (Schlatter et al., 2017; Mareckova et al., 2022). Which soil microorganisms are involved in the interactions essential for plant health and what variables drive these key microbial taxa remains to be unravelled. A perspective by Toju et al. (2018) discussed this concept, and highlighted the potential for multiple states of healthy soil microbiome (core microbiomes) akin to human gut-associated microbiomes (enterotypes) which, if understood, could allow us to unlock the potential of sustainable agroecosystems.

In this study, our overall aims was to determine whether there are interactions between agricultural management practices, potato seed stocks, crop health and how these link to soil microbial communities. Using agricultural field trials at NIAB Cambridge, UK we tested whether irrigation practices and/or starting pathogen burden (*Pectobacterium* sp.) in different stocks of seed potatoes affected crop yields, disease symptoms (common scab and blackleg) and overall diversity (taxonomic and functional) of the core soil microbial communities. To more specifically quantify differences in ‘key’ microbial taxa or the contributions of rare taxa to the microbiome, we then used differential heat-tree analysis and generalised linear latent variable modelling (GLLVM) to specifically compare microbial community composition in relation to the experimental treatments and used ensemble quotient analysis (EQO) to test whether there was a correlation between disease symptoms (blackleg and common scab) with particular microbial genera.

## Materials and Methods

### Seed Stock Selection

To select appropriate seed stocks with varying *Pectobacterium* levels, we used blackleg-susceptible commercial potato stocks which were tested for pathogen prevalence by SASA (Science and Advice for Scottish Agriculture). Two of the stocks were of the Jelly variety: JellyHigh (Pre-basic Class Generation 3 stock [PB3] with peel containing a mean of 1.47 x 10^3^ colony forming units (cfu)/g of *P. atrosepticum* and 5.74 x 10^4^ cfu/g of *Pectobacterium carotovorum*) and JellyLow (Pre-basic Class Generation 2 stock [PB2] containing a mean of 400 cfu/g of *P. atrosepticum* and 1.88 x 10^3^ cfu/g of *P. carotovorum*). No commercial Jelly stocks available to us were free of *Pectobacterium* and Jelly mini-tubers were unavailable. Therefore, for the stock without *Pectobacterium* we used Estima mini-tubers (EstimaZero) stock [Pre-basic PB1], which were found contain zero *Pectobacterium* species. Note: *Pectobacterium atrosepticum* is the causative agent of blackleg disease, while *Pectobacterium carotovorum* is not typically associated with the disease in the field, although authors have shown potential for disease outcome using inoculation experiments (de Haan et al., 2008).

### Irrigation Regimes

We compared four irrigation treatments: 1) rainfed only (Unirrigated); 2) irrigated when the soil moisture deficit (SMD) reached 40 mm (Irrigation 1); 3) irrigated maintaining SMD < 15 mm during the common scab control period (the 4 weeks following the onset of tuber initiation) and unirrigated for the rest of the season (Irrigation 2); and 4) irrigated maintaining SMD < 15 mm during the common scab control period and then < 25 mm throughout the rest of the season (Irrigation 3). Irrigation was applied using 20-25 mm applications for all irrigation regimes. The total amount of water applied during the season was 150 mm for Irrigation 1 and Irrigation 2, and 251 mm for Irrigation 3. The total rainfall recorded from plant emergence was 202 mm. No plots were protected from rainfall.

### Field Trials

The field-trial experiments were carried out at NIAB, Cambridge (52.2417 °N, 0.0987 °E). The soil profile is included in the Supplementary Methods. The field was randomised into a factorial block design using combinations of the three seed stocks (JellyHigh, JellyLow, and EstimaZero) and four irrigation regimes (Unirrigated, Irrigation 1, Irrigation 2, and Irrigation 3), which were replicated in triplicate plots. Details of the schematic of the experimental design (Supplementary Figure 1) and layout of the plots (Supplementary Datasheet 1) is provided in Supplementary Information. The experiment was planted on the 23^rd^ of April 2020 using 30-40 mm Jelly seed (1206 tuber count/50 kg) and 25-30 mm Estima mini-tubers (2078 tuber count/50 kg) at a within-row spacing of 30 cm in 75 cm rows. The seed was dibbed 12 cm deep into pre-formed ridges, which were raked after planting to re-form the original ridge. Plots were 3.6 m long and four rows wide, with a 2 m pathway between strips of plots. An extra guard row was planted either side of each plot to act an additional buffer against overland and aerial movement of water from the irrigator. Plots were tie-bunded at either end to prevent over-land water flow between plots to minimise the risk of between-plot transfer of blackleg. Post planting (30/04/2020), ammonium nitrate was applied at a rate of 200 kg N/ha and herbicides and fungicides were applied as required to keep the experiment free from weeds and blight. Irrigation was scheduled using the Cambridge University Farm (CUF) Potato Irrigation Scheduling Model based on meteorological data obtained from a Delta-T Devices weather station c. 450 m from the experimental plot. The irrigation was carried out using a diesel engine-driven Briggs VR4 90/400 hosereel and R50 boom equipped with Senninger LDN UP3 Single Pad nozzle dropper pipes to allow discrete irrigation between plots.

### Crop Monitoring and Crop Yield

Plant emergence was recorded by counting the number of plants that had emerged within the two central harvest rows. Emergence was first recorded on the 19^th^ of May 2020 and recorded every 3-4 days thereafter until the 15^th^ of June 2020. The number of plants with initiated tubers in a two-plant sample harvested in every plot were counted every two days, commencing on the 8^th^ of June. At harvest (23-25^th^ September 2020), tubers were graded, counted and weighed in 10 mm increments to provide total tuber yield and volume per hectare.

### Blackleg Incidence and Common Scab Prevalence

Plants were scored for blackleg symptoms on 18th June, 26th June, 6th July, 31st July and 20th of August 2020. Blackleg symptoms were considered as either black decaying lesions on stems or wilted leaves on a single stem. At harvest tubers showing rot symptoms were bagged separately, counted and the weight recorded. A total of 50 tubers from final harvest were selected for common scab assessment, which was determined as the total surface area infected with common scab which was further sorted into the following threshold levels: 0, 1, 5, 10, 15, 20, 25, 30, 40, 50, 60, 70, 80 and 90 % surface area infected (AHDB, 2015).

Complete metadata for the field-trial was collated and included plot number, plot area (based on location in the field – Supplementary Data Files 1-2), seed stock, irrigation regime, rainfall, temperature, irrigation volumes, starting levels of *P. atrosepticum* and of *P. carotovorum* in the starting seed stock), blackleg incidence, blackleg percentage prevalence, the weight of rotted tubers and common scab percentage.

### DNA extraction and amplicon sequencing of the 16S rRNA gene

Soil samples were taken for microbiome analysis at 50% plant emergence (22^nd^ May 2020) and at final harvest (15^th^ September 2020). Samples were taken across treatments (Unirrigated, Irrigation 1, Irrigation 2, and Irrigation 3) and potato stock/pathogen level (JellyHigh, JellyLow, and EstimaZero). Comparisons in microbial communities across time were made using soil (20 samples per plot) sampled from the ridge (near the top of the furrow) at 50% plant emergence (T_E Ridge) and harvest (T_H Ridge). In addition, to determine if there were differences in microbial communities sampled closer to the plant roots, additional samples (20 per plot) were also taken at the bottom of the furrow at harvest only (T_H Root), to avoid damaging roots in the emerging plants (Supplementary Figure 1). The soil was mixed well and stored in cold storage at NIAB before shipping to the University of Glasgow for further homogenisation of sample material by mixing in large sampling bags and taking aliquots for storage at −80°C. DNA was extracted with the Fast DNA™ SPIN Kit for Soil (MP Biomedicals, California, United States of America) using 0.5 g of soil material. Negative controls (nuclease-free water) were additionally passed through the extraction procedure for each set of extractions. DNA quality was assessed by agarose gel electrophoresis and quantified using a QuBit 3 Fluorometer (Thermo-Fisher Scientific, Renfrew, Scotland). Amplicon sequencing was then carried out using the universal bacterial and archaeal primer set (515f and 806r; Caporaso et al., 2012) that contained the Illumina adapter sequence ‘TCGTCGGCAGCGTCAGATGTGTATAAGAGACAG’ on the forward primer and the adapter sequence ‘GTCTCGTGGGCTCGGAGATGTGTATAAGAGACAG’ on the reverse primer. Samples were indexed using the Nextera XT DNA Library Preparation Kit (Illumina Inc., Hayward, California, USA Illumina). Amplicon libraries were pooled and quality checked using a Bioanalyser (Agilent, Santa Clara, California, USA) and RT-qPCR (quantitative reverse transcription polymerase chain reaction) using the Kapa library Quantification kit on an Applied Biosystems StepOne plus system (Applied Biosystems, Massachusetts, USA). Sequencing was carried out using the Illumina technology at the Glasgow Polyomics sequencing facility using a standard flow cell and 600 cycle v3 reagent cartridge for 2 x 300 bp (base pair) reads on an Illumina MiSeq instrument (Illumina Inc., Hayward, California, USA).

### Bioinformatic Analysis

The paired-end reads were demultiplexed and converted to FastQ format sequence files for further analysis. These raw sequences were submitted to the sequence read archive (SRA) database under Bioproject Submission PRJNA992106. A total number of 13,575,540 reads were obtained from 222 samples. Briefly, we used Qiime2 software for the sequencing analysis (Caporaso et al., 2010). Within the Qiime2 framework, the DEBLUR algorithm (Amir et al., 2017) was used for generation of amplicon sequencing variants (ASVs), after quality trimming the reads with a Phred quality score of 20. The final ASVs (*p* = 33,464 ASVs for *n* = 222 samples) were then classified against the SILVA v138.1 SSU Ref NR database (Quast et al., 2012). The abundance table was then combined with the taxonomy to produce a biom file, on which downstream statistics were applied. In addition, predictive functional analysis was carried out using PICRUSt2 software within the Qiime2 framework, which produced sample level KEGG orthologs and MetaCyc pathways as biom and functional.tsv files (Douglas et al., 2020). As a pre-processing step, we removed typical eukaryotic contaminants such as Mitochondria, Chloroplasts, and any ASVs unassigned at all levels, as per recommendations given at https://docs.qiime2.org/2022.8/tutorials/filtering/. Moreover, singletons and samples with reads < 5,000 were excluded from analysis. This resulted in 31,114 ASVs from 208 samples. Prior to the downstream statistical analysis, we used the ‘decontam’ package in R to remove contaminants by comparing the prevalence of ASVs in the negative controls to true samples (Davis et al., 2018).

#### Abundant and Core Microbial Genera

The top 25 most abundant microbial families as a proportion of total relative abundance per sample were visualised using ggplot2 (Wickham, 2016). We considered the common ‘core’ microbiome to constitute the genera present in all field-trial samples (with >1% compositional abundance in at least 85% of samples). The common core microbiome has been summarised in a review by Shetty et al. (2017). Analysis was conducted in R (R version 4.1.3: R Core Team, 2022) using ‘phyloseq’ (McMurdie and Holmes, 2013). The core microbiome was calculated using R’s microbiome package (Lahti et al., 2017) based on the preliminary work of Jalanka-Tuovinen et al. (2011).

### Statistical Analysis

Statistical analysis methods are provided in the Supplementary Material.

## Results

### Crop Yields

The choice of potato stock (with associated pathogen levels) and irrigation regime both impacted crop yield. Not irrigating reduced the tuber yield for all potato stocks, particularly for the EstimaZero mini-tubers (Table 1). Irrigation regime 1 (irrigated when SMD reached 40 mm) resulted in the highest yield for both Jelly stocks (JellyHigh 69 tons/HA; JellyLow 67.4 tons/HA). In contrast, irrigation regime 3 (irrigated maintaining SMD < 15 mm during the common scab control period and then < 25 mm throughout the rest of the season) resulted in the highest yield for the EstimaZero stock, which was the highest tuber yield observed for all stocks and treatments (75 tons/HA).

**Table 1.** Potato crop yields from the experimental field trials at harvest. Tuber yield information for each potato stock (JellyHigh, JellyLow and EstimaZero) and each irrigation regime (Unirrigated, Irrigation 1, Irrigation 2, and Irrigation 3). Tuber yield was calculated by the total number of tubers per hectare (HA) and the tuber yield in tonnes per hectare. Results of the statistical analysis (P values) testing the impact of ‘Stock’, ‘Irrigation’, and ‘Stock and Irrigation’ are shown at the bottom of the table. Values are significant when P < 0.05.

| STOCK | IRRIGATION | Total No.<br>Of Tubers<br>(000/HA) | Total<br>Tuber Yield<br>(T/HA) |
| --- | --- | --- | --- |
| JELLYHIGH | Unirrigated | 267 | 45.2 |
|  | Irrigation 1 | 349 | 69.0 |
|  | **Irrigation 2** | 277 | 50.0 |
|  | **Irrigation 3** | 299 | 63.8 |
| **JELLYLOW** | **Unirrigated** | 339 | 49.9 |
|  | **Irrigation 1** | 369 | 67.4 |
|  | **Irrigation 2** | 341 | 54.5 |
|  | **Irrigation 3** | 308 | 64.9 |
| **ESTIMAZERO** | **Unirrigated** | 264 | 29.8 |
|  | **Irrigation 1** | 324 | 60.5 |
|  | **Irrigation 2** | 394 | 59.0 |
|  | **Irrigation 3** | 401 | 75.0 |
|  | **Fprob Stock** | 0.020 | 0.501 |
|  | **Fprob Irrigation** | 0.032 | <0.001 |
|  | **Fprob Stock\*Irrigation** | 0.011 | 0.006 |

### Disease Prevalence

The Jelly stocks with both high and low initial *Pectobacterium* loads both showed a high percentage of blackleg symptoms at the end of the field trial (Table 2). The percentage of blackleg symptoms was strongly impacted by irrigation regime, with the unirrigated treatment showing the lowest symptom prevalence (16.7% JellyHigh and 18.1% JellyLow) and irrigation 3 showing the highest symptom prevalence (34% for both Jelly stocks), although there was some variation among plots (Supplementary Table 1). Examining the blackleg symptoms over time highlights that disease incidence developed at a similar rate in the stock with low starting levels of *Pectobacterium* (JellyLow) as compared to the stock with high starting levels (JellyHigh) (Figure 1). This trend was exacerbated in the irrigation treatments (Irrigation 1, Irrigation 2 and Irrigation 3). In contrast, the EstimaZero with no initial *Pectobacterium* detected showed very low levels of blackleg symptoms across the entire experiment (reaching only 0.7% in the unirrigated treatment and ranging from 2.8-5.6% in the irrigated treatments at harvest; Table 2). Despite substantial blackleg symptoms recorded in the field for the Jelly stocks, overall, the degree of tuber rotting at harvest was low, with < 1.5% of rotted tubers (Supplementary Table 2). Moreover, no significant difference was observed with respect to irrigation or potato stock on the proportion of rotting tubers.

**Figure 1.**
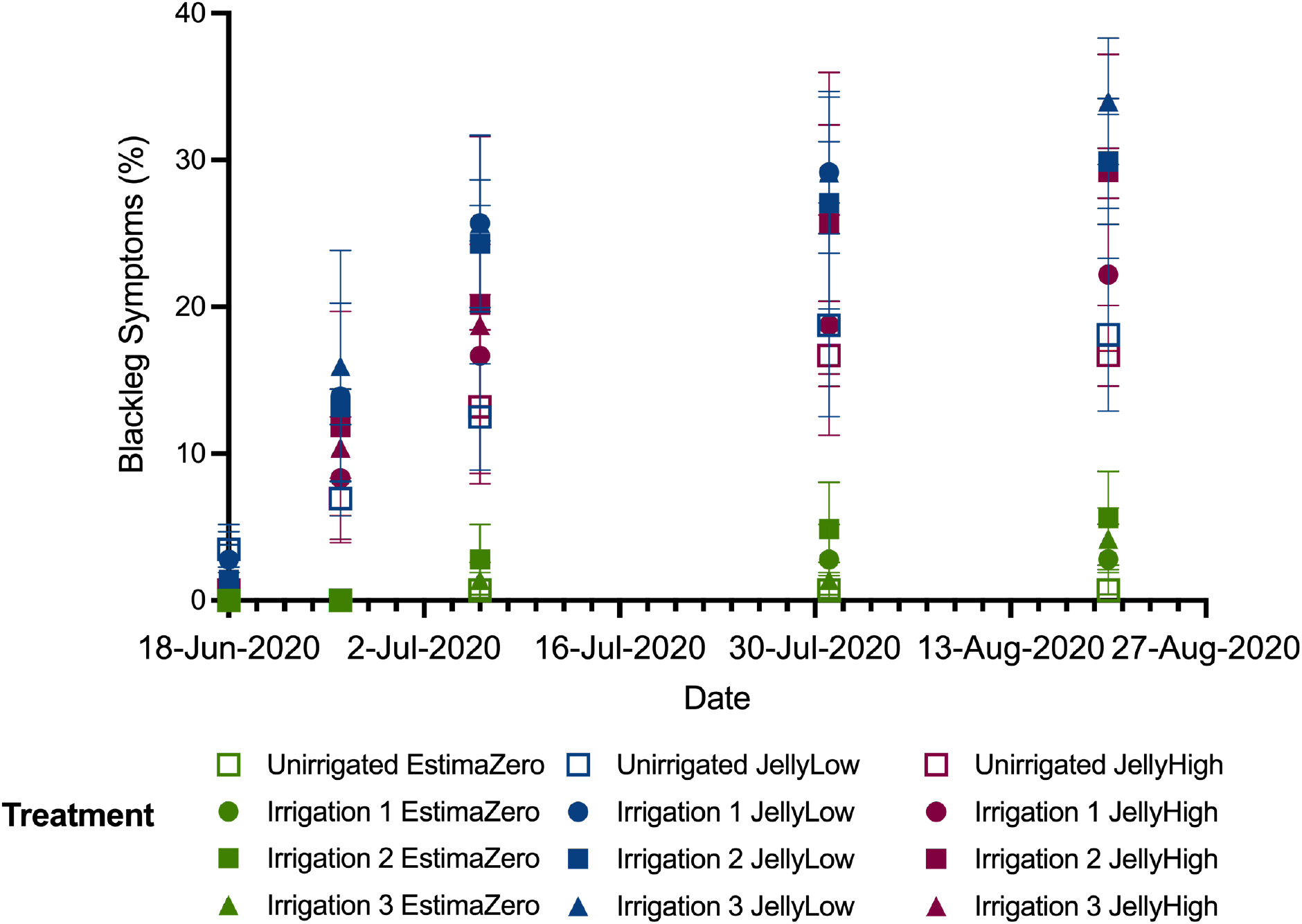
Blackleg symptoms (%) plotted against time from plant emergence (18/06/2020) to plant harvest (27/08/2020). Symptoms are based on the average of blackleg symptoms per plot based on the incidences of recorded symptoms in the field. Values are shown by stock (EstimaZero, JellyLow and JellyHigh) and irrigation regimes (Unirrigated, Irrigation 1, Irrigation 2 and Irrigation 3). Standard deviation values are shown in grey lines.

**Table 2.**
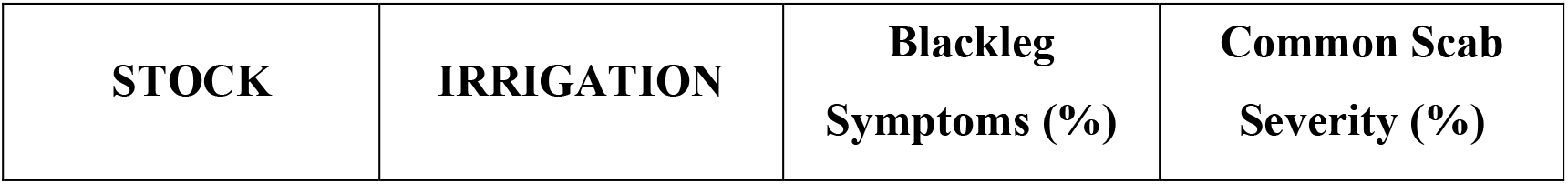
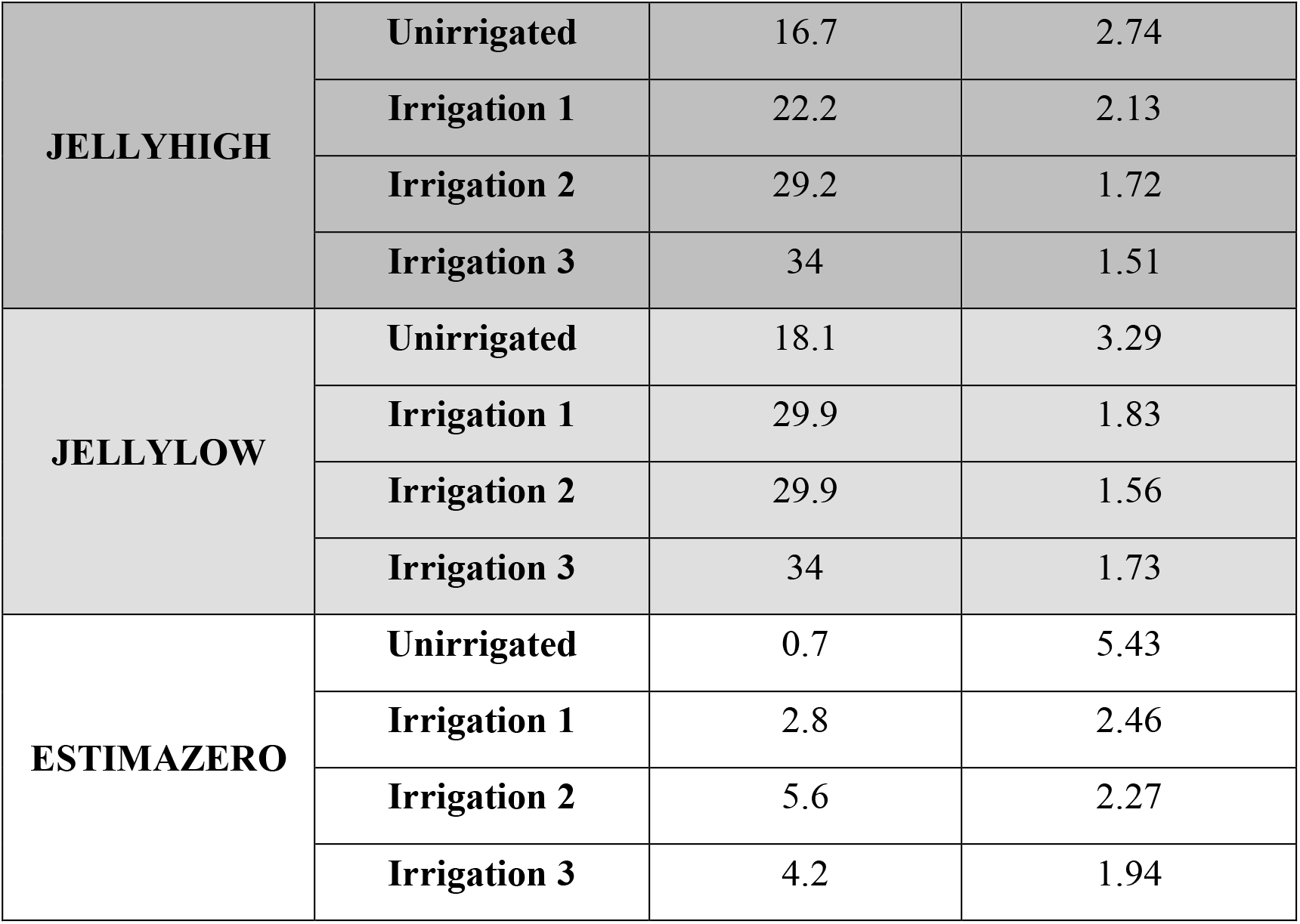
Blackleg and Common Scab symptoms at the end of the experimental trial, showing the percentage of the crop showing blackleg symptoms on 20^th^ August 2020 and the percentage common scab severity from tubers from final harvest (23-25^th^ September 2020) for the three potato stocks (JellyHigh, JellyLow and EstimaZero) and the four irrigation regimes (Unirrigated, Irrigation 1, Irrigation 2, and Irrigation 3).

While irrigation increased blackleg disease symptoms, a lack of irrigation (rainfed only) strongly impacted common scab severity in the tubers upon harvest (Table 2). The unirrigated treatments showed the highest common scab levels, and the Estima tubers were the most affected (5.43% EstimaZero, 3.29% JellyLow and 2.74% JellyHigh). In general, tubers from irrigation regime 3 (irrigated maintaining SMD < 15 mm during the common scab control period and then < 25 mm throughout the rest of the season) showed the lowest common scab severity (1.94% EstimaZero, 1.73% JellyLow and 1.51% JellyHigh – Figure 1; Table 2).

### Dominant Microbial Community Composition and Core Microbiome

The 25 most abundant microbial families in the soil communities were remarkably stable (Figure 2) in relation to initial *Pectobacterium* levels (Zero, Low and High), potato variety (Jelly seed or Estima mini-tubers) and irrigation regime (Unirrigated, Irrigation_1, Irrigation_2 and Irrigation_3). Some of the most abundant taxa were species within the *Vicinamibacterales* (~ 20%), *Nitrososphaeraceae* (~ 12%), *Nocardiodaceae* (~ 8%) *and Sphingomonodaceae* (~ 8%) families (Figure 2). Although we did observe some shifts in relative abundance across time in the ridge samples (T-E – 50% plant emergence and T-H – plant harvest) and in the relative abundances of the top 25 families between the ridge and root samples at harvest, the pattern of changes were similar across stocks and irrigation treatments (Figure 2). Based on a threshold of presence in at least 85% of samples, we observed that 65 genera were part of the ‘core microbiome’ that did not change in relation to treatments (Supplementary Figure 3; Supplementary Table 4). Of these, *Vicinibacterales*, *Nocardioides*, *Pyrinomonadaceae* (RB41) and *Chloroflexi* (KD4-96) species showed the highest prevalence (75 - 100%) and the highest number of amplicon sequencing variants (ASVs) detected (298-1362 ASVs).

**Figure 2.**
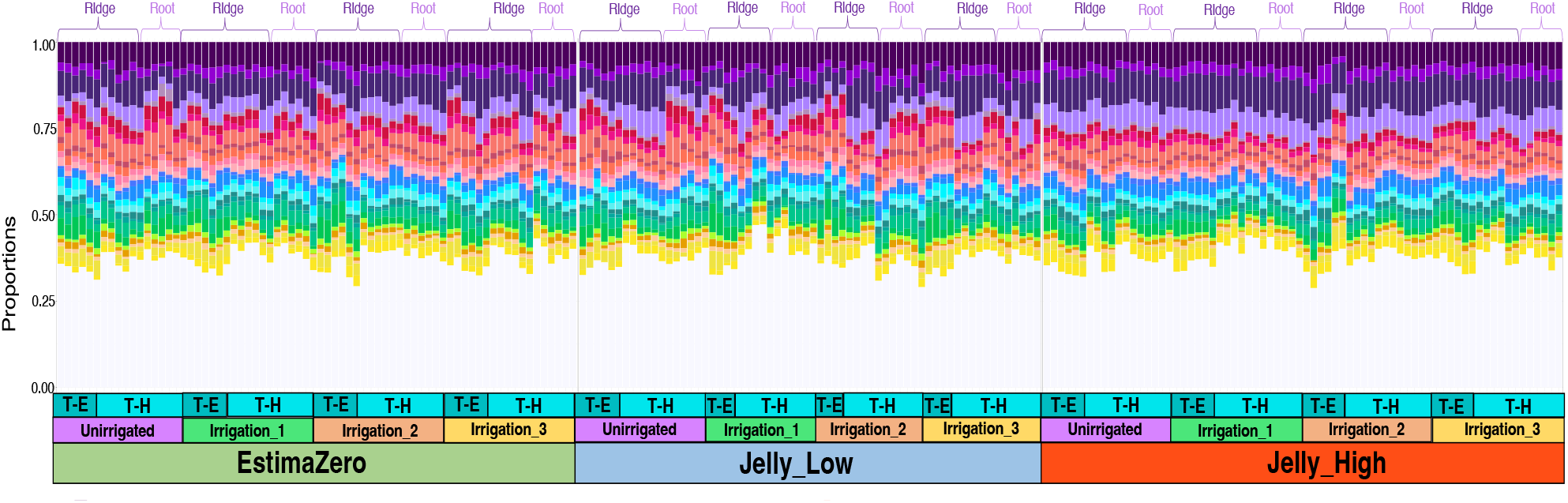
Stacked vertical bar chart comparing the relative abundance of the top 25 microbial families found in soil across treatments (irrigation regime), stocks/ pathogen levels (JellyHigh, JellyLow, Estima), time (50% plant emergence T_E, or harvest, T-H) and whether samples were taken from the ridge (background soil community at both timepoints) or root (rhizosphere at harvest). Colours represent the relative abundance of each taxonomic family. See Supplementary Table 3 for the taxonomic description). The white bars indicate the contribution of microbial families that were not represented in the top 25.

### Alpha and Beta Diversity

Alpha diversity represents the number and evenness of different ASVs *within* a single sample, while beta diversity is a measure of similarity or dissimilarity *between* ASVs from two communities (Finotello et al., 2018). In addition to considering ASVs (taxonomic diversity) we also considered functional diversity, which was based on the number and evenness of KEGG orthologs (KOs) within a single sample (alpha diversity) and the dissimilarity of KOs between two communities (beta diversity).

#### Irrigation

At 50% plant emergence, the irrigation management practice did not significantly impact taxonomic or functional microbial diversity either within or between treatments (Supplementary Figure 4-5).

At harvest, there were no significant differences in taxonomic diversity within treatments (Supplementary Figure 6A). However, there were differences in taxonomic diversity between treatments, with samples showing some clustering with respect to Irrigation 2 and whether samples were from the root or ridge (Supplementary Figure 6B; Dim 1 – 6.85%, Dim 2 – 2.9%). At harvest, functional diversity within treatments was altered, with more unique KOs detected in the Irrigation 1 ridge samples, which was significantly higher (0.01 or more) than the ridge samples from Unirrigated, Irrigation 2 and Irrigation 3 regimes (Supplementary Figure 7A). We also observed differences with respect to functional diversity between treatments, with samples clustering according to irrigation regime (particularly Unirrigated and Irrigation 2) and whether samples were from root or ridge (Supplementary Figure 7B; Dim 1 – 77.89%, Dim 2 – 5.16%).

#### Potato Stock

In the above analysis, the taxonomic and functional diversity of the microbial communities between treatments indicated that samples from the same potato stock were highly similar irrespective of irrigation regime. We, therefore, repeated the analysis using potato stock as the grouping variable. At 50% plant emergence, taxonomic diversity was significantly different within treatments. For example, JellyHigh treatments showed both an increased number of ASVs (richness values of ~480-920) and a more balanced distribution in the abundance of these ASVs (evenness values of ~0.935-0.975), while JellyLow showed reduced values, with richness values of 400-550 and evenness values of 0.90-0.96 – Figure 3A). It must be noted that the scale of these changes was relatively low, with evenness values ranging from 0.90-1.0 and richness values ranging from 400-900 (Figure 3A). There were also significant differences in taxonomic diversity between treatments. JellyHigh samples with a high pathogen burden were highly similar and clustered together, whereas in general, the low (JellyLow) and zero (EstimaZero) stocks were more similar to each other and dissimilar to the JellyHigh (Figure 3B; Dim1 – 8.7%, Dim2 – 4.19%). Some Estima samples (from plots 3-4, 3-8, 3-9, 3-11) clustered within this JellyHigh cluster. At harvest, there were fewer differences in taxonomic diversity within treatments. However, values for JellyLow root samples were lower than the other treatments, with evenness values of 0.95 and richness values of 450 (Figure 4A). At harvest, we again observed significant differences in taxonomic diversity between treatments, with EstimaZero and JellyLow samples showing greater similarity, while JellyHigh samples were more dissimilar and formed a distinct cluster (Figure 4B; Dim1 – 6.85%, Dim2 – 2.9%). Interestingly, while some JellyLow/EstimaZero samples clustered with the JellyHigh samples, no JellyHigh were observed outside the JellyHigh signature cluster (Figure 4B). When we considered the taxonomic diversity within treatments over time (ridge samples only), evenness values decreased over time in the EstimaZero and JellyHigh but were increased in the JellyLow (Supplementary Figure 8A), while richness values increased over time in all but the JellyHigh samples (Supplementary Figure 8A). No distinct temporal pattern was observed in taxonomic diversity between treatments (Supplementary Figure 8B).

**Figure 3.**
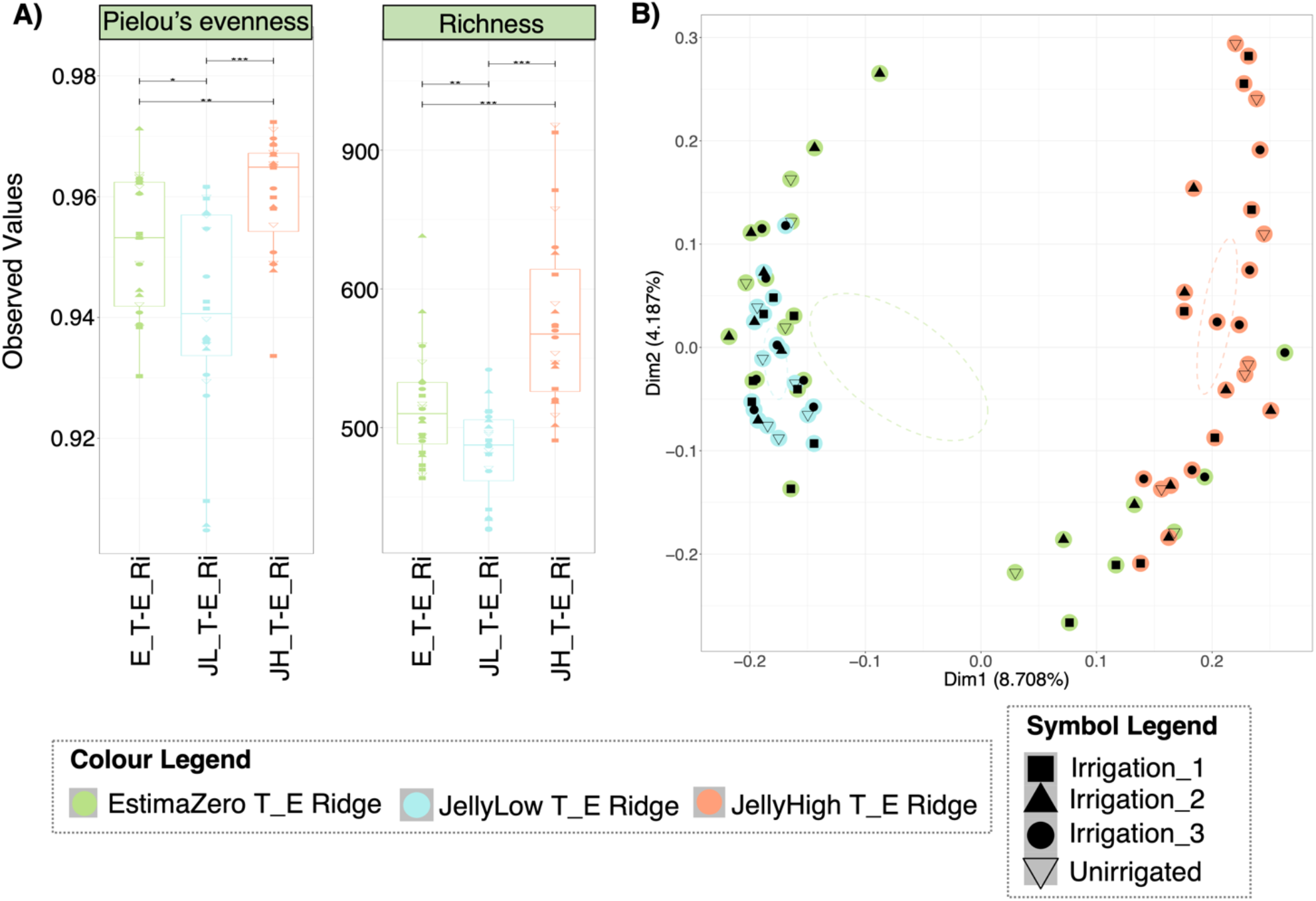
Taxonomic diversity of the soil microbiomes at 50% plant emergence (T_E). Diversity is influenced by potato stock *Pectobacterium* levels and potentially potato variety. Colours represent potato stock (EstimaZero – E, JellyLow – JL, and JellyHigh – JH), time-point (50% plant emergence – T_E, and soil sample type (Ridge – Ri). Symbols represent the irrigation regimes (Unirrigated, Irrigation_1, Irrigation_2, and Irrigation 3). **A)** shows the relative distribution of taxa (Pielou’s evenness) and diversity (Richness) within the communities. **B)** Principal coordinate analysis (PCoA) based on Bray-Curtis distance of beta diversity dissimilarity between the communities.

**Figure 4.**
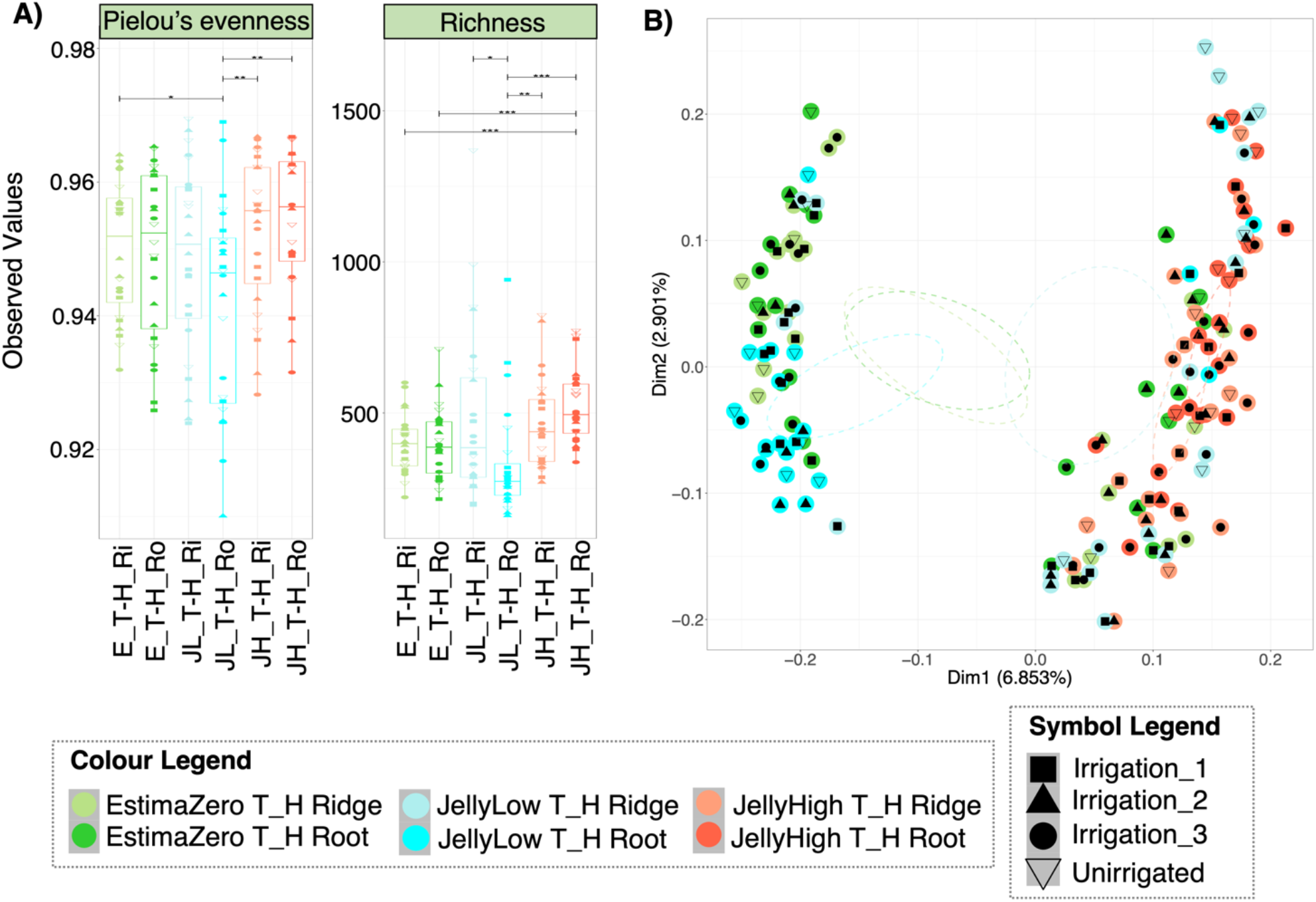
Taxonomic diversity of the soil microbiomes at plant harvest (T_H). Diversity is influenced by potato stock *Pectobacterium* levels and potentially potato variety. Colours represent potato stock (EstimaZero – E, JellyLow – JL, and JellyHigh – JH), time-point (plant harvest – T_H, and soil sample type (Ridge – Ri, Root – Ro). Symbols represent the irrigation regimes (Unirrigated, Irrigation_1, Irrigation_2, and Irrigation 3). **A)** shows the relative distribution of taxa (Pielou’s evenness) and diversity (Richness) within the communities. **B)** Principal coordinate analysis (PCoA) based on Bray-Curtis distance of beta diversity dissimilarity between the communities.

In terms of functional diversity within treatments, at 50% plant emergence and at plant harvest, the distribution in the abundance of KEGG Orthologs (KOs) was lower in the JellyHigh samples (evenness), and the number of detected functions was slightly higher in the JellyHigh samples (Supplementary Figure 9A, Supplementary Figure 10A). Again, the scale of these changes was very low (differences of 0.01 for evenness and 100 for richness). At 50% plant emergence, functional diversity between treatments showed some clustering with respect to potato stock (Supplementary Figure 9B; Dim 1 – 74.48, Dim2 – 7.29%). At harvest, this was more pronounced with samples clustering according to potato stock and sample type (Ridge or Root) (Supplementary Figure 10B; Dim 1 – 77.29, Dim2 – 5.17%). However, these were not as clearly differentiated as observed in the taxonomic diversity plots (Figure 4B; Dim1 – 6.85%, Dim2 – 2.9%). When we considered the functional diversity within treatments over time (ridge samples only), evenness values increased in the EstimaZero and JellyLow, but reduced in the JellyLow (Supplementary Figure 11A), while richness values increased over time in all but the JellyHigh samples, which remained stable (Supplementary Figure 8A). Functional diversity between treatments showed that samples loosely clustered according to time-point and potato stock (Supplementary Figure 11B).

#### Statistical Analysis of Taxonomic Diversity Between Treatments

For the taxonomic diversity between treatments (beta diversity), we also implemented permutational multivariate analysis of variance (PERMANOVA) to determine what categorical variables could explain dissimilarity between such treatments. We determined that potato stock and plot number were highly significant variables (P = 0.001), explaining 8.3% and 43% of the variance between treatments, respectively. The irrigation regime was also a significant variable (P = 0.043; R2 4.5%). At harvest, potato stock, plot area, and plot number were again highly significant (P = 0.001), and sample type (Root or Ridge; P = 0.007) and explained 3.4%, 6%, 17.85%, and 0.9% of the variance, respectively. Irrigation was also significant (P = 0.007; R2 2.4%). Notably, the remaining residual variation was 69%, suggesting that the experimental variables explained a relatively low amount of variation. PERMANOVA analysis of the ridge samples over time showed that time was also a highly significant variable (P = 0.001; R2 1.9%). Redundancy analysis (RDA) with forward selection and subsequent PERMANOVA analysis revealed that blackleg and common scab symptoms per plot were highly significant between the ridge samples from EstimaZero, JellyLow and JellyHigh stocks (P < 0.001), explaining 42% and 17% of the variance, respectively. Irrigation volume was determined not to be significant (P = 0.138). The same pattern was observed in the root samples, although here irrigation volume was slightly significant (P = 0.049).

### Contribution of Rare Taxa to Beta-Diversity

We further assessed how ‘Rare’ microbial taxa (< 1% relative abundance) contribute to taxonomic diversity between potato stocks. We found that all taxa at the ASV-level fell into the ‘Rare’ category and at this level, no ASVs were categorised as ‘Abundant’ (> 1% relative abundance) or ‘Conditionally Rare’ (ASVs with max:min > 100 across groups) [data not shown]. The majority of ASVs were ‘Persistently Rare’ (ASVs with a maximum relative abundance < 5 times their minimum value [max:min<=5]) with some ‘Other Rare’ [ASVs whose abundances were outwith the thresholds for ‘Conditionally Rare’ and ‘Persistently Rare’] ASVs (data not shown). ‘Persistently Rare’ ASVs contributed 42-95% to taxonomic diversity between potato stocks, and ‘Other Rare’ ASVs contributed 10-55% (Supplementary Figure 9). The contribution of ‘Persistently Rare’ taxa to taxonomic diversity increased over time in the EstimaZero and JellyHigh samples but decreased in the JellyLow, while the opposite was observed for the contribution of ‘Other Rare’ ASVs (Supplementary Figure 12).

### Differential Microbial Taxa Associated with Irrigation Management Practice

To assess the impact of irrigation regimes on microbial communities, we first implemented differential heat tree analysis, which determines microbial taxa with a Log_2_ fold difference in abundance in pair-wise comparisons between irrigation regimes. This analysis revealed that no taxa showed a Log_2_ fold difference between irrigation regimes at 50% plant emergence or at harvest (Supplementary Figures 13-16). We then applied a generalised linear latent variable model (GLLVM) to the dataset to find individual microbial genera either positively or negatively associated with each irrigation regime. At 50% plant emergence, we found no significant associations between microbial taxa and irrigation regime (data not shown). However, at harvest, we found that seven microbial genera (out of the top 50 genera) were positively associated with a lack of irrigation in the root samples. These included *Massilia*, *Sphingomonas*, *Gemmatimonas*, *Streptomyces, Microlunatus*, *Nocardiodes*, and *Mycobacterium* genera (Figure 5), while in the ridge sample 16 microbial genera (out of the top 50 genera) were negatively associated with Irrigation 3. These included *Acidimicrobiia*, *Chthoniobacter*, *Streptomyces*, *Sphingomonas*, *Nocardiodes*, and *Vicinimibacteraceae* (Supplementary Figure 17).

**Figure 5.**
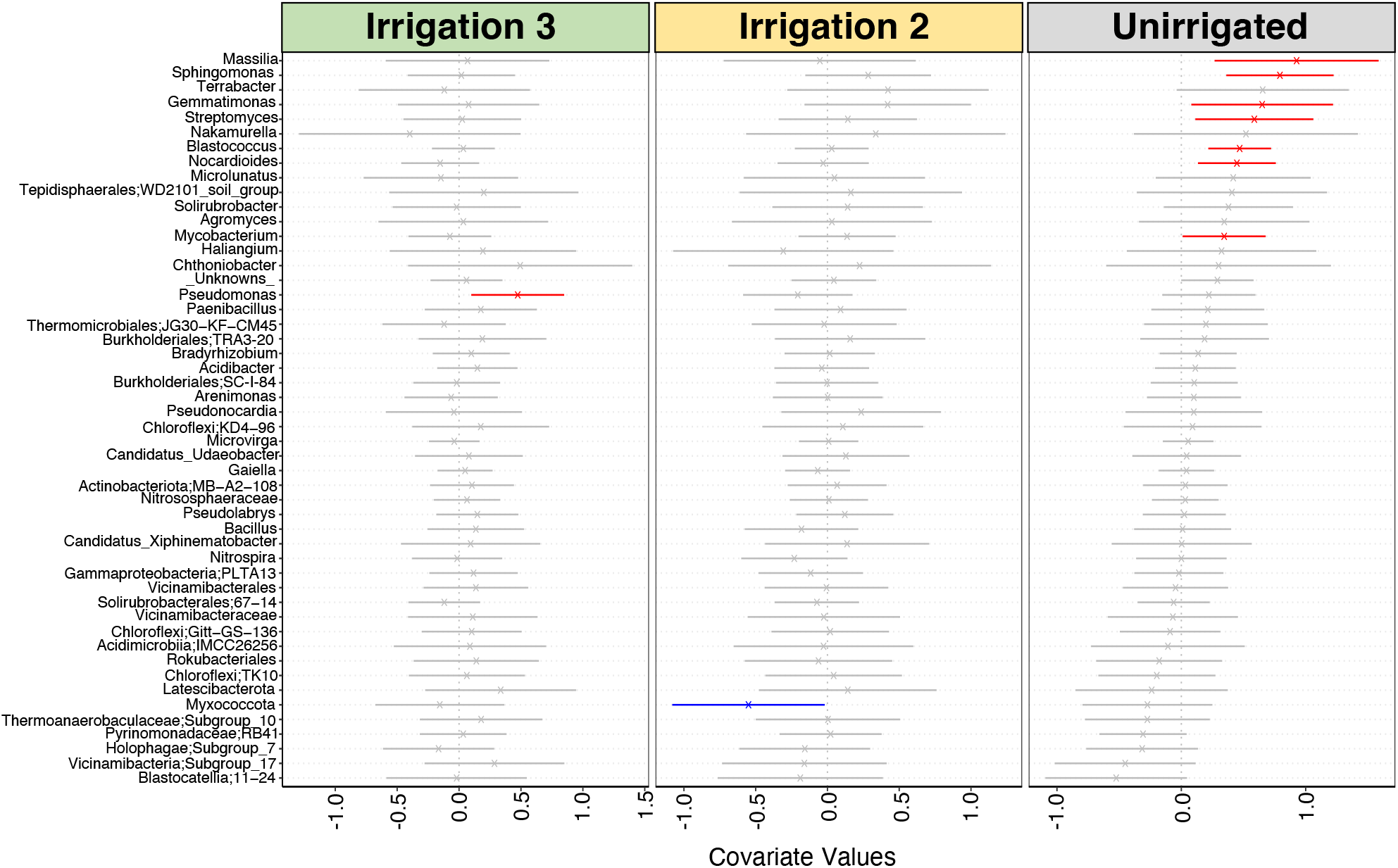
Plot highlighting the impact of ‘Irrigation Regime’ on the abundance of microbial taxa in the potato harvest (T_H) root soil samples based on the generalised linear latent variable model (GLVMM) analysis. The microbial genera names are indicated on the y-axis and the environmental covariate values are indicated on the x-axis, which in this case is (e.g. Unirrigated, Irrigation_1, and Irrigation_3 as compared to the Irrigation 1 treatment). The values range from positive, neutral to negative associations with the specific taxa on the y-axis. The red lines show taxa that are positively associated with the indicated environmental covariate, and the blue lines show taxa that are negatively associated; the lines in grey are not significantly different. Note: Irrigation 1 treatment is not displayed as the analysis uses this treatment as the reference. The results showing the analysis excluding the Unirrigated treatment instead are shown in Supplementary Figure 18.

### Differential Microbial Taxa Associated with Potato Stock and Time

We repeated the above analysis with potato stock as the grouping variable. The differential heat tree analysis revealed substantial differentiation in the abundance of particular taxonomic groups between potato stocks at both time points (T_E and T_H) and in root and ridge samples. At 50% plant emergence, the soil planted with potato stock with a high *Pectobacterium* load (JellyHigh) showed Log_2_ fold increases in Planctomycetota phylum (*Phycisphaeae* [WD2101 soil group], *Pirellulales* [*Pirellula*] and *Gemmatales* [*Gemmatela*]), Chloroflexi phylum (*Anaerolineae* [*Caldilineales*, SBR1031, A4B]) and Acidobacteria phylum (*Vincinamibacteraceae* members) as compared to both the JellyLow and Estima stocks (Figure 6A). In contrast, there were almost no differences between the low (JellyLow) and zero (EstimaZero) stocks. This trend was also observed in the harvest root samples (Figure 6B). However, in the ridge samples at potato harvest, the JellyLow showed significantly increased *Planctomycetota*, *Vicinibacteria*, and *Anaerolinea* as compared to the EstimaZero stock and showed no difference in taxa with the JellyHigh (Figure 6C).

**Figure 6.**
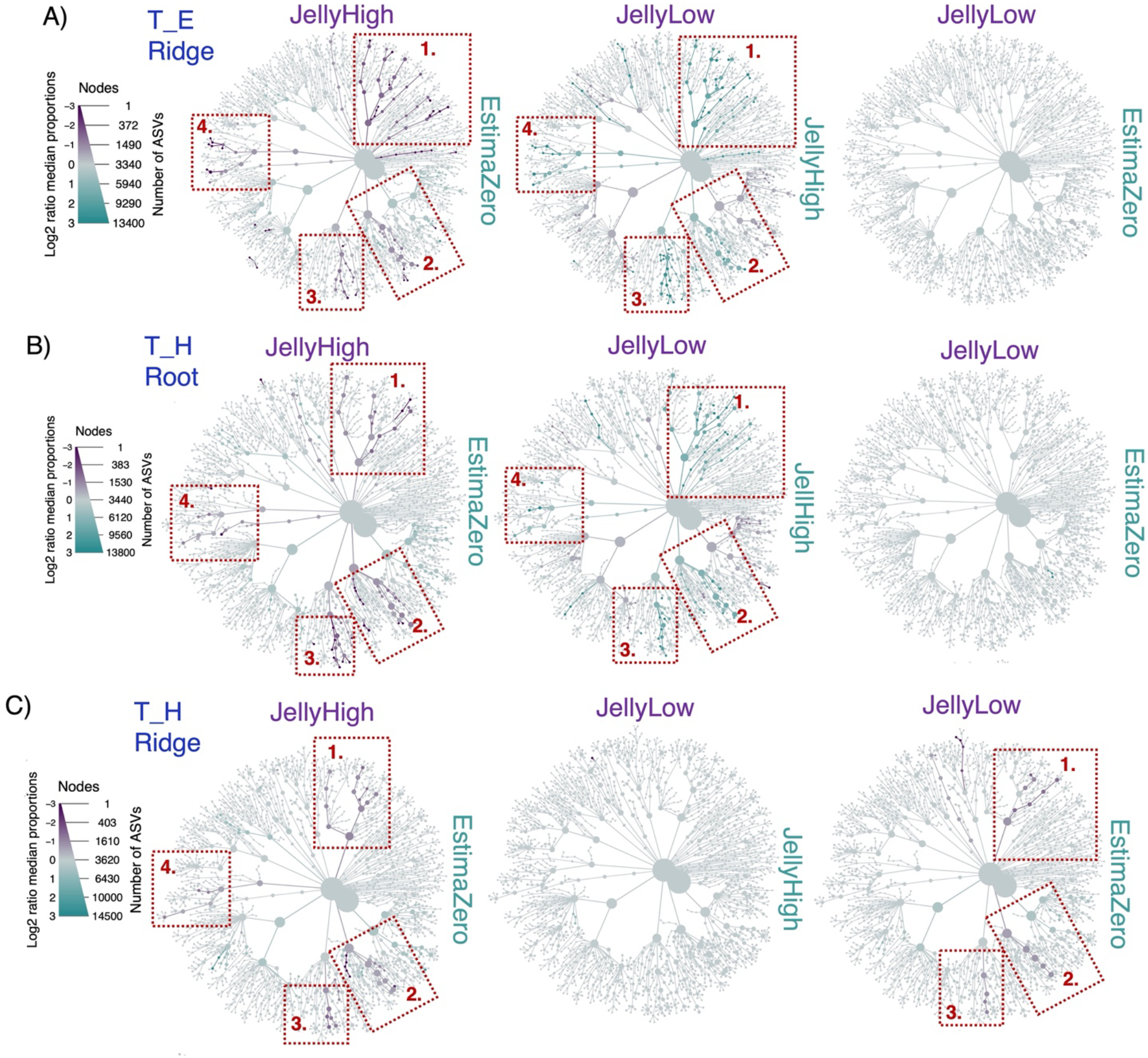
Heat tree analysis to visualise the differential taxa between potato stock soil microbiomes. A high initial pathogen burden in the potato seed stocks impacts the ridge and root soil microbial communities. The full legend for the tree branches is shown in Supplementary Figure 13. In each plot four red boxes are highlighted, which correspond to: 1. *Planctomycetota*; 2. *Vicinibacteria;* 3. *Anaerolinea*; and 4. *Verrucomicrobiae*. The size of the branch nodes shows the number of ASVs, while the differential Log_2_ ratio median proportion is shown by the colours. The figures show a comparison between JellyHigh (purple) compared to EstimaZero (blue), JellyLow (purple) compared to JellyHigh (blue), and JellyLow (purple) compared to EstimaZero (blue) **A)** at the 50% plant emergence (T_E) sampling period of the ridge microbiome; **B)** at the harvest (T_H) sampling period of the ridge microbiome; and **C)** at the harvest (T_H) sampling period of the root microbiome.

Using the GLLVM analysis we further observed that at 50% plant emergence 16 (out of the top 50) microbial genera were positively associated with the stock with a high pathogen burden [JellyHigh] (Supplementary Figure 19). The most positively associated were *Chthoniobacter*, *Anaerolineae*, *Planctomycetota*, *Tepidisphaerales*, *Pirellula*, and *Vicinamibacteria* species (Supplementary Figure 19). Eight of these were also positively associated with the JellyLow samples. Interestingly, at harvest, we observed a different pattern in the JellyLow root samples, which showed more negative or neutral associations with microbial genera that were typically positively associated with the JellyHigh samples (e.g. *Chthoniobacter* and *Vicinamibacteria*; Supplementary Figure 20). Typically, genera that were positively associated with JellyHigh or JellyLow (e.g. *Planctomycetota*, *Anaerolineae*, *Tepidisphaerales*, *Latescibacterota*, and *Vicinamibacteria*) were negatively associated with the EstimaZero group (Supplementary Figure 21).

The GLLVM analysis on temporal samples (ridge samples from 50% plant emergence and harvest) further revealed that *Planctomycetota* (OM190), *Anaerolinea* (SBR1031, A4b, and RBG-13), *Haliangium*, *Pseudocardia*, *Latescibacterota*, *Pirellula*, and *Pseudomonas* were all positively associated with the potato harvest time point (Supplementary Figure 21). In contrast, the following taxa were negatively associated with the potato harvest time point; *Massilia*, *Sphingomonas*, *Flavobacterium*, *Devosia*, *Arenomonas*, and *Acidobacteria* (Supplementary Figure 22).

### Microbial Taxa Associated with Blackleg and Common Scab Disease Symptoms

Finally, we wanted to determine which microbial genera were correlated to blackleg disease symptoms or common scab disease symptoms at potato harvest. This was achieved using Ensemble Quotient Optimisation (EQO), which returns the subset of microbial genera (ensemble) associated with a continuous variable. In the ridge samples, approximately eight genera comprised the minimum subset correlated with blackleg disease symptoms. Within these, the *Anaerolineae* (SBR1031) consistently showed increased relative abundance with increasing blackleg symptoms (Figure 7). A clear trend was seen between increasing percentage blackleg symptoms and the increasing abundance of the minimum subset of taxa identified by EQO. This contrasts with the patterns observed in the harvest root samples, where there was no trend indicating a key microbe to blackleg disease prevalence, with many genera in the subset (12-20) present across low to high blackleg symptoms (Supplementary Figure 23). For common scab symptoms, there was also no trend correlating key taxa to common scab symptoms, with most genera of the subset found across low to high common scab symptoms (Supplementary Figure 24). A similar result was observed in the root samples, though the relative abundance of *Terrabacter* and *Saccharamonidales* appeared to increase with increasing common scab symptoms (Supplementary Figure 25).

**Figure 7.**
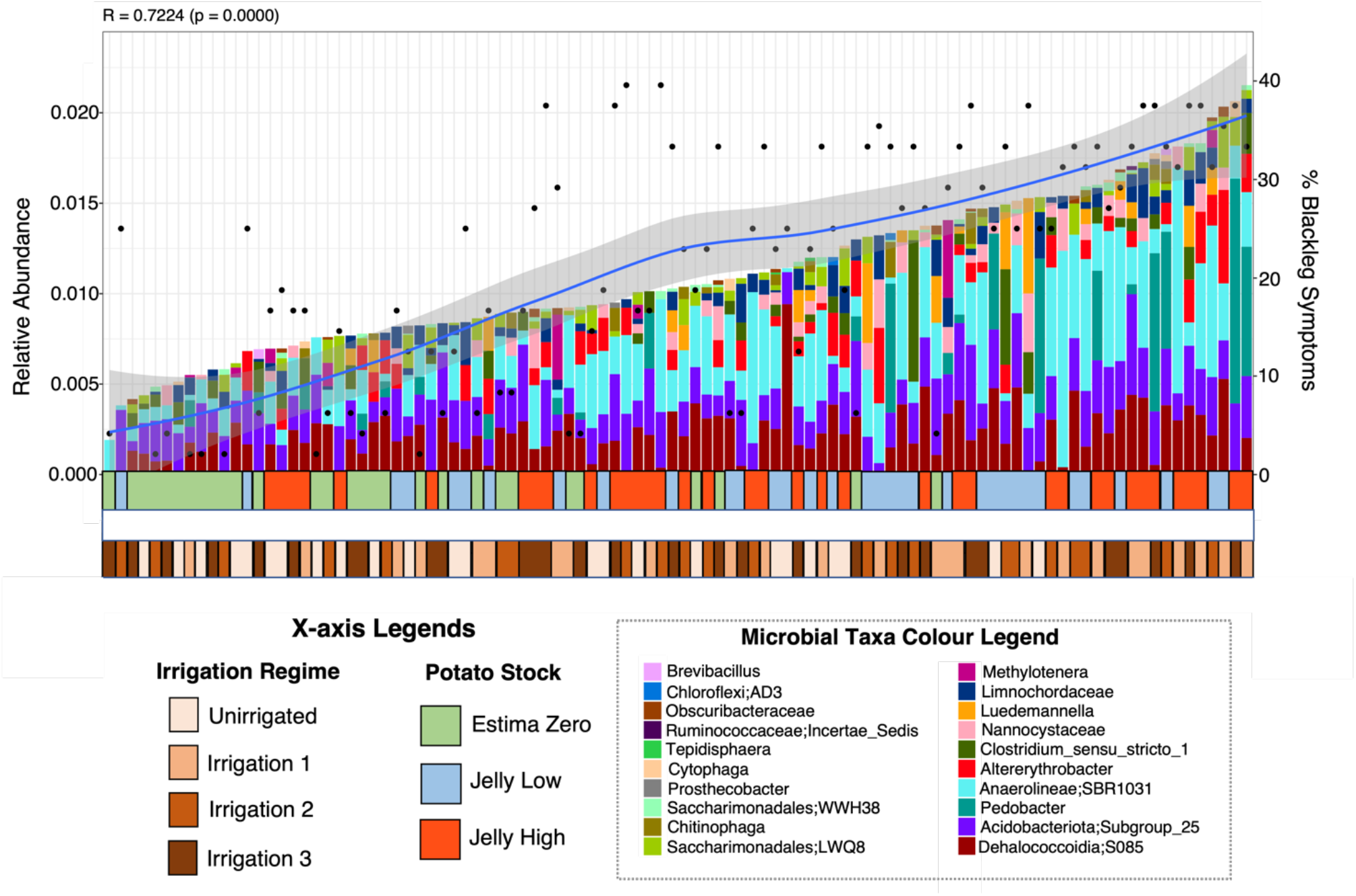
Minimum subset of amplicon sequencing variants (ASVs) associated with percentage blackleg symptoms during the field trial, based on the Ensemble Quotient Optimisation (EQO) analysis, considering the at most 20 microbial taxa for the ridge samples at harvest. The left y-axis shows the relative abundance of the microbial taxa, and the right y-axis shows the percentage blackleg symptoms observed. The x-axis shows the sample identity, with colour legends for potato stock (Green EstimaZero, Blue JellyLow, and Red JellyHigh) and the irrigation regimes (Unirrigated, Irrigation 1, Irrigation 2, and Irrigation 3).

## Discussion

The factors underlying disease development in food crops like potatoes are a global issue. Management practices and climate conditions can mitigate or exacerbate disease symptoms, influence crop yields and perturb the supporting microbial communities (Chamberlain et al., 2020; Carretta et al., 2021; Longepierre et al., 2021). In this study, we assessed the influence of *Pectobacterium* load, seed stock and irrigation on potato yields, common scab and blackleg disease symptoms and the impact on the soil microbial communities. While we observed that the irrigation regimes impacted both crop yield and disease prevalence, we found that neither this nor pathogen burden translate<u>d</u> to disruption in the dominant soil microbial communities, since the most abundant members of the microbial communities were stable irrespective of irrigation regime or initial pathogen burden of the seed potato stocks. However, by also including analyses that allowed consideration of changes in less abundant taxa, we observed that the initially high *Pectobacterium* pathogen burden on the JellyHigh seed stock was correlated with an increased abundance of particular microbial groups, even though the final disease burdens were similar between the low and high Jelly stocks. In contrast, we did not observe substantial changes in rarer taxa in comparisons between the JellyLow or EstimaZero stocks or in relation to irrigation regimes. This suggests that the conditions under which seed stocks are maintained and which lead to a high pathogen burden are more of an important driver of microbial community composition than the irrigation regime. Our results also emphasise the value of basing interpretation of microbiome community dynamics not only on the effects of treatments on the “core microbiome” but also more fine-scale analyses of rarer taxa, as suggested in a meta-analysis of agricultural studies in China by Jiao et al. (2019).

### Crop Yields

Crop yields were highly influenced by the irrigation regime. Not irrigating reduced yields for all planted stocks, and this was most pronounced for the EstimaZero mini-tubers, indicating that the rainfall for the region (Cambridge, UK) was insufficient to fulfil the growth requirements of the crop, as is often the case, given the need for irrigation management as reported by UK growers. However, irrigating when the soil moisture deficit (SMD) reached 40 mm increased the yields for all stocks. Potatoes are vulnerable to drought stress, as the rooting system is relatively shallow (Yamaguchi and Tanaka, 1990) but there is variation in water requirements for different potato varieties (Djaman et al., 2021; Xing et al., 2022). In addition, the method of irrigation supply can impact crop yields (Onder et al., 2005; Sarker et al., 2019a). Like other studies (Larkin et al., 2011; Sarker et al., 2019) we noted an increase in crop yields with irrigation. However, we only analysed the conventional furrow irrigation methodology that would be the typical system of growers in England. Rainfed growth (unirrigated) is a practice that is used in Scotland for seed crops, which experiences 2-3 times more rainfall than England (Statista, 2023). This emphasises regional variations and the need for adaptable management practices, particularly in the face of future climactic changes (Haverkort and Verhagen, 2008). Potato production models simulating climate change (temperature, solar radiation, precipitation, and CO_2_ levels) on potato growth development and yield have shown that by 2055 yields will increase in some areas (e.g. Western Europe) and decrease in others (e.g. North America and Eastern Europe) but that overall, production will have declined globally by 2085 (Raymundo et al., 2018). Given that such models do not include impacts of biotic interactions (pests, pathogens and beneficial microbes) or management practices, the impacts of climate change could be even more dramatic. Our results emphasise the importance of matching management practices to local climatic conditions and carefully monitoring of impacts on the most serious threats to production.

### Disease Prevalence

Although we observed only very low levels of rotting in our study, we observed that blackleg symptoms increased to a similar level in seed stocks with an initial *Pectobacterium* species burden regardless of whether the pathogen level was low or high. Importantly, even low levels of *Pectobacterium* in the seed stock translated into symptoms in the field that were on par with stock carrying a much higher initial burden. Our results contrast with those of Bain et al. (1990) and Toth et al. (2003) who correlated the development of disease with *Pectobacterium* inoculum levels in seed tubers in both field trials with native levels and with artificial infections following *in vitro* inoculation of blackleg-causing species. However, while our results were based on multiple repetitions, they may have differed from those above due to only single stocks being used in our study and the levels of *P. atrosepticum* in both our high and low stocks are low as compared to both studies. de Werra et al. (2020) observed in field trials in Switzerland that the levels of *Dickeya* spp., *Pectobacterium wasabiae*, and *P. carotovorum* subsp. brasiliense (now classified as *P. brasiliense*) could be correlated to initial inoculum levels but did not see the same trend with *P. atrosepticum*. In our study, blackleg symptoms were worsened with applied irrigation regimes and were highest in the regime that maintained a soil moisture deficit of less than 15 mm during the common scab control period and continued irrigation through the rest of the growing season. This confirms the link between blackleg disease and soil moisture content (Pérombelon, 2002). In contrast, blackleg symptoms were very low in the stock without initial *Pectobacterium* contamination (EstimaZero). These results, therefore, reinforce the requirement for stringent seed certification for blackleg as even low levels of initial bacterial contamination greatly increased disease incidence.

Although common scab disease prevalence was under 5% in our study, we observed increased disease symptoms in the unirrigated (rainfed only) treatments compared to all irrigated treatments; this is in agreement with previous field-trial studies (Lapwood et al., 1973; Wilson et al., 2001). The irrigation regime specifically designed to reduce common scab (Irrigation 2) was effective for all the stocks, and the regimes that continued watering post tuber-initiation (Irrigation 1 and 3) also reduced common scab as well as increasing the total yield of tubers. This emphasises the potential trade-off in disease management practices since these irrigation treatments also resulted in higher blackleg incidence. Once again, local decisions about irrigation practices are predicted to have even more importance as climate change will increase the unpredictability of rainfall (Raymundo et al., 2018).

### Impact of Irrigation on the Soil Microbial Communities

It is encouraging that our results show that differences in irrigation regimes did not perturb the dominant species or overall diversity of microbial communities associated with potatoes. Many studies have assessed the impact of irrigation on potato crop production (Shock et al., 2007; Djaman et al., 2021; Xing et al., 2022) but so far there has been limited specific considerations about interactions between crop irrigation and soil microbial communities Mavrodi et al. (2018). Of those studies that have, they explore the impacts of irrigation to soil microbial communities, and do so with the aim of understanding recycled wastewater rather than in an agricultural setting (Entry et al., 2008; Ibekwe et al., 2018; Zolti et al., 2019; Dang et al., 2019). The GLLVM approach by Niku et al (2019a) allows computationally efficient analysis of the correlations between environmental variables and microbial taxa. Using this method, we observed that certain microbial groups were positively associated with the unirrigated treatments. These include *Streptomyces*, *Terrabacter*, *Gemmatimonas*, *Sphingomonas*, and *Massilia* genera. *Streptomyces*, *Sphingomonas* and *Gemmatimonas*, which are common taxa that have been observed previously in soil from potato field trials (Wright et al., 2022; Marković et al., 2022). Although amplicon sequencing alone did not allow resolution to species, *Streptomyces scabies* and other *Streptomyces* spp. are the causative agents of common scab. However, species of *Streptomyces* and those of *Terrabacter* have also been linked to disease suppressive soils for many bacterial and fungal plant diseases (Wei et al., 2019; Heinsch et al., 2019). *Massilia* spp. play a role in the hydrolysis of inorganic phosphates and are thought to be more abundant in soils with limited phosphate (Samaddar et al., 2019). Similarly, *Gemmatimonas* spp. are phosphate-solubilizing species which can make phosphate bioavailable to plants (Zhang et al., 2019; Lu et al., 2020). These results suggest more complex interactions between the microbial communities and other environmental parameters (such as nutrient content).

### Impact of Pathogen Burden on the Soil Microbial Communities

Interestingly, we observed that the initial seed stock pathogen burden impacted beta diversity (between treatments) of soil microbial communities and the differential abundance of key taxa. Using Bray-Curtis dissimilarities, the communities from the EstimaZero and JellyLow stocks were more similar than the JellyHigh stock. The differential heat tree and GLLVM analysis confirmed that a high *Pectobacterium* burden in the starting seed stock resulted in increased *Anaerolineae*, *Chthoniobacter*, *Planctomycetota*, *Tepidisphaerales*, *Pirellula*, and *Vicinamibacteria* species, at both 50% plant emergence and at harvest, compared to the other stocks. The JellyLow stocks also became more similar to the JellyHigh stocks at harvest, showing increased *Planctomycetota*, *Vicinibactiera* and *Anaerolinea* compared to the EstimaZero stocks. Buckley et al. (2006) found that *Planctomycetota* taxa could be correlated with soil management history and these taxa showed increased diversity with increased nitrate concentrations. *Vicinamibacteria* are characteristic species in loamy soil, confirming the soil type of our field conditions (Kandasamy et al., 2021). *Anaerolineae* are strictly anaerobic bacteria that are fermentative species capable of growth on a diverse range of substrates (Sekiguchi, 2003; Xia et al., 2016; Song et al., 2020) and have been associated previously with waterlogged agricultural fields (Gschwend et al., 2020). Our research provides evidence of a distinct soil response to the potato stock with a high pathogen burden that is stronger than the effect of irrigation.

The challenge in microbiome studies is moving beyond statistical correlations to a mechanistic understanding. Therefore, we used the ensemble quotient (EQO) analysis, which identified the minimal subset of taxa linked to blackleg or common scab disease prevalence values based on patterns of statistical variation. This method further revealed that in the ridge samples the *Anaerolinea* (SBR1031) showed a high relative proportion in the subset of taxa that were linked to increasing blackleg prevalence. The underlying cause for the link between the taxa and the planting of seed stock with a high *Pectobacterium* pathogen burden (as revealed by the differential and EQO analysis) is unclear. These taxa could merely be responding to environmental conditions that coincidentally favour blackleg disease but could also be involved in synergistic activities with *Pectobacterium*. An alternate hypothesis is the ‘cry-for-help’ model, where plants modulate their phytohormones to alter the composition of the rhizosphere microbiome to promote pathogen-suppressive microbes to help protect them from attack (Halim et al., 2006; Doornbos et al., 2012; Rolfe et al., 2019). Thus, taxa from the *Anaerolinea* may get recruited during infection. Evidence for this hypothesis could be the development of a similar signature in the JellyLow at harvest. The passing of altered microbial signatures to the next ‘seed’ generation has been shown previously (Kong et al., 2019). This theory may be possible given that the JellyHigh was a generation 3 stock and blackleg levels accumulate over time (Pérombelon, 1992). Why we observe a strong signature in the ridge microbiome as compared to the root microbiome remains to be ascertained but may relate to the larger build-up of *Pectobacterium* at the ridge site. Moreover, our study only considers bacterial populations and, given the symbiotic association of potatoes with fungi (mycorrhiza), we are almost certainly not be capturing the complexity of these interactions. Therefore, future research efforts should focus on untangling the interactions between potato pathogen burden, plant defence responses and the soil microbiome (for multiple trophic groups including fungi), and more specifically on investigating further the roles of genera such as *Streptomyces* and *Anaerolinea* e.g., in highly controlled pot experiments. Understanding the associations between genera and management practices such as irrigation (e.g. *Streptomyces*) or a high pathogen burden (e.g. *Anaerolinea*) could help to develop strategies to modulate the response of agricultural potato soil microbiomes.

## Acknowledgements

We would like to thank the field and sampling staff at NIAB, Cambridge, UK, for their contributions to the fieldwork. We thank Simon Alexander and Eric Anderson for useful feedback on potato production within the UK. In addition, we thank Julie Galbraith, David McGuinness and others at the Glasgow Polyomics sequencing facility for processing the sequencing libraries.

## Author Contributions

C.K. (Data curation: Equal; Methodology: Equal; Formal analysis: Equal; Investigation: Equal; Visualisation; Lead; Writing – original draft: Lead; Writing – review & editing: Lead).

E.K. (Investigation: Equal; Methodology: Equal; Data curation: Equal; Writing – review & editing: Supporting).

M.A.S. (Conceptualisation: Supporting; Formal analysis: Supporting; Methodology: supporting; Writing – review & editing: supporting).

C.F.N. (Resources: Supporting; Writing – review & editing: Supporting)

J.M. (Resources: Supporting; Conceptualisation: Supporting; Investigation; Supporting: Writing – reviewing and editing; Supporting)

S.H. (Conceptualisation: Supporting; Visualization: Supporting; Writing – review and editing: Supporting)

I.T. (Conceptualization: Supporting; Visualization: Supporting; Writing – review and editing: Supporting)

B.K.M. (Conceptualisation: Equal; Resources: Equal; Investigation: Equal; Formal analysis: Supporting; Writing – review & editing: Equal).

U.Z.I. (Conceptualisation: Equal; Resources: Equal; Software: Lead; Investigation: Equal; Formal analysis: Equal; Writing – review & editing: Equal).

