## Supplementary Information for "Balancing the scales: Impact of irrigation and pathogen burden on potato blackleg disease and soil microbial communities"

**Supplementary Methods**

*Soil Profile and History*

The soil profile was approximately 74 % sand, 17 % silt, 9 % clay with 8-20 % stone, 2.7 % organic matter content and a pH of 7.4. The phosphate (P), potassium (K) and magnesium (Mg) indices were 2+, 2+ and 2, respectively. The field was subsoiled on 8 September 2019 and drilled with a cover crop of winter oats on 11 September. Fertilizer (101 kg/ha $P_2O_5$ and 102 kg/ha $K_2O$) was broadcast on the cover crop on 23-24 March. The cover crop was sprayed with glyphosate on 26 March 2020 and mowed to a height of 10-12 cm using a tractor-mounted mower on 27 March. The area was cultivated twice on 30 and 31 March using a Kuhn Cultimer and roto-ridged on 3 April with a Rumptstad rototiller.

*Statistical Analyses*

<u>Crop Yield and Crop Health Statistical Analyses</u>

Variates were analysed by analysis of variance using the GenStat® Release 16.1 statistical package. Treatment means were significantly different only if the probability of differences occurring by chance were less than 5 % ($P < 0.05$).

<u>Microbial Statistical Analyses</u>

The microbial statistical analyses described below were performed in R (R version 4.1.3: R Core Team, 2022) using 'phyloseq' (McMurdie and Holmes, 2013), the biom files from above, and in some cases, the functional .tsv files, and the metadata associated with the field trials (Supplementary Data File 1).

<u>Taxonomic and Functional Diversity</u>

Our rarefaction curves indicate that sampling depth was sufficient to estimate true microbial diversity of samples (Supplementary Figure 2). Taxonomic diversity analyses were conducted using the vegan package in R (Oksanen et al., 2020). For alpha-diversity (differences within treatments) we rarified the samples to minimum library size (5,154 reads) and considered richness and Pielou's evenness measures based on the number of unique ASVs and balance in the distribution of ASV abundances, respectively. Beta diversity (differences between communities) was visualised using principal coordinate analysis (PCoA) of ASVs using Bray-Curtis distance (Bray and Curtis, 1957). Analysis of variance was performed using Vegan's Adonis() against the Bray-Curtis dissimilarity results (PERMANOVA) using the variables of 'Time', 'Sample Type', 'Irrigation Type', 'Plot Number' and 'Plot Area' (based on plot locations) and 'Pathogen burden'.

To select the variables most strongly associated with the variance of the observed communities, redundancy analysis (RDA) with forward selection (based on 999 permutations) was applied to filter out environmental variables (including irrigation volume, blackleg symptoms per plot, the weight of rotten tubers and scab symptoms per plot; variables retained at $p < 0.05$) before using PERMANOVA (Vass et al., 2020).

Functional profiles generated from the PiCrust2 analysis were used to calculate functional diversity in the same approach as above. However, since there is redundancy in KEGG orthologs (i.e. a single KO can be part of multiple pathways), using the KEGG ortholog table to calculate beta-diversity may not be done efficiently with traditional measures such as Bray-Curtis distance. To circumvent this issue, a new beta-diversity measure called hierarchical meta-storms (Zhang et al., 2021) was used, which assumes the hierarchy of pathways in terms of KOs and considers a weighted approach by propagating the abundance of KOs higher up the hierarchy to the actual pathways these KOs belong to. This was implemented using the 'hrms' and 'ape' packages in R (Paradis et al., 2023; Zhang et al., 2021). For alpha-diversity functional analysis, 'Evenness' refers to the balance in the distribution of KEGG ortholog (KOs) abundance and 'Richness' refers to the number of unique KOs.

<u>Contribution of Rare Microbial Taxa to Taxonomic Diversity</u>

One of the challenges with understanding dynamics in soil communities is the high diversity of the samples and the cumulative impact (or response) of less abundant or 'rare' taxa as compared to abundant species. Therefore, we implemented analysis and code from Yang et al. (2017) to quantify rare taxa and specifically investigate whether there are specific taxa that potentially increase in abundance under specific environmental or treatment conditions (conditionally rare taxa). Broadly, ASVs were categorised as either 'Abundant' (ASVs having an average relative abundance above 1% across all samples) or 'Rare' (ASVs having an average relative abundance below 1% across all samples). Rare taxa fell into one of three categories - A) Conditionally Rare – ASVs with 100-fold difference between their minimum and maximum relative abundance (max:min>100); B) Persistently Rare – ASVs with a maximum relative abundance < 5 times their minimum value (max:min<=5), and C) Other Rare – Rare ASVs that fall out with the range of B (i.e. an ASV with a maximum relative abundance that is > 5 times their minimum value but does not have 100-fold difference between the minimum and maximum value). We then estimated how much 'Abundant' or 'Rare' (Conditionally Rare, Persistently Rare and Other Rare) ASVs contribute to Bray-Curtis dissimilarity in our samples as compared to the total microbial community, using a modification of the custom R script from Yang et al. (2017).

### Differential Microbial Genera Associated with Irrigation Regime, Potato Stock and Time Sampled

Differential heat tree analysis was performed on proportionally normalised microbial community data to identify taxonomic clades that are differentially expressed between groups. The groups tested were as follows: i) irrigation regime (Unirrigated, Irrigation 1, Irrigation 2 and Irrigation 3); ii) potato stock/pathogen burden (Zero, Low and High); and iii) time sampled (Plant emergence [T_E] and Harvest [T_H]). The complete methodology is outlined in McKenna et al. (2020) and uses the Metacoder package (Foster et al., 2017). The differential heat tree analysis provides better visual cues than the traditional analyses by virtue of showing the taxonomic lineages, it is however, less robust. This is primarily because the differential analysis within Metacoder is based on Wilcoxon Rank-Sum test, and while this is useful for hypothesis testing, the statistical robustness of the Wilcoxon Rank-Sum test has been questioned (Simmel, 1958; Whitley and Ball, 2002). As an alternative approach to finding individual microbial genera associated with experimental treatments, we applied a generalised linear latent variable model (GLLVM) to the dataset (Niku et al. 2019a, 2019b) where the abundance of a specific taxon is fitted with a distribution, typically negative binomial. This method extends a basic generalised linear model that fits a regression line of the mean taxa abundances of individual taxa (we have considered the top 50 most abundant microbial genera) against the experimental sources of variation, and by incorporating latent variables. The coefficients associated with the latent variables were not considered in our study. We finally obtain microbe-specific beta-coefficient values associated with an individual experimental variable (along with 95% confidence intervals using a bootstrapping approach). The range of the interval of these coefficients with values greater than 0 indicates a positive association of that variable with the abundance of that taxon, whilst a value less than 0 implies a negative association. Where the 95% confidence interval crosses the 0 boundary, that variable is determined to not affect taxon abundance (insignificant relationship). Further details are described in Niku et al. (2019a, 2019b).

Microbial Taxa Associated with Blackleg and Common Scab Disease Prevalence

We used the Ensemble Quotient Optimisation (EQO) method of Shan et al. (2023) to find the subset of microbial genera (called an ensemble) associated with blackleg disease symptoms or common scab disease symptoms at potato harvest. Briefly, the method creates a vector of 0s and 1s as entries (called $\boldsymbol{x}$, with each entry specific to a single taxon to suggest if it is part of the ensemble or not). This ensemble is recovered in the context of a continuous variable, $\boldsymbol{y}$ (e.g. percentage blackleg or common scab symptoms), by optimizing an *Ensemble Quotient* $EQ = \frac{\boldsymbol{x}^T\boldsymbol{Q}\boldsymbol{x}}{\boldsymbol{x}^T\boldsymbol{P}\boldsymbol{x}}$, through a genetic algorithm, where $\boldsymbol{P}$ and $\boldsymbol{Q}$ are algebraic transformations of the community matrix that captures the covariance between ASVs, and the covariance between ASVs and $\boldsymbol{y}$. Once optimised, we obtain the ensemble whose collated relative abundance correlates highly with the continuous variable $\boldsymbol{y}$.

In the genetic algorithm to optimize the EQ to obtain $\boldsymbol{x}$, we used the following parameterizations: a population size of 100 solutions, using maximum of 1,200 generations, and a maximum 20 taxa to be returned as an ensemble; further details can be found at https://github.com/Xiaoyu2425/Ensemble-Quotient-Optimization.

**Supplementary Figures and Tables**

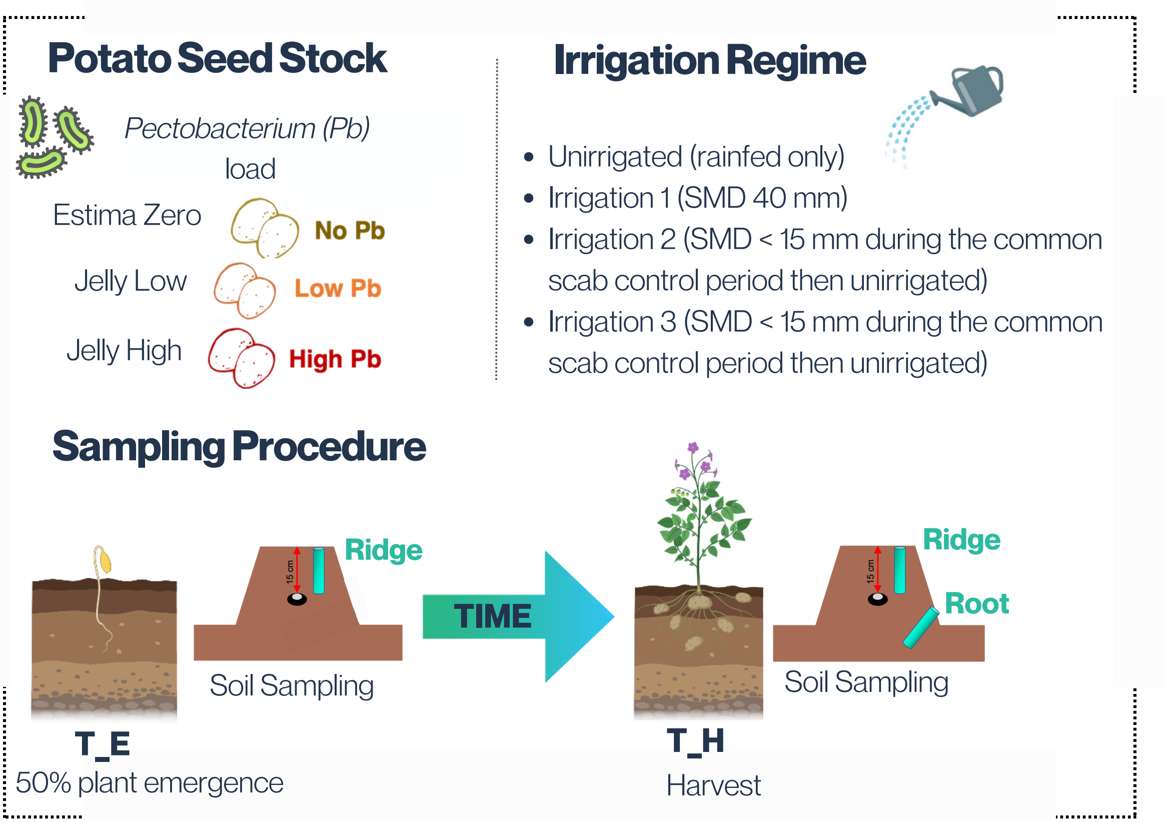

**Supplementary Figure 1.** Schematic diagram showing the experimental design of this study. This includes the choice of potato seed stock with high, low (both Jelly varieties) and zero (Estima mini-tubers) starting levels of *Pectobacterium* species and irrigation regimes (Unirrigated [rainfed only], Irrigation 1 [SMD 40 mm], Irrigation 2 [SMD < 15 mm during the common scab control period then unirrigated], and Irrigation 3 [SMD < 15 mm during the common scab control period then unirrigated]. Comparisons in microbial communities across time were made using soil sampled from the ridge (near the top of the furrow) at 50% plant emergence (T_E Ridge) and harvest (T_H Ridge). In addition, to determine if there were differences in microbial communities sampled closer to the plant roots, samples were also taken at the bottom of the furrow at harvest only (T_H Root), to avoid damaging roots in the emerging plants. Details about the plot layout can be found in Supplementary Data File 2.

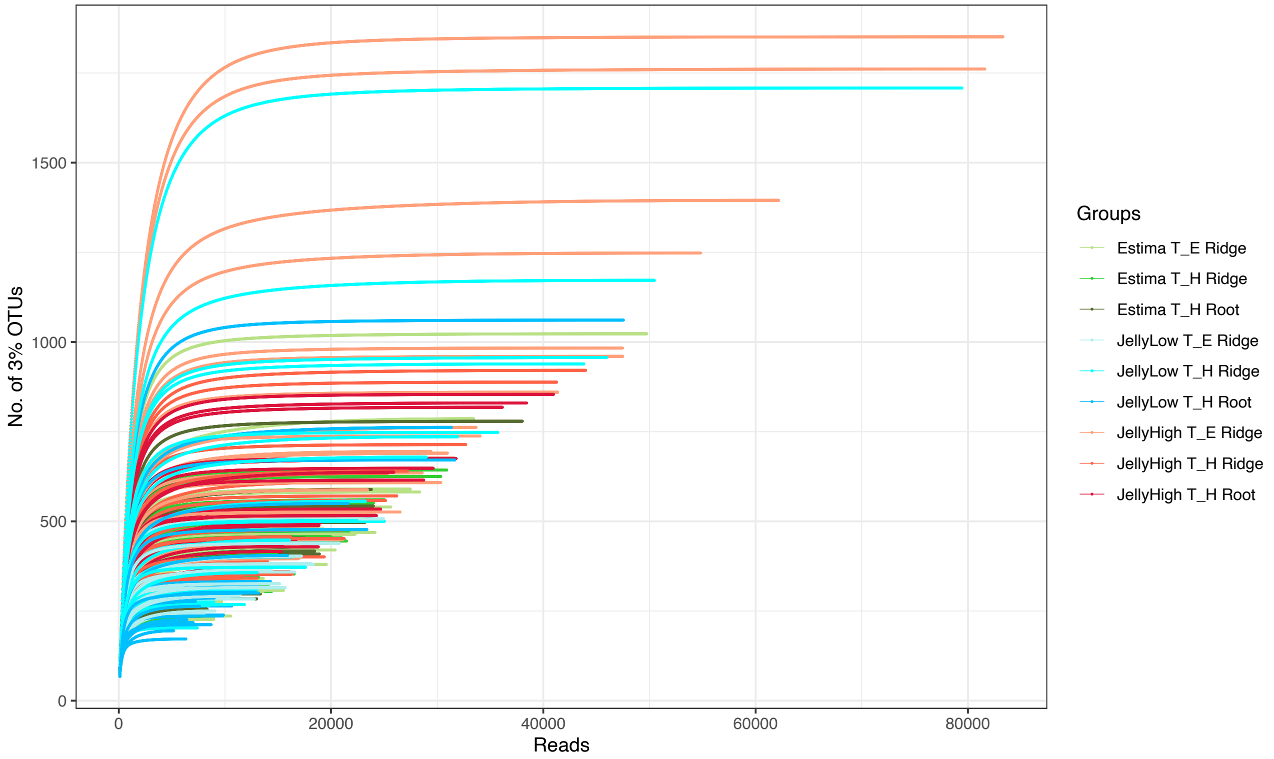


**Supplementary Figure 2.** Rarefaction curves showing the number of reads from the 16S rRNA gene in DNA from the soil samples on the x-axis and the number of OTUs within a 97% percent sequence similarity threshold on the y-axis. Treatment groups are indicated by colour-coded lines.

**Supplementary Table 1.** Percentage of blackleg disease symptoms in potato plants per plot over the course of the experimental field trial in 2020.

| Potato Stock | Irrigation | Plot Number | 18-Jun | 26-Jun | 06-Jul | 31-Jul | 20-Aug |
|---|---|---|---|---|---|---|---|
| **JellyLow** | **Irrigation 1** | 1_1 | 0.000 | 8.333 | 25.000 | 27.083 | 27.083 |
| **JellyLow** | **Irrigation 2** | 1_2 | 0.000 | 12.500 | 29.167 | 31.250 | 33.333 |
| **JellyLow** | **Irrigation 3** | 1_3 | 4.167 | 12.500 | 29.167 | 31.250 | 35.417 |
| **EstimaZero** | **Unirrigated** | 1_4 | 0.000 | 0.000 | 0.000 | 0.000 | 0.000 |
| **JellyHigh** | **Unirrigated** | 1_5 | 0.000 | 6.250 | 18.750 | 16.667 | 14.583 |
| **JellyHigh** | **Irrigation 3** | 1_6 | 0.000 | 10.417 | 22.917 | 22.917 | 33.333 |
| **EstimaZero** | **Irrigation 2** | 1_7 | 0.000 | 0.000 | 4.167 | 4.167 | 6.250 |

| | | | | | | | |
|---|---|---|---|---|---|---|---|
| **EstimaZero** | **Irrigation 1** | 1_8 | 0.000 | 0.000 | 4.167 | 4.167 | 4.167 |
| **EstimaZero** | **Irrigation 3** | 1_9 | 0.000 | 0.000 | 2.083 | 2.083 | 4.167 |
| **JellyLow** | **Unirrigated** | 1_10 | 4.167 | 8.333 | 16.667 | 25.000 | 22.917 |
| **JellyHigh** | **Irrigation 1** | 1_11 | 2.083 | 8.333 | 12.500 | 25.000 | 27.083 |
| **JellyHigh** | **Irrigation 2** | 1_12 | 0.000 | 20.833 | 33.333 | 37.500 | 39.583 |
| **JellyHigh** | **Irrigation 1** | 2_1 | 0.000 | 12.500 | 20.833 | 20.833 | 22.917 |
| **JellyHigh** | **Unirrigated** | 2_2 | 0.000 | 8.333 | 12.500 | 18.750 | 18.750 |
| **EstimaZero** | **Irrigation 3** | 2_3 | 0.000 | 0.000 | 0.000 | 0.000 | 2.083 |
| **JellyLow** | **Irrigation 1** | 2_4 | 4.167 | 12.500 | 27.083 | 29.167 | 29.167 |
| **JellyLow** | **Irrigation 2** | 2_5 | 4.167 | 12.500 | 22.917 | 31.250 | 31.250 |
| **JellyLow** | **Irrigation 3** | 2_6 | 0.000 | 10.417 | 18.750 | 22.917 | 29.167 |
| **EstimaZero** | **Irrigation 1** | 2_7 | 0.000 | 0.000 | 0.000 | 0.000 | 0.000 |
| **JellyHigh** | **Irrigation 2** | 2_8 | 0.000 | 8.333 | 14.583 | 20.833 | 25.000 |
| **JellyHigh** | **Irrigation 3** | 2_9 | 0.000 | 8.333 | 12.500 | 22.917 | 31.250 |
| **JellyLow** | **Unirrigated** | 2_10 | 4.167 | 6.250 | 10.417 | 12.500 | 12.500 |
| **EstimaZero** | **Unirrigated** | 2_11 | 0.000 | 0.000 | 0.000 | 0.000 | 0.000 |
| **EstimaZero** | **Irrigation 2** | 2_12 | 0.000 | 0.000 | 0.000 | 2.083 | 2.083 |
| **JellyLow** | **Irrigation 1** | 3_1 | 4.167 | 20.833 | 25.000 | 31.250 | 33.333 |
| **JellyHigh** | **Unirrigated** | 3_2 | 2.083 | 6.250 | 8.333 | 14.583 | 16.667 |
| **JellyLow** | **Irrigation 2** | 3_3 | 0.000 | 14.583 | 20.833 | 18.750 | 25.000 |
| **EstimaZero** | **Unirrigated** | 3_4 | 0.000 | 0.000 | 2.083 | 2.083 | 2.083 |
| **JellyHigh** | **Irrigation 2** | 3_5 | 0.000 | 6.250 | 12.500 | 18.750 | 22.917 |
| **JellyLow** | **Irrigation 3** | 3_6 | 0.000 | 25.000 | 29.167 | 33.333 | 37.500 |
| **JellyLow** | **Unirrigated** | 3_7 | 2.083 | 6.250 | 10.417 | 18.750 | 18.750 |
| **EstimaZero** | **Irrigation 1** | 3_8 | 0.000 | 0.000 | 4.167 | 4.167 | 4.167 |
| **EstimaZero** | **Irrigation 3** | 3_9 | 0.000 | 0.000 | 2.083 | 2.083 | 6.250 |
| **JellyHigh** | **Irrigation 1** | 3_10 | 0.000 | 4.167 | 16.667 | 10.417 | 16.667 |
| **EstimaZero** | **Irrigation 2** | 3_11 | 0.000 | 0.000 | 4.167 | 8.333 | 8.333 |
| **JellyHigh** | **Irrigation 3** | 3_12 | 2.083 | 12.500 | 20.833 | 33.333 | 37.500 |

**Supplementary Table 2.** Proportion of tubers with rotting (%) and weight of rotted tubers (tubers/hectare) at potato harvest. Table also shows the results of the statistical analysis of individual stocks and irrigation regimes. The respective degrees of freedom (D.F.) is given within the standard error (S.E.).

| Potato Stock | Irrigation Treatment | Proportion of tubers with rotting (%) | Weight of rotted tubers (t/ha) |
|---|---|---|---|
| **Jelly High** | **Unirrigated** | 0.00 | 0.00 |
| **Jelly High** | **Irrigation 3** | 1.37 | 1.23 |
| **Jelly High** | **Irrigation 2** | 0.00 | 0.00 |
| **Jelly High** | **Irrigation 1** | 0.43 | 0.03 |
| **Jelly Low** | **Unirrigated** | 0.00 | 0.00 |
| **Jelly Low** | **Irrigation 3** | 0.00 | 0.00 |
| **Jelly Low** | **Irrigation 2** | 0.43 | 0.01 |
| **Jelly Low** | **Irrigation 1** | 1.07 | 0.57 |
| **Estima Zero** | **Unirrigated** | 0.53 | 0.13 |
| **Estima Zero** | **Irrigation 3** | 1.17 | 0.77 |

| | | | |
|---|---|---|---|
| **Estima Zero** | **Irrigation 2** | 0.73 | 0.30 |
| **Estima Zero** | **Irrigation 1** | 0.40 | 0.13 |
| S.E. (22 D.F.) | | 0.470 | 0.391 |
| Jelly High | | 0.45 | 0.32 |
| Jelly Low | | 0.38 | 0.14 |
| Estima | | 0.71 | 0.33 |
| S.E. (22 D.F.) | | 0.235 | 0.196 |
| Unirrigated | | 0.18 | 0.04 |
| Irrigated | | 0.84 | 0.67 |
| Scab only | | 0.39 | 0.10 |
| Processing | | 0.63 | 0.24 |
| S.E. (22 D.F.) | | 0.272 | 0.226 |
| Fprob Stock | | 0.583 | 0.748 |
| Fprob Irrigation | | 0.355 | 0.230 |
| Fprob Stock*Irrigation | | 0.345 | 0.451 |

**Supplementary Table 3.** Genus identifier list for Supplementary Figure 2 and the average percentage prevalence in all samples.

| Genus Identifier | Taxonomy |
|---|---|
| | *Archaea;Crenarchaeota;Nitrososphaeria;Nitrososphaerales;Nitrososphaeraceae;Nitrososphaeraceae* |
| | *Bacteria;Acidobacteriota;Blastocatellia;Pyrinomonadales;Pyrinomonadaceae* |
| | *Bacteria;Acidobacteriota;Vicinamibacteria;Vicinamibacterales;uncultured;uncultured* |
| | *Bacteria;Acidobacteriota;Vicinamibacteria;Vicinamibacterales;Vicinamibacteraceae;Vicinamibacteraceae* |
| | *Bacteria;Actinobacteriota;Actinobacteria;Frankiales;Geodermatophilaceae* |
| | *Bacteria;Actinobacteriota;Actinobacteria;Micrococcales;Intrasporangiaceae* |
| | *Bacteria;Actinobacteriota;Actinobacteria;Micrococcales;Micrococcaceae* |
| | *Bacteria;Actinobacteriota;Actinobacteria;Propionibacteriales;Nocardioidaceae* |
| | *Bacteria;Actinobacteriota;MB−A2−108* |
| | *Bacteria;Actinobacteriota;Thermoleophilia;Gaiellales;Gaiellaceae* |
| | *Bacteria;Actinobacteriota;Thermoleophilia;Solirubrobacterales;67−14;67−14* |
| | *Bacteria;Verrucomicrobiota;Verrucomicrobiae;Chthoniobacterales;Chthoniobacteraceae* |
| | *Bacteria;Chloroflexi;Chloroflexia;Thermomicrobiales;JG30−KF−CM45;JG30−KF−CM45* |
| | *Bacteria;Chloroflexi;KD4−96;KD4−96;KD4−96;KD4−9* |
| | *Bacteria;Firmicutes;Bacilli;Bacillales;Bacillaceae* |

| | |
|---|---|
| | *Bacteria;Gemmatimonadota;Gemmatimonadetes;Gemmatimonadales;Gemmatimonadaceae* |
| | *Bacteria;Methylomirabilota;Methylomirabilia;Rokubacteriales;Rokubacteriales;Rokubacteriales* |
| | *Bacteria;Proteobacteria;Alphaproteobacteria;Rhizobiales;Methyloligellaceae* |
| | *Bacteria;Proteobacteria;Alphaproteobacteria;Rhizobiales;Xanthobacteraceae* |
| | *Bacteria;Proteobacteria;Alphaproteobacteria;Sphingomonadales; Sphingomonadaceae* |
| | *Bacteria;Proteobacteria;Gammaproteobacteria;Burkholderiales; Comamonadaceae* |
| | *Bacteria;Proteobacteria;Gammaproteobacteria;Burkholderiales; SC−I−84* |
| | *Bacteria;Proteobacteria;Gammaproteobacteria;Steroidobacterales;Steroidobacteraceae* |
| | *Bacteria;Proteobacteria;Alphaproteobacteria;Rhizobiales;Xanthobacteraceae* |
| | *Bacteria;Verrucomicrobiota;Verrucomicrobiae;Chthoniobacterales; Chthoniobacteraceae* |

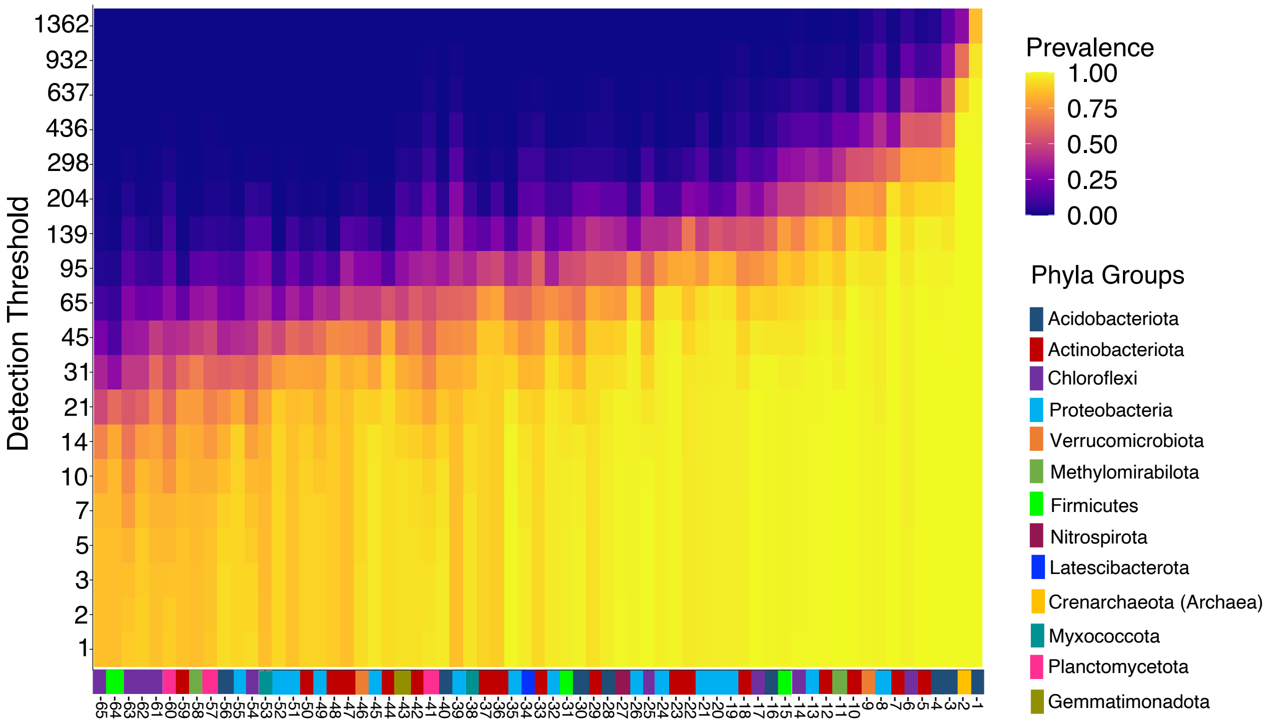


**Supplementary Figure 3. The 'core microbiome of the soil microbial communities from the field trial.** The 'core microbiome' was set at 85% minimum prevalence of samples. Numbers referring to the individual genera are shown on the x-axis (defined in Supplementary Table 3), with colours highlighting the taxonomic phyla. The y-axis shows the detection threshold of absolute genera abundances and heatmap colours showing the average prevalence of each genus within all samples, ranging from 0-1.

**Supplementary Table 4.** Genus identifier list for Supplementary Figure 2 and the average percentage prevalence in all samples.

| Genus Identifier | Taxonomy |
|---|---|
| **1** | *Bacteria;Acidobacteriota;Vicinamibacteria;Vicinamibacterales;uncultured;uncultured* |
| **2** | *Archaea;Crenarchaeota;Nitrososphaeria;Nitrososphaerales;Nitrososphaeraceae;Nitrososphaeraceae* |
| **3** | *Bacteria;Acidobacteriota;Vicinamibacteria;Vicinamibacterales;Vicinamibacteraceae;Vicinamibacteraceae* |
| **4** | *Bacteria;Acidobacteriota;Blastocatellia;Pyrinomonadales;Pyrinomonadaceae;RB41* |
| **5** | *Bacteria;Actinobacteriota;Actinobacteria;Propionibacteriales;Nocardioidaceae;Nocardioides* |
| **6** | *Bacteria;Chloroflexi;KD4−96;KD4−96;KD4−96;KD4−9* |
| **7** | *Bacteria;Actinobacteriota;Thermoleophilia;Gaiellales;Gaiellaceae; Gaiella* |
| **8** | *Bacteria;Proteobacteria;Alphaproteobacteria;Sphingomonadales;Sphingomonadaceae;Sphingomonas* |
| **9** | *Bacteria;Verrucomicrobiota;Verrucomicrobiae;Chthoniobacterales; Chthoniobacteraceae;Candidatus_Udaeobacter* |
| **10** | *Bacteria;Actinobacteriota;Thermoleophilia;Solirubrobacterales;67−14;67−14* |
| **11** | *Bacteria;Methylomirabilota;Methylomirabilia;Rokubacteriales;Rokubacteriales;Rokubacteriales* |
| **12** | *Bacteria;Actinobacteriota;MB−A2−108;MB−A2−108;MB−A2−108;MB−A2−108* |
| **13** | *Bacteria;Proteobacteria;Gammaproteobacteria;Burkholderiales;SC−I−84;SC−I−84* |
| **14** | *Bacteria;Chloroflexi;Chloroflexia;Thermomicrobiales;JG30−KF−CM45;JG30−KF−CM45* |
| **15** | *Bacteria;Firmicutes;Bacilli;Bacillales;Bacillaceae;Bacillus* |
| **16** | *Bacteria;Acidobacteriota;Holophagae;Subgroup_7;Subgroup_7;Subgroup_7* |
| **17** | *Bacteria;Chloroflexi;Gitt−GS−136;Gitt−GS−136;Gitt−GS−136;Gitt−GS−136* |
| **18** | *Bacteria;Actinobacteriota;Actinobacteria;Streptomycetales;Streptomycetaceae;Streptomyces* |
| **19** | *Bacteria;Proteobacteria;Gammaproteobacteria;Xanthomonadales;Xanthomonadaceae;Arenimonas* |
| **20** | *Bacteria;Proteobacteria;Alphaproteobacteria;Rhizobiales;Xanthobacteraceae;Bradyrhizobium* |
| **21** | *Bacteria;Proteobacteria;Gammaproteobacteria;Pseudomonadales;Pseudomonadaceae;Pseudomonas* |
| **22** | *Bacteria;Actinobacteriota;Actinobacteria;Corynebacteriales;Mycobacteriaceae;Mycobacterium* |
| **23** | *Bacteria;Actinobacteriota;Actinobacteria;Frankiales;Geodermatophilaceae;Blastococcus* |
| **24** | *Bacteria;Proteobacteria;Alphaproteobacteria;Rhizobiales;Xanthobacteraceae;Pseudolabrys* |

| | |
|---|---|
| **25** | *Bacteria;Chloroflexi;TK10;TK10;TK10;TK10* |
| **26** | *Bacteria;Proteobacteria;Alphaproteobacteria;Rhizobiales;Beijerinckiaceae;Microvirga* |
| **27** | *Bacteria;Nitrospirota;Nitrospiria;Nitrospirales;Nitrospiraceae;Nitrospira* |
| **28** | *Bacteria;Acidobacteriota;Blastocatellia;11−24;11−24;11−24* |
| **29** | *Bacteria;Actinobacteriota;Thermoleophilia;Solirubrobacterales;Solirubrobacteraceae;Solirubrobacter* |
| **30** | *Bacteria;Acidobacteriota;Vicinamibacteria;Subgroup_17;Subgroup_17;Subgroup_17* |
| **31** | *Bacteria;Firmicutes;Bacilli;Paenibacillales;Paenibacillaceae;Paenibacillus* |
| **32** | *Bacteria;Proteobacteria;Gammaproteobacteria;Gammaproteobacteria_Incertae_Sedis;Unknown_Family;Acidibacter* |
| **33** | *Bacteria;Actinobacteriota;Actinobacteria;Propionibacteriales;Propionibacteriaceae;Microlunatus* |
| **34** | *Bacteria;Latescibacterota;Latescibacterota;Latescibacterota;Latescibacterota;Latescibacterota* |
| **35** | *Bacteria;Proteobacteria;Gammaproteobacteria;PLTA13;PLTA13;PLTA13* |
| **36** | *Bacteria;Actinobacteriota;Actinobacteria;Micrococcales;Intrasporangiaceae;Terrabacter* |
| **37** | *Bacteria;Actinobacteriota;Actinobacteria;Micrococcales;Microbacteriaceae;Agromyces* |
| **38** | *Bacteria;Myxococcota;bacteriap25;bacteriap25;bacteriap25;bacteriap25* |
| **39** | *Bacteria;Proteobacteria;Gammaproteobacteria;Burkholderiales;Oxalobacteraceae;Massilia* |
| **40** | *Bacteria;Acidobacteriota;Thermoanaerobaculia;Thermoanaerobaculales;Thermoanaerobaculaceae;Subgroup_10* |
| **41** | *Bacteria;Planctomycetota;Phycisphaerae;Tepidisphaerales;WD2101_soil_group;WD2101_soil_group* |
| **42** | *Bacteria;Actinobacteriota;Acidimicrobiia;IMCC26256;IMCC26256;IMCC26256* |
| **43** | *Bacteria;Gemmatimonadota;Gemmatimonadetes;Gemmatimonadales;Gemmatimonadaceae;Gemmatimonas* |
| **44** | *Bacteria;Actinobacteriota;Actinobacteria;Micrococcales;Cellulomonadaceae;Cellulomonas* |
| **45** | *Bacteria;Proteobacteria;Alphaproteobacteria;Reyranellales;Reyranellaceae;Reyranella* |
| **46** | *Bacteria;Verrucomicrobiota;Verrucomicrobiae;Chthoniobacterales;Xiphinematobacteraceae;Candidatus_Xiphinematobacter* |
| **47** | *Bacteria;Actinobacteriota;Actinobacteria;Pseudonocardiales;Pseudonocardiaceae;Pseudonocardia* |
| **48** | *Bacteria;Actinobacteriota;Actinobacteria;Streptosporangiales;Streptosporangiaceae;Streptosporangium* |
| **49** | *Bacteria;Proteobacteria;Alphaproteobacteria;Tistrellales;Geminicoccaceae;Candidatus_Alysiosphaera* |
| **50** | *Bacteria;Actinobacteriota;Acidimicrobiia;Microtrichales;Ilumatobacteraceae;CL500−29_marine_group* |

| | |
|---|---|
| **51** | *Bacteria;Proteobacteria;Gammaproteobacteria;Burkholderiales;TRA3−20;TRA3−20* |
| **52** | *Bacteria;Proteobacteria;Alphaproteobacteria;Dongiales;Dongiaceae;Dongia* |
| **53** | *Bacteria;Myxococcota;Polyangia;Haliangiales;Haliangiaceae;Haliangium* |
| **54** | *Bacteria;Chloroflexi;Anaerolineae;SBR1031;A4b;A4b* |
| **55** | *Bacteria;Proteobacteria;Alphaproteobacteria;Rhizobiales;Rhizobiales_Incertae_Sedis;Nordella* |
| **56** | *Bacteria;Acidobacteriota;Subgroup_25;Subgroup_25;Subgroup_25;Subgroup_25* |
| **57** | *Bacteria;Planctomycetota;Planctomycetes;Pirellulales;Pirellulaceae;Pirellula* |
| **58** | *Bacteria;Methylomirabilota;Methylomirabilia;Rokubacteriales;WX65;WX65* |
| **59** | *Bacteria;Actinobacteriota;Rubrobacteria;Rubrobacterales;Rubrobacteriaceae;Rubrobacter* |
| **60** | *Bacteria;Planctomycetota;OM190;OM190;OM190;OM190* |
| **61** | *Bacteria;Chloroflexi;Dehalococcoidia;S085;S085;S085* |
| **62** | *Bacteria;Chloroflexi;Anaerolineae;RBG−13−54−9;RBG−13−54−9;RBG−13−54−9* |
| **63** | *Bacteria;Chloroflexi;Anaerolineae;SBR1031;SBR1031;SBR1031* |
| **64** | *Bacteria;Firmicutes;Bacilli;Alicyclobacillales;Alicyclobacillaceae;Tumebacillus* |
| **65** | *Bacteria;Chloroflexi;JG30−KF−CM66;JG30−KF−CM66;JG30−KF−CM66;JG30−KF−CM66* |

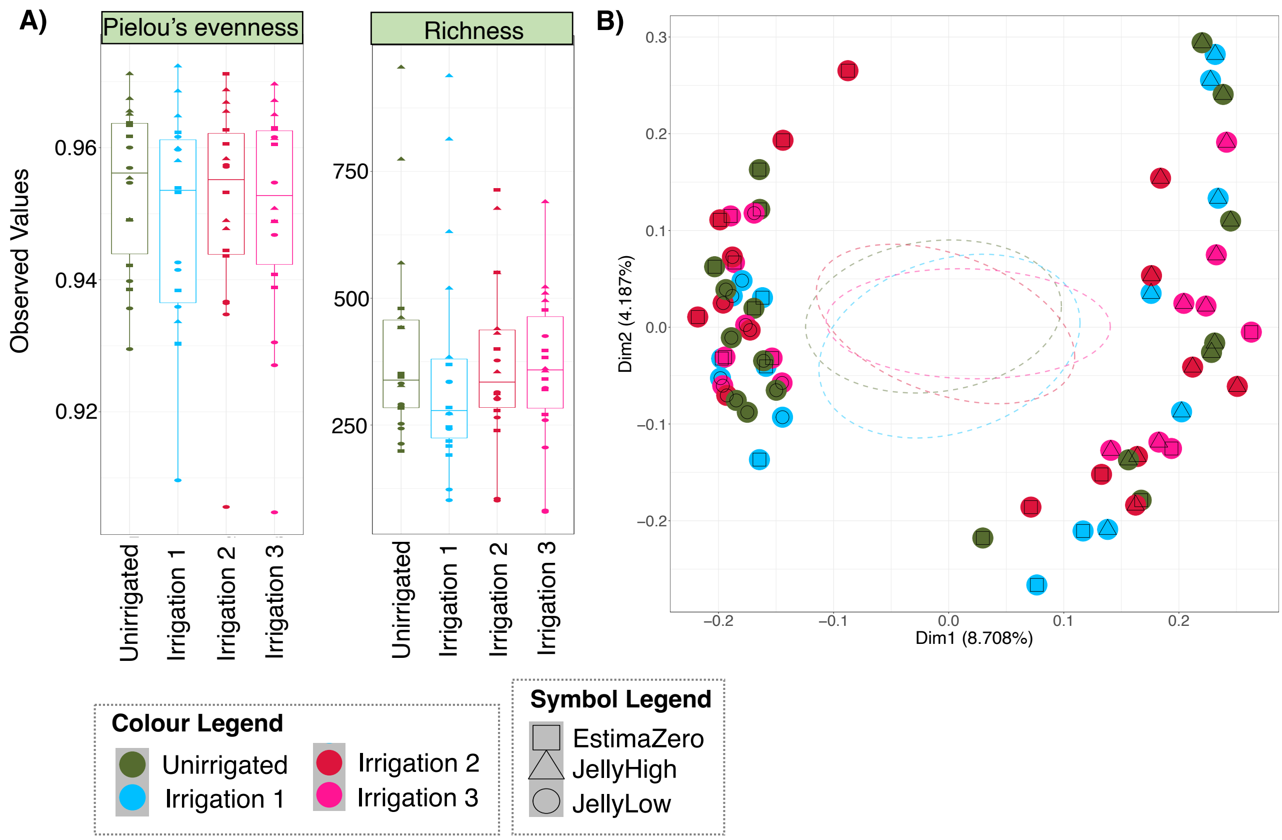


**Supplementary Figure 4. Taxonomic diversity of the soil microbiomes (ridge samples) at the 50% plant emergence (T_E) timepoint.** Colours represent the irrigation regime (Unirrigated, Irrigation 1, Irrigation 2, and Irrigation 3). Symbols represent the potato seed stock (EstimaZero, JellyLow, and JellyHigh). **A)** Shows alpha diversity based on the relative distribution of taxa (Pielou's evenness) and the number of different ASVs (Richness) within the communities; there were no significant differences between the watering regimes and no obvious clustering based on potato stock. **B)** Principal coordinate analysis (PCoA) based on Bray-Curtis distance of beta diversity dissimilarity between the communities; note that there was no distinct clustering in relation to irrigation regime, but there was clustering with respect to potato stock.

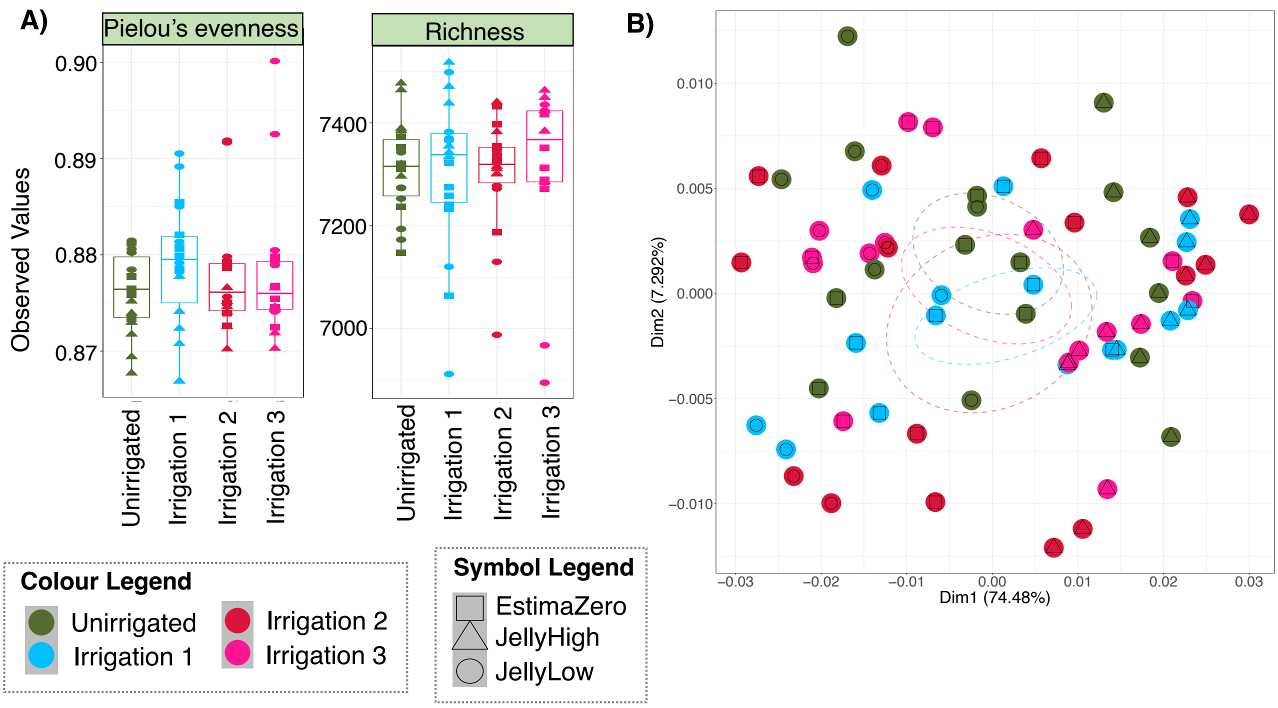


**Supplementary Figure 5. Functional diversity of the soil microbiomes (ridge samples) at the 50% plant emergence (T_E) timepoint.** Colours represent the irrigation regime (Unirrigated, Irrigation 1, Irrigation 2, and Irrigation 3). Symbols represent the potato seed stock (EstimaZero, JellyLow, and JellyHigh). **A)** Shows diversity based on the relative distribution of KEGG Orthologs (KOs; Pielou's evenness) and number of KEGG Orthologs (KOs; Richness) within the communities; there were no significant differences in relation to irrigation treatment and no obvious clustering by potato stock. **B)** Principal coordinate analysis (PCoA) based on Bray-Curtis distance of functional beta diversity dissimilarity between the communities as assessed through PiCrust2 and hierarchical metastorms analysis; note that there was no distinct clustering in relation to any of the variables.

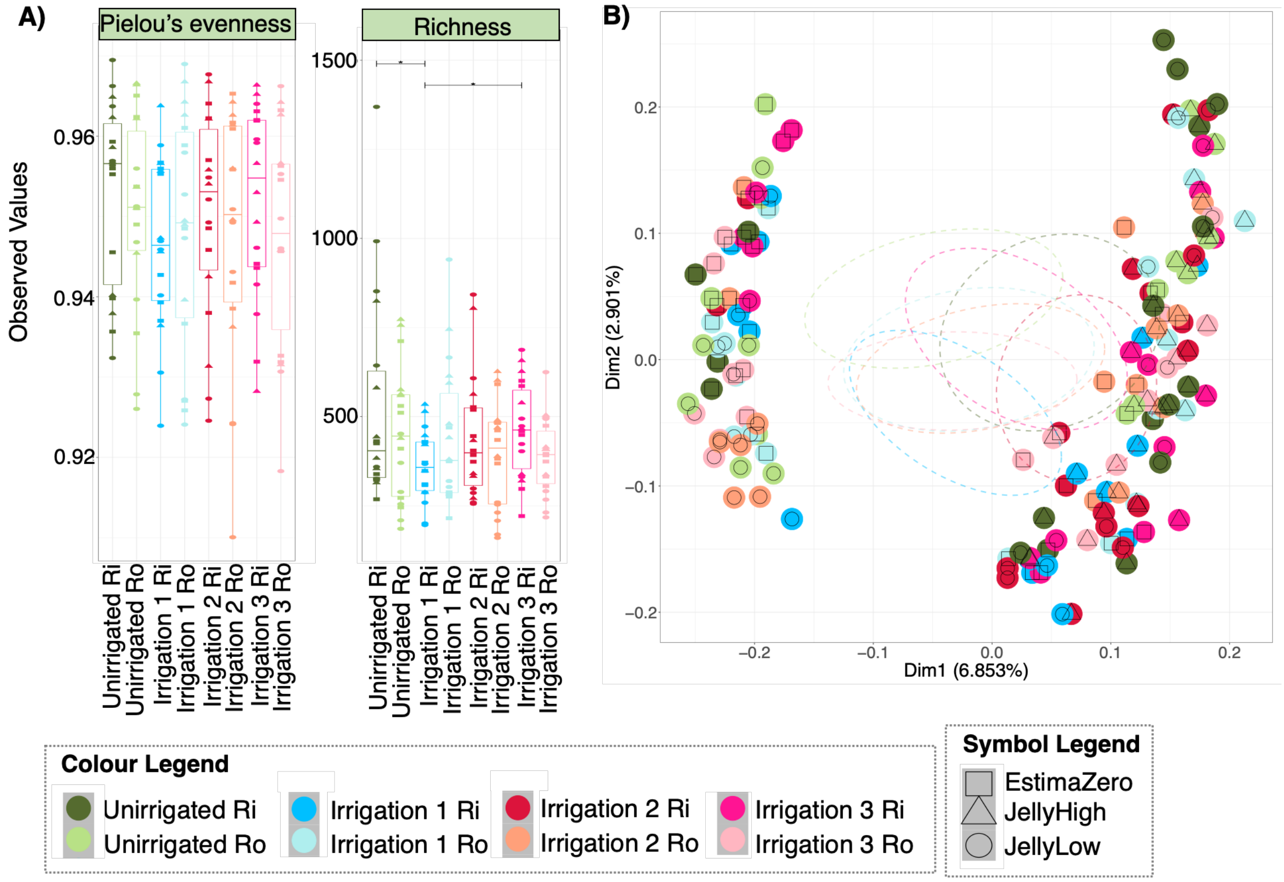


**Supplementary Figure 6. Taxonomic diversity of the soil microbiomes at the harvest (T_H) timepoint.** Colours represent the irrigation regime (Unirrigated, Irrigation 1, Irrigation 2, and Irrigation 3) and whether the sample was from the 'Ridge' or 'Root'. Symbols represent the potato seed stock (EstimaZero, JellyLow, and JellyHigh). **A)** Shows diversity based on the relative distribution of taxa (Pielou's evenness) and the number of different ASVs (Richness) within the communities; which were similar across irrigation regimes. **B)** Principal coordinate analysis (PCoA) based on Bray-Curtis distance of beta diversity dissimilarity between the communities; note that samples clustered according to potato stock, sample type (ridge or root) and Irrigation 2 samples showed some clustering.

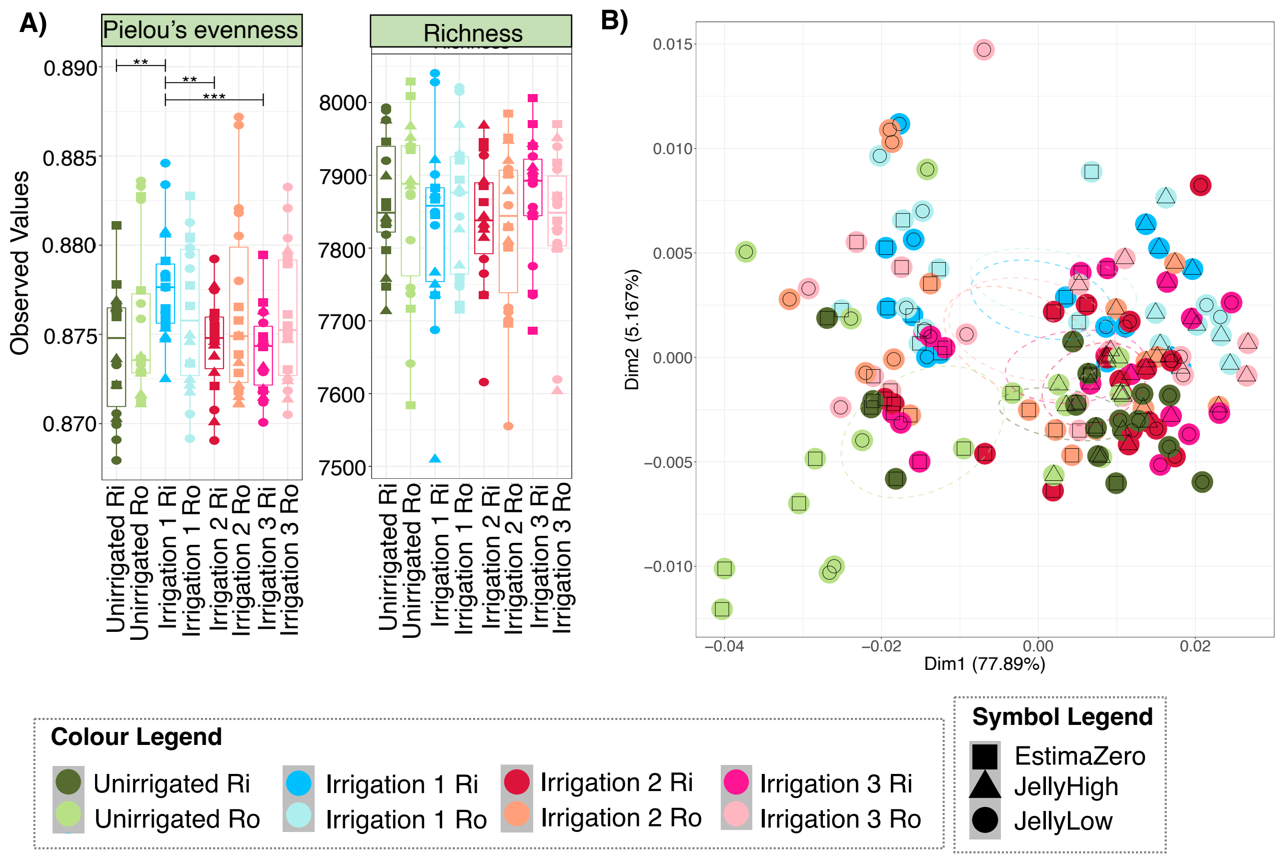


**Supplementary Figure 7. Functional diversity of the soil microbiomes at the harvest (T_H) timepoint.** Colours represent the irrigation regime (Unirrigated, Irrigation 1, Irrigation 2, and Irrigation 3) and the sample was from the 'Ridge' or 'Root'. Symbols represent the potato seed stock (EstimaZero, JellyLow, and JellyHigh). **A)** Shows diversity based on the relative distribution of KEGG Orthologs (KOs; Pielou's evenness) and number of KEGG Orthologs (KOs; Richness) within the communities; there were no significant differences in relation to irrigation treatment and no obvious clustering by potato stock. **B)** Principal coordinate analysis (PCoA) based on Bray-Curtis distance of functional beta diversity dissimilarity between the communities as assessed through PiCrust2 and hierarchical metastorms analysis; note that samples loosely cluster according to irrigation regime, sample type (root or ridge) and primarily cluster according to potato stock.

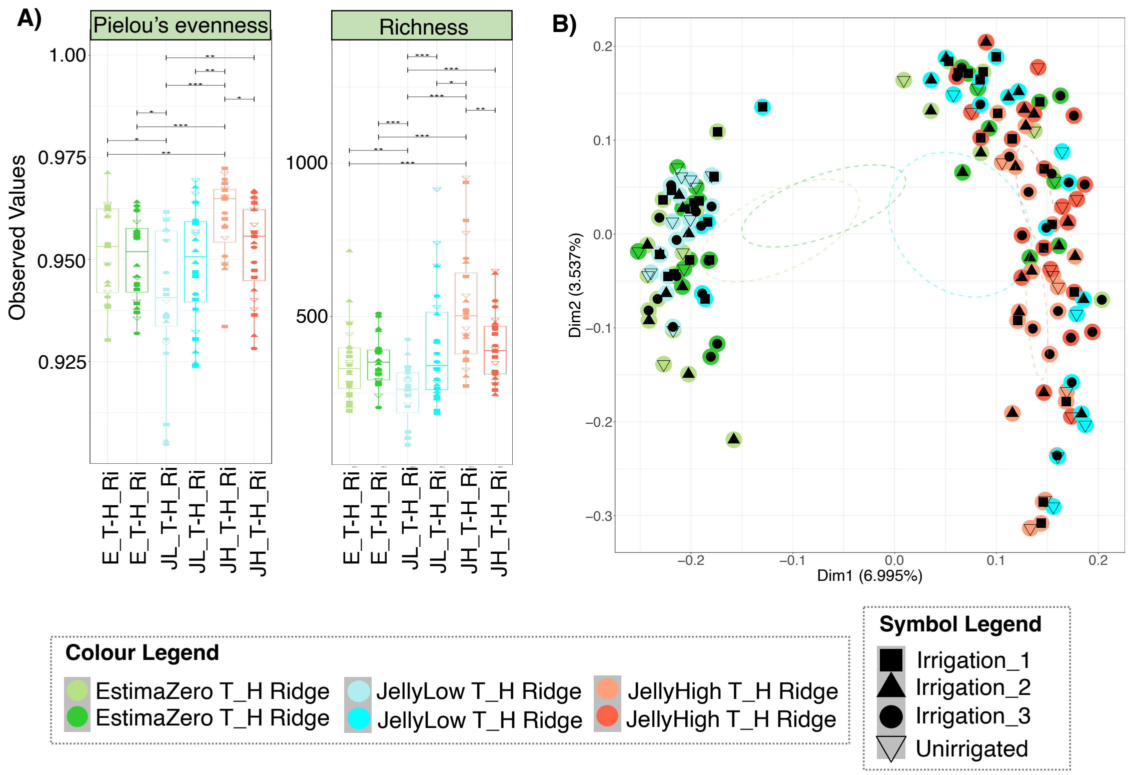


**Supplementary Figure 8. Taxonomic diversity of the soil microbiomes at the 50% plant emergence (T_E) and plant harvest (T_H) time points from 'Ridge' samples only.** Colours represent the irrigation regime (Unirrigated, Irrigation 1, Irrigation 2, and Irrigation 3). Symbols represent the potato seed stock (EstimaZero, JellyLow, and JellyHigh). **A)** Shows diversity based on the relative distribution of taxa (Pielou's evenness) and the number of different ASVs (Richness) within the communities; which were similar across irrigation regimes. **B)** Principal coordinate analysis (PCoA) based on Bray-Curtis distance of beta diversity dissimilarity between the communities; note that samples clustered according to potato stock.

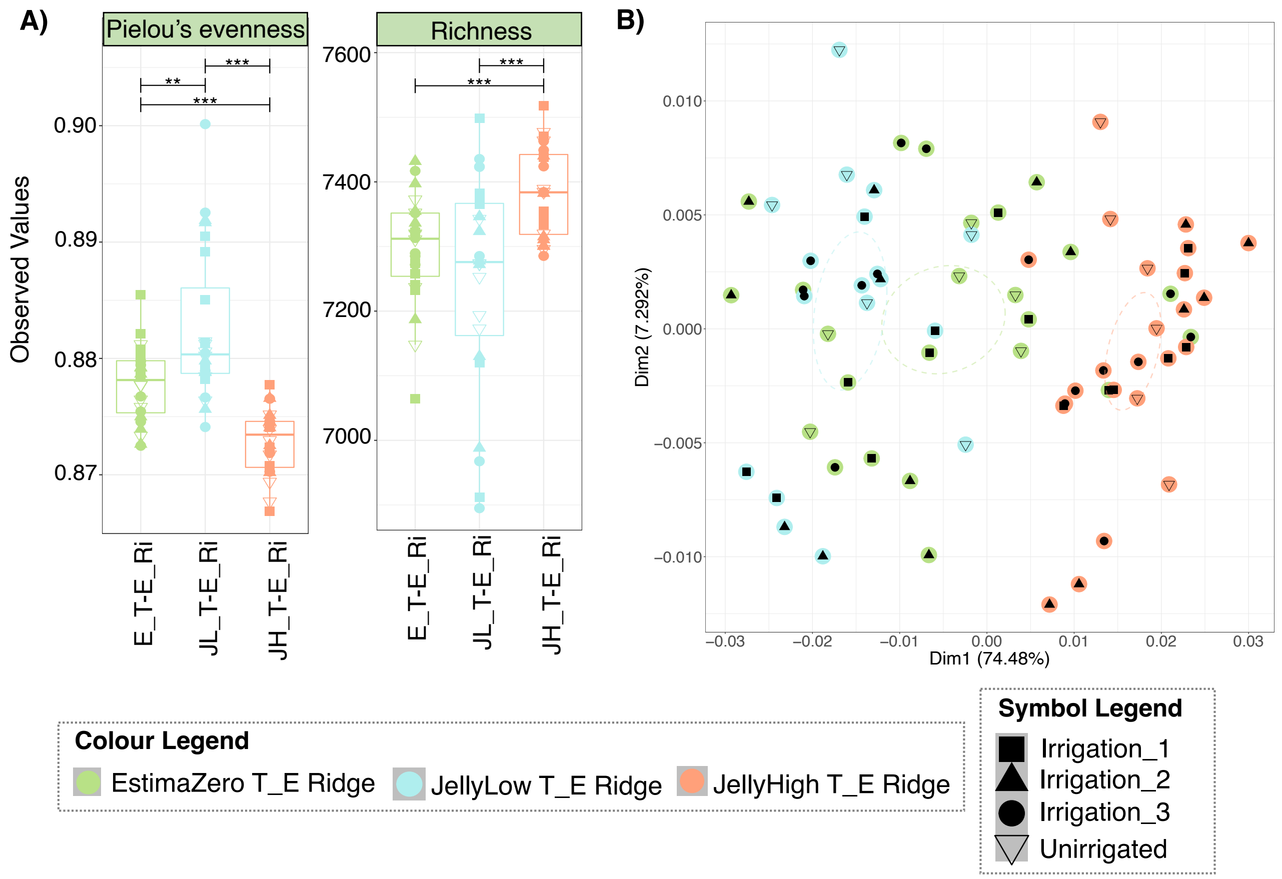


**Supplementary Figure 9. Functional diversity of the soil microbiomes at 50% plant emergence (T_E).** Colours represent potato stock (EstimaZero – E, JellyLow – JL, and JellyHigh – JH). Symbols represent the irrigation regimes (Unirrigated, Irrigation_1, Irrigation_2, and Irrigation 3). **A)** Shows diversity based on the relative distribution of KEGG Orthologs (KOs; Pielou's evenness) and number of KEGG Orthologs (KOs; Richness) within the communities; there were no significant differences in relation to irrigation treatment and no obvious clustering by potato stock. **B)** Principal coordinate analysis (PCoA) based on Bray-Curtis distance of functional beta diversity dissimilarity between the communities as assessed through PiCrust2 and hierarchical metastorms analysis; note that samples very loosely cluster according to potato stock.

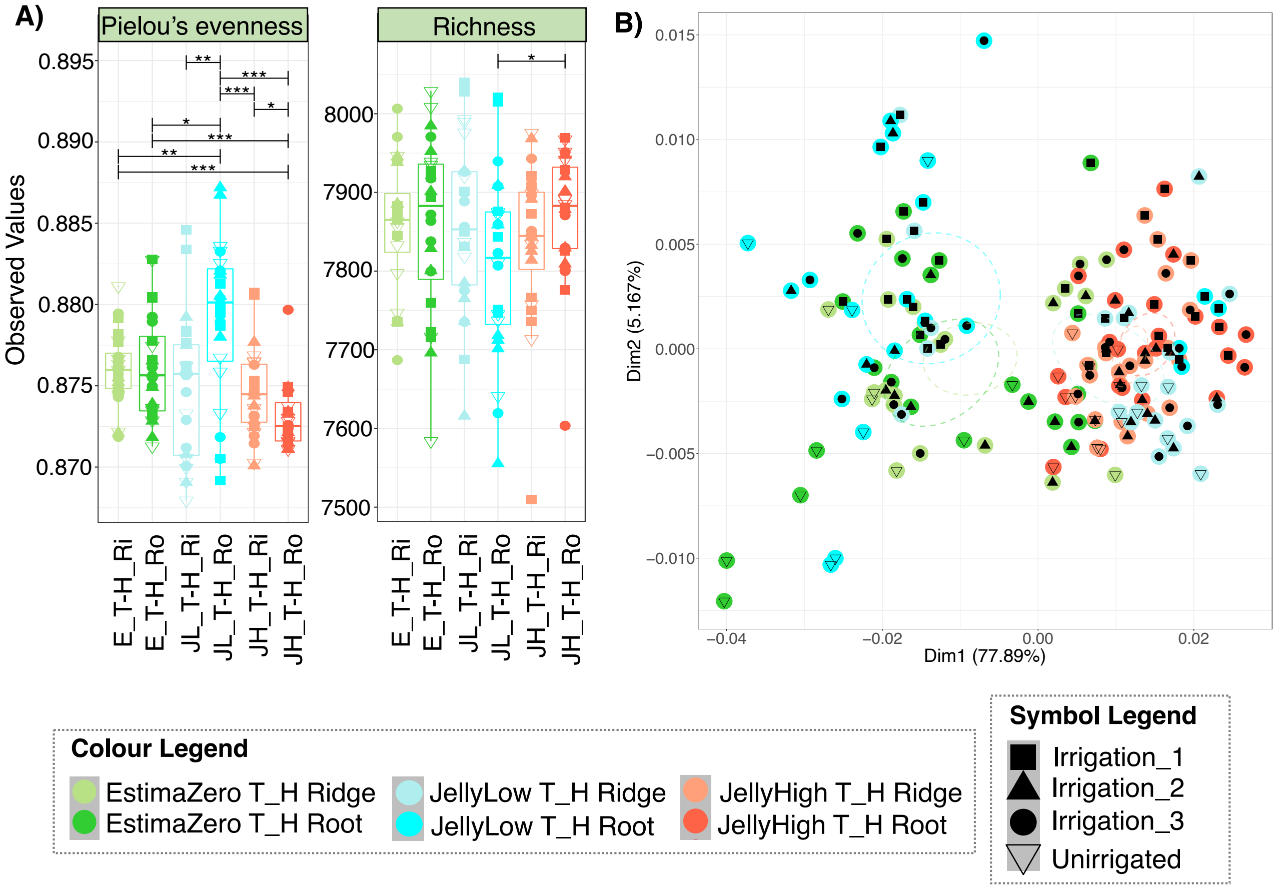


**Supplementary Figure 10. Functional diversity of the soil microbiomes at plant harvest (T_H).** Colours represent potato stock (EstimaZero – E, JellyLow – JL, and JellyHigh – JH), time-point (50% plant emergence – T_H), and soil sample type (Ridge – Ri, Root – Ro). Symbols represent the irrigation regimes (Unirrigated, Irrigation_1, Irrigation_2, and Irrigation 3). **A)** Shows diversity based on the relative distribution of KEGG Orthologs (KOs; Pielou's evenness) and number of KEGG Orthologs (KOs; Richness) within the communities; there were no significant differences in relation to irrigation treatment and no obvious clustering by potato stock. **B)** Principal coordinate analysis (PCoA) based on Bray-Curtis distance of functional beta diversity dissimilarity between the communities as assessed through PiCrust2 and hierarchical metastorms analysis; note that samples loosely cluster according to potato stock and sample type.

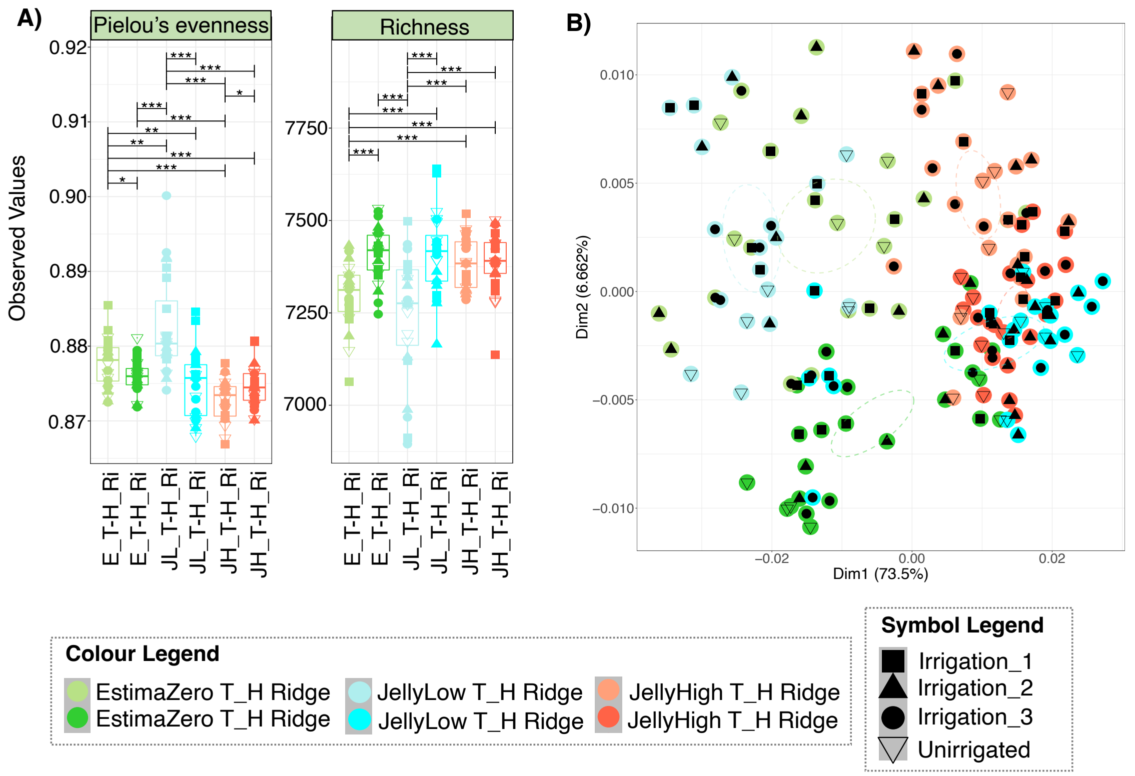


**Supplementary Figure 11. Functional diversity of the soil microbiomes at the 50% plant emergence (T_E) and plant harvest (T_H) time points from 'Ridge' samples only.** Colours represent the irrigation regime (Unirrigated, Irrigation 1, Irrigation 2, and Irrigation 3). Symbols represent the potato seed stock (EstimaZero, JellyLow, and JellyHigh). **A)** Shows diversity based on the relative distribution of KEGG Orthologs (KOs; Pielou's evenness) and number of KEGG Orthologs (KOs; Richness) within the communities; there were no significant differences in relation to irrigation treatment and no obvious clustering by potato stock. **B)** Principal coordinate analysis (PCoA) based on Bray-Curtis distance of functional beta diversity dissimilarity between the communities as assessed through PiCrust2 and hierarchical metastorms analysis; note that samples loosely cluster according to potato stock and time point.

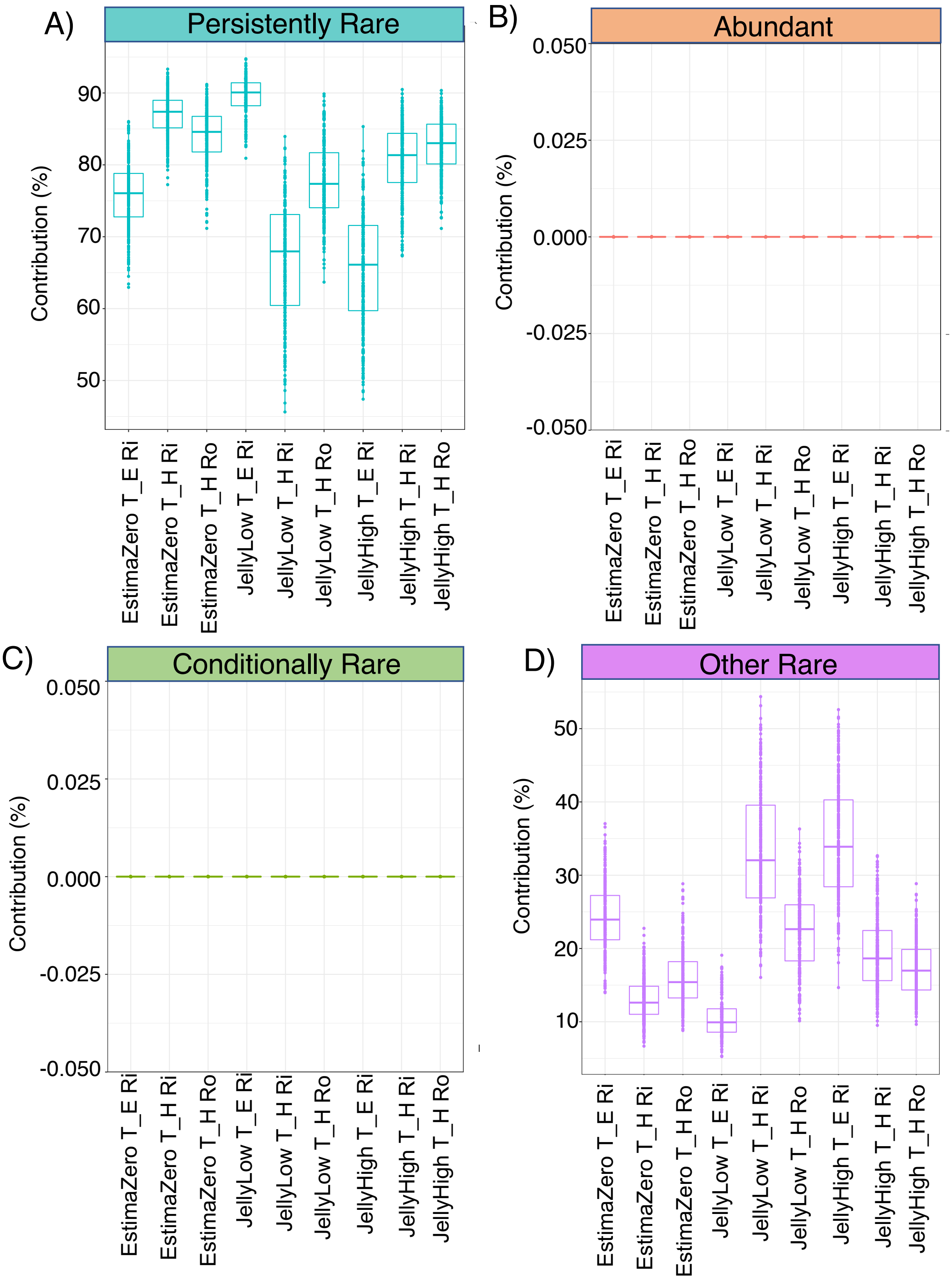


**Supplementary Figure 12.** Analysis of rare taxa from the Estima, JellyLow and JellyHigh samples at T_E (50% plant emergence) and T_H (plant harvest) divided into ridge and root samples (Ri and Ro). These figures show the percentage contribution (%) to taxonomic beta-diversity dissimilarity of: **A)** ‘Persistently Rare’ taxa **B)** ‘Abundant’ taxa; **C)** ‘Conditionally Rare’ taxa; and **D)** ‘Other Rare’ taxa.

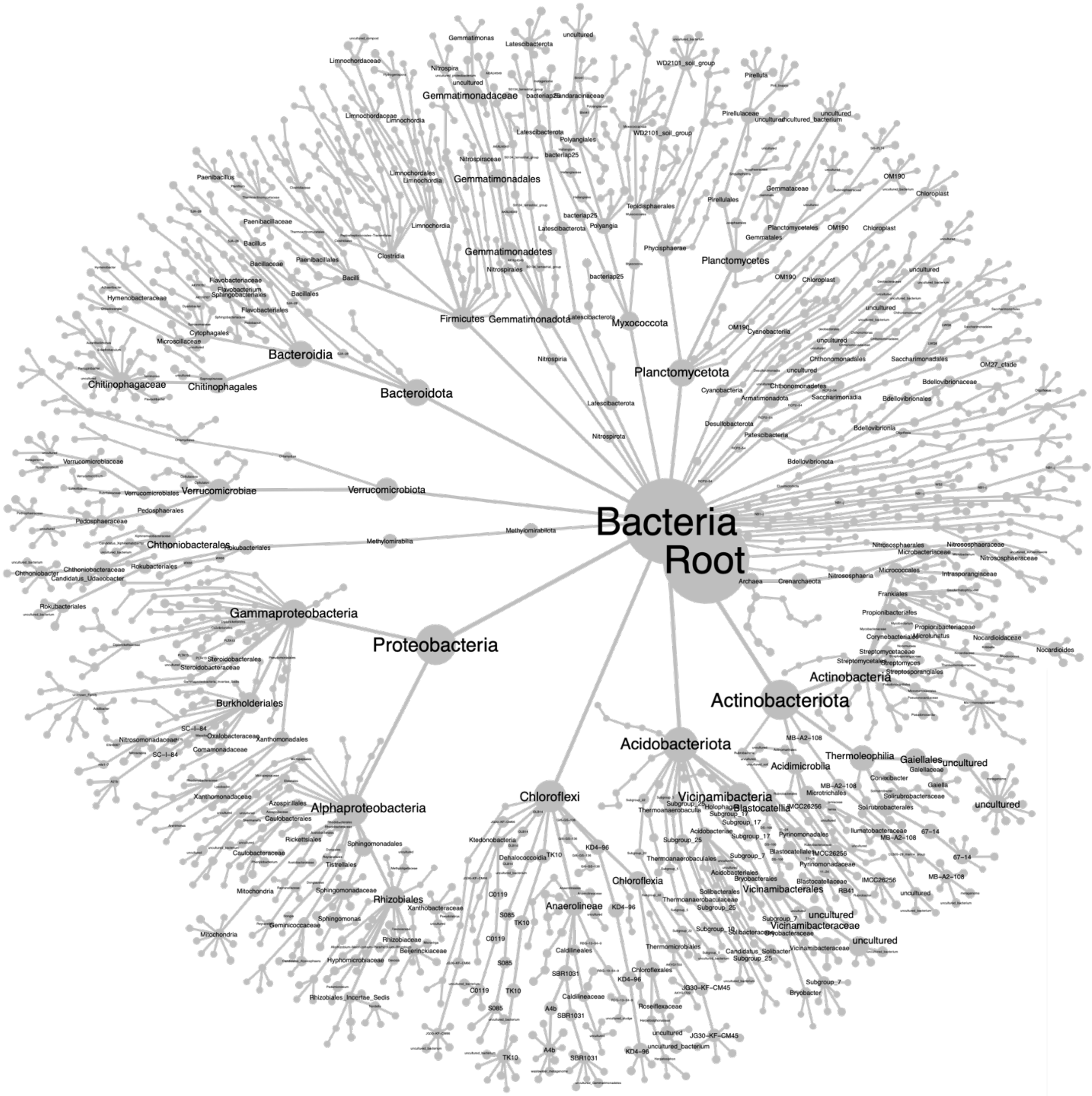


**Supplementary Figure 13. Taxonomic map for differential heat tree analysis.** The branches show the taxonomic identification of all ASVs. Note that the same tree can be used for each differential heat tree analysis instance.

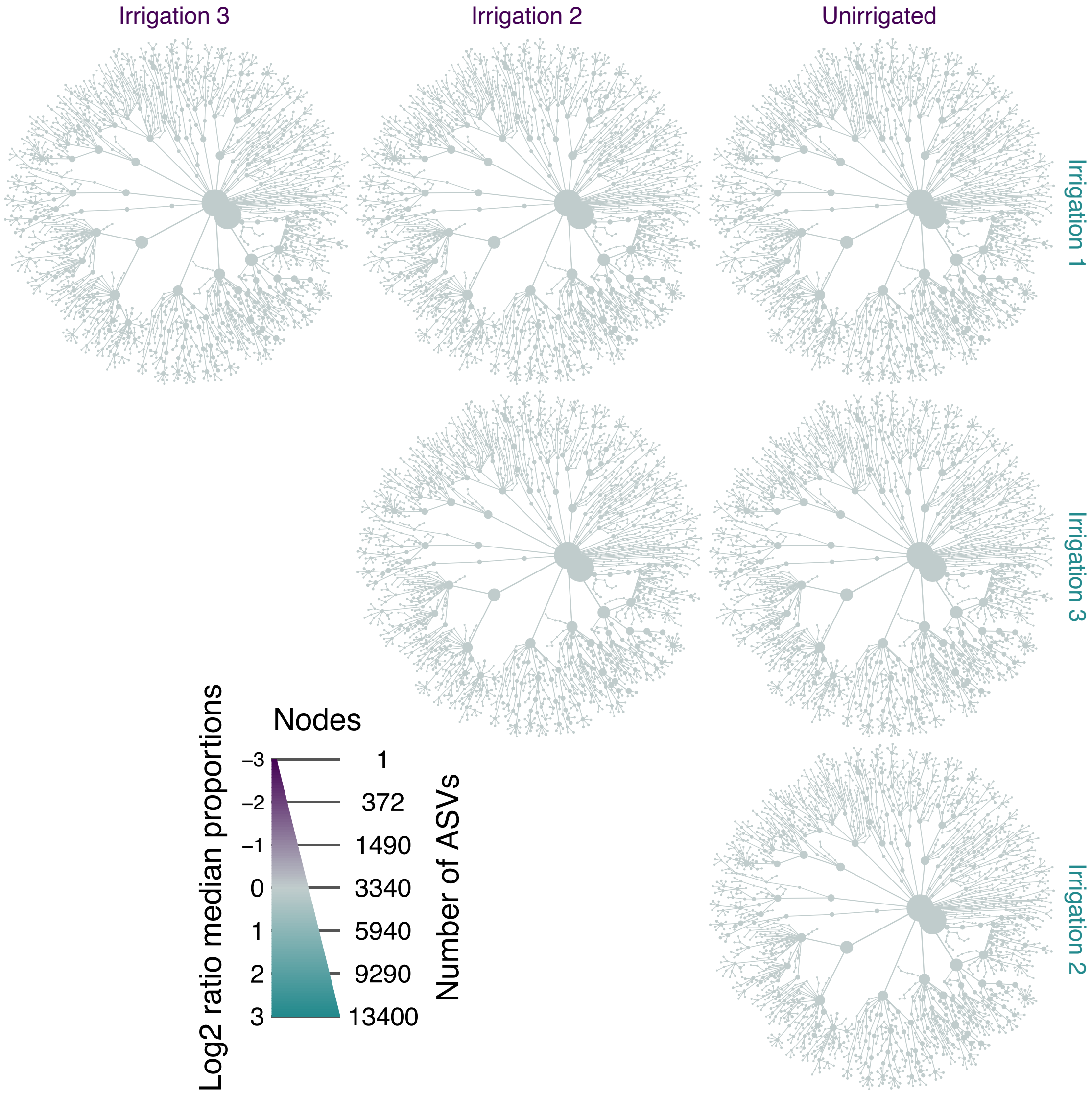


**Supplementary Figure 14. Heat tree analysis to visualise the differential taxa between irrigation regimes at 50% plant emergence (T_E).** The full legend for the tree branches is shown in Supplementary Figure 10. Each tree shows a pair-wise comparison between irrigation regimes (denoted by either purple or blue). The size of the branch nodes shows the number of ASVs, while the differential $Log_2$ ratio median proportion is shown by the colours. Note that there are no $Log_2$ fold differences between taxa across the irrigation regimes.

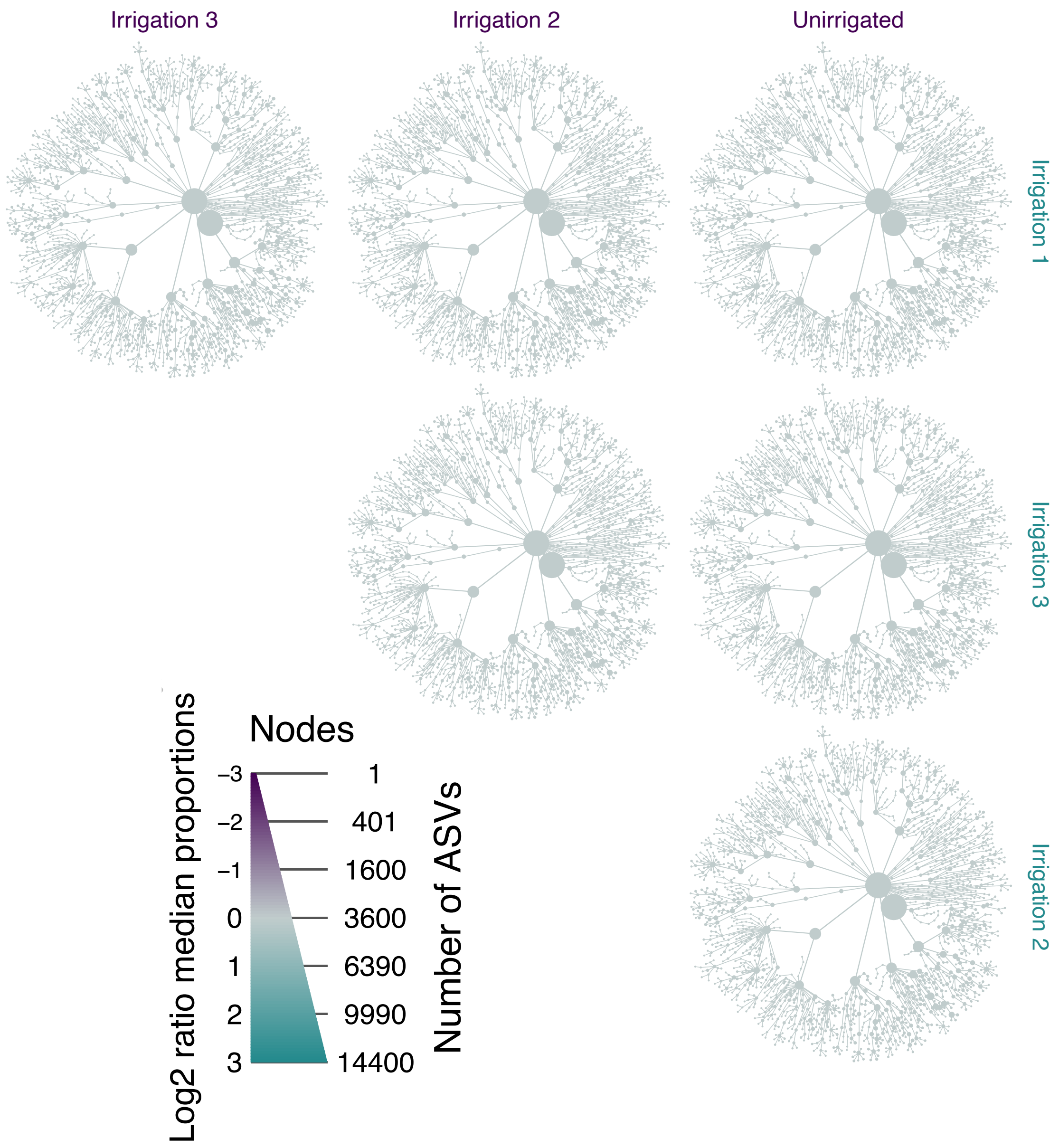


**Supplementary Figure 15. Heat tree analysis to visualise the differential taxa between irrigation regimes at potato harvest (T_H) in soil ridge samples.** The full legend for the tree branches is shown in Supplementary Figure 10. Each tree shows a pair-wise comparison between irrigation regimes (denoted by either purple or blue). The size of the branch nodes shows the number of ASVs, while the differential $Log_2$ ratio median proportion is shown by the colours. Note that there are no $Log_2$ fold differences between taxa across the irrigation regimes.

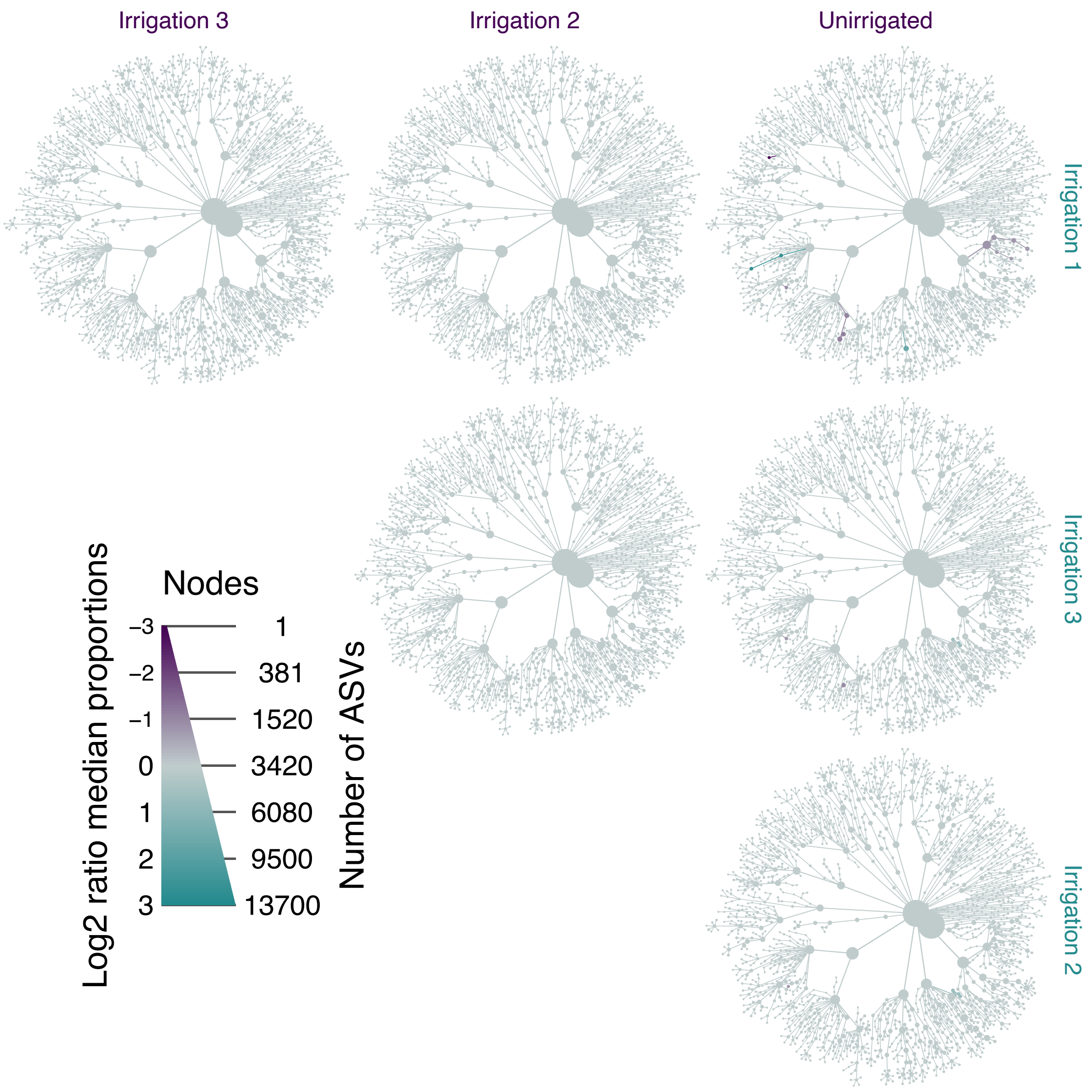


**Supplementary Figure 16. Heat tree analysis to visualise the differential taxa between irrigation regimes at potato harvest (T_H) in soil root samples.** The full legend for the tree branches is shown in Supplementary Figure 10. Each tree shows a pair-wise comparison between irrigation regimes (denoted by either purple or blue). The size of the branch nodes shows the number of ASVs, while the differential $Log_2$ ratio median proportion is shown by the colours. Note that there are no $Log_2$ fold differences between taxa across the irrigation regimes.

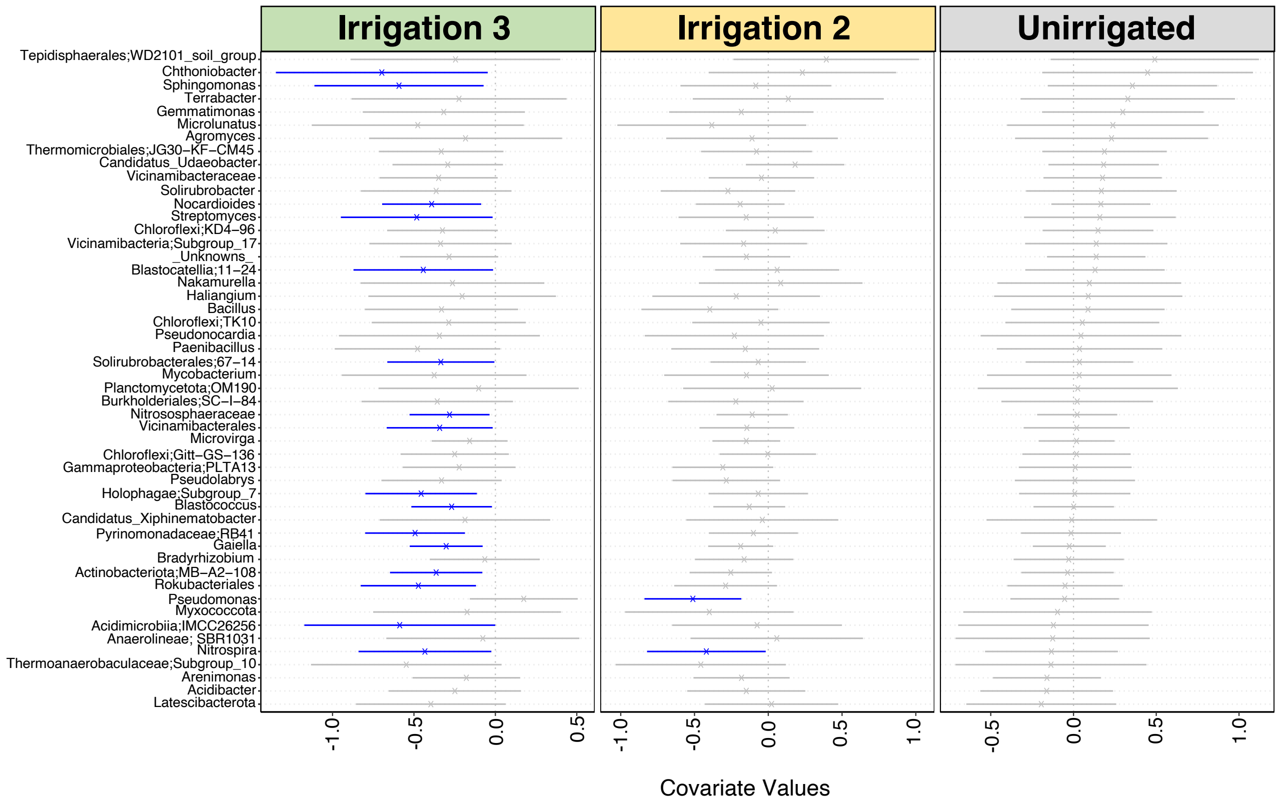


**Supplementary Figure 17. Plot highlighting the impact of 'Irrigation Regime' on the abundance of microbial taxa in the potato harvest (T_H) ridge soil samples, based on the GLVMM analysis.** The microbial genera names are indicated on the y-axis. The environment covariates values are indicated on the x-axis which are the coefficient values against the environmental covariates (e.g. Unirrigated, Irrigation_2, and Irrigation_3 as compared to the Irrigation 1 treatment). The values range from positive, neutral to negative associations with the specific taxa on the y-axis. The red lines show significantly positive taxa associated with the indicated environmental covariate and the blue lines show significantly negative taxa; grey lines are not statistically significant. Note: Irrigation 1 treatment is not displayed as the analysis uses this treatment as the reference.

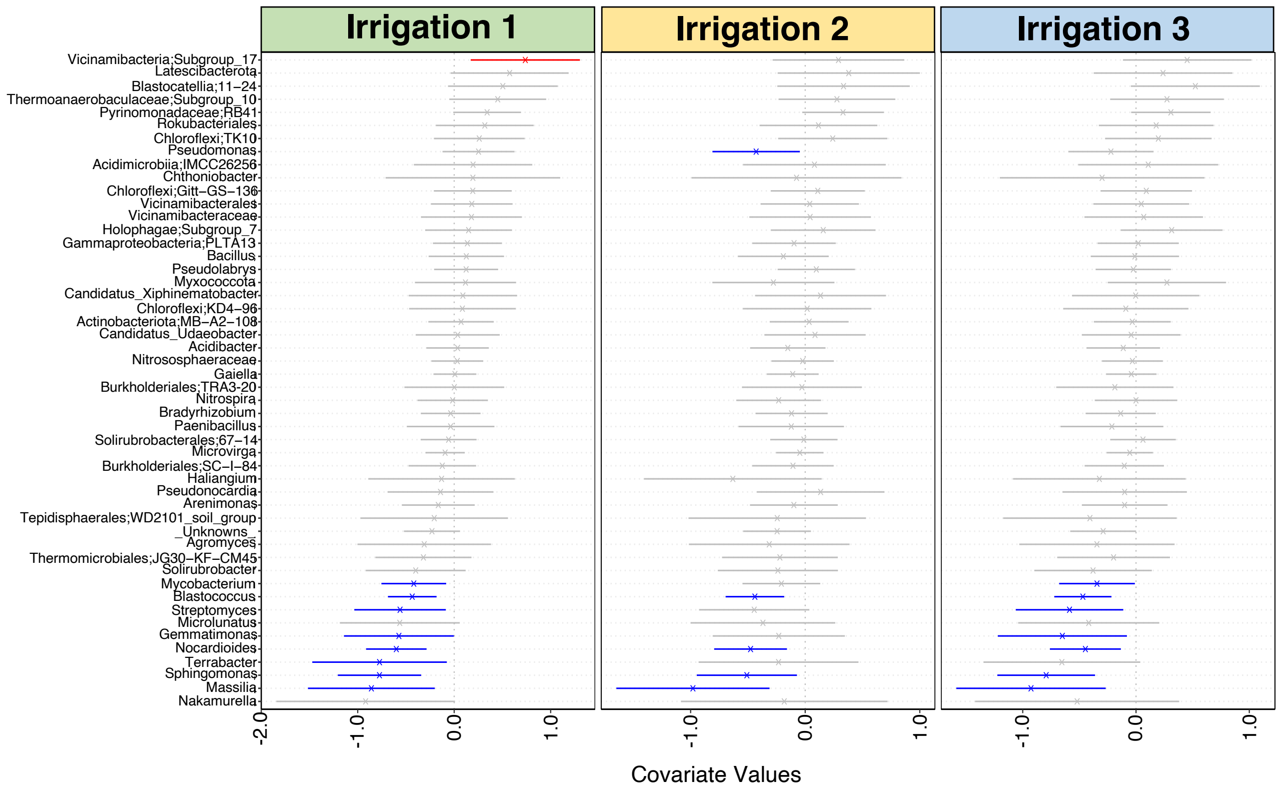


**Supplementary Figure 18. Plot highlighting the impact of 'Irrigation Regime' on the abundance of microbial taxa in the potato harvest (T_H) root soil samples, based on the GLVMM analysis.** The microbial genera names are indicated on the y-axis. The environment covariates values are indicated on the x-axis which are the coefficient values against the environmental covariates (e.g. Irrigation_1, Irrigation_2, and Irrigation_3, as compared to the Unirrigated treatment). The values range from positive, neutral to negative associations with the specific taxa on the y-axis. The red lines show significantly positive taxa associated with the indicated environmental covariate and the blue lines show significantly negative taxa; grey lines are not statistically significant. Note: Unirrigated treatment is not displayed as the analysis uses this as the reference.

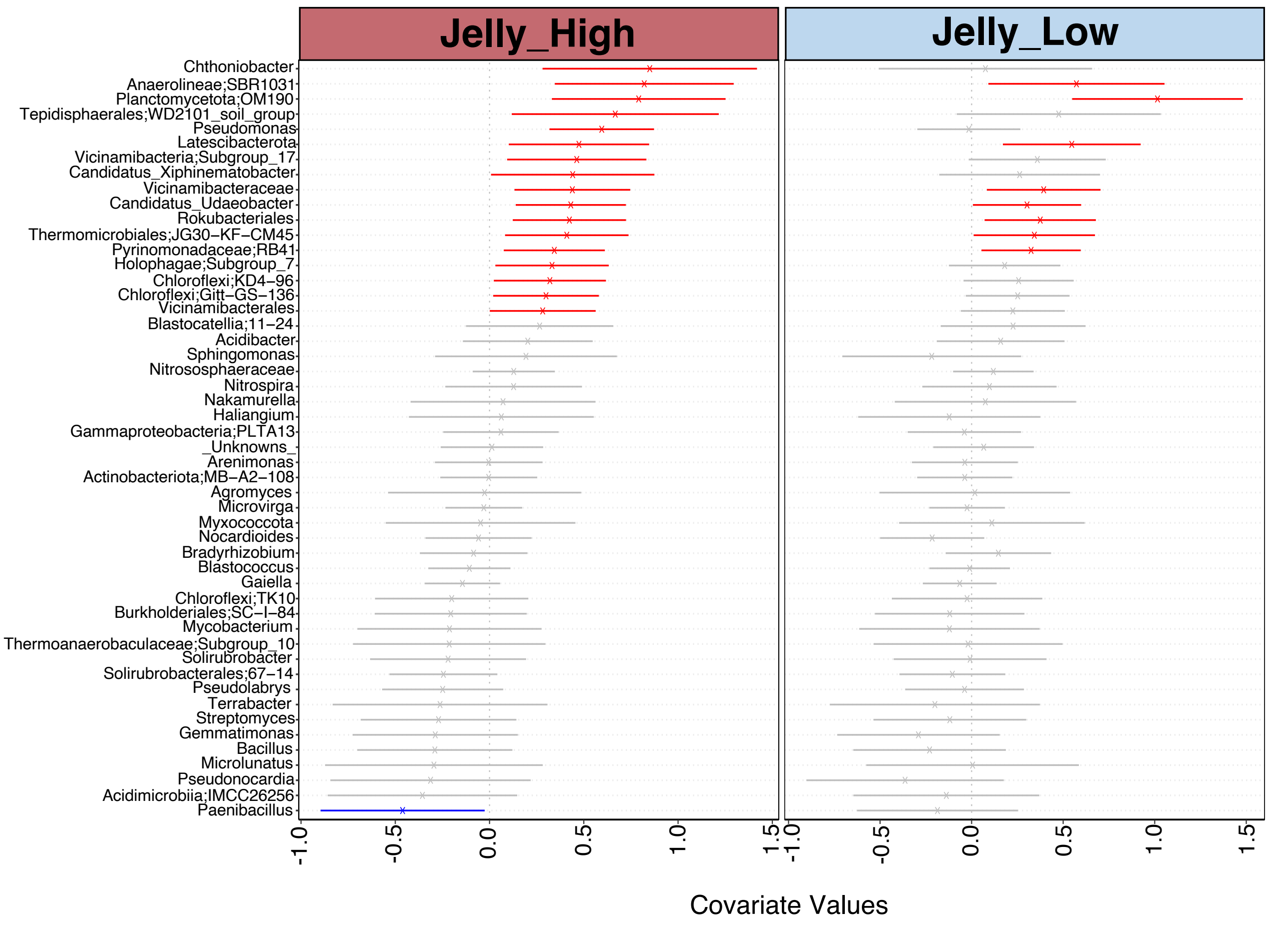


**Supplementary Figure 19. Plot highlighting the impact of 'Potato Stock' on the abundance of microbial taxa at 50% plant emergence (T_E), based on the GLVMM analysis.** The microbial genera names are indicated on the y-axis. The environmental covariates values are indicated on the x-axis which are the coefficient values against the environmental covariates (e.g. 'Potato Stock' either JellyHigh, JellyLow as compared to the EstimaZero. The values range from positive, and neutral to negative associations with the specific taxa on the y-axis. The red lines show significantly positive taxa associated with the indicated environmental covariate and the blue lines show significantly negative taxa; grey lines are not statistically significant. Note: The EstimaZero group is not displayed as the analysis uses this as the reference.

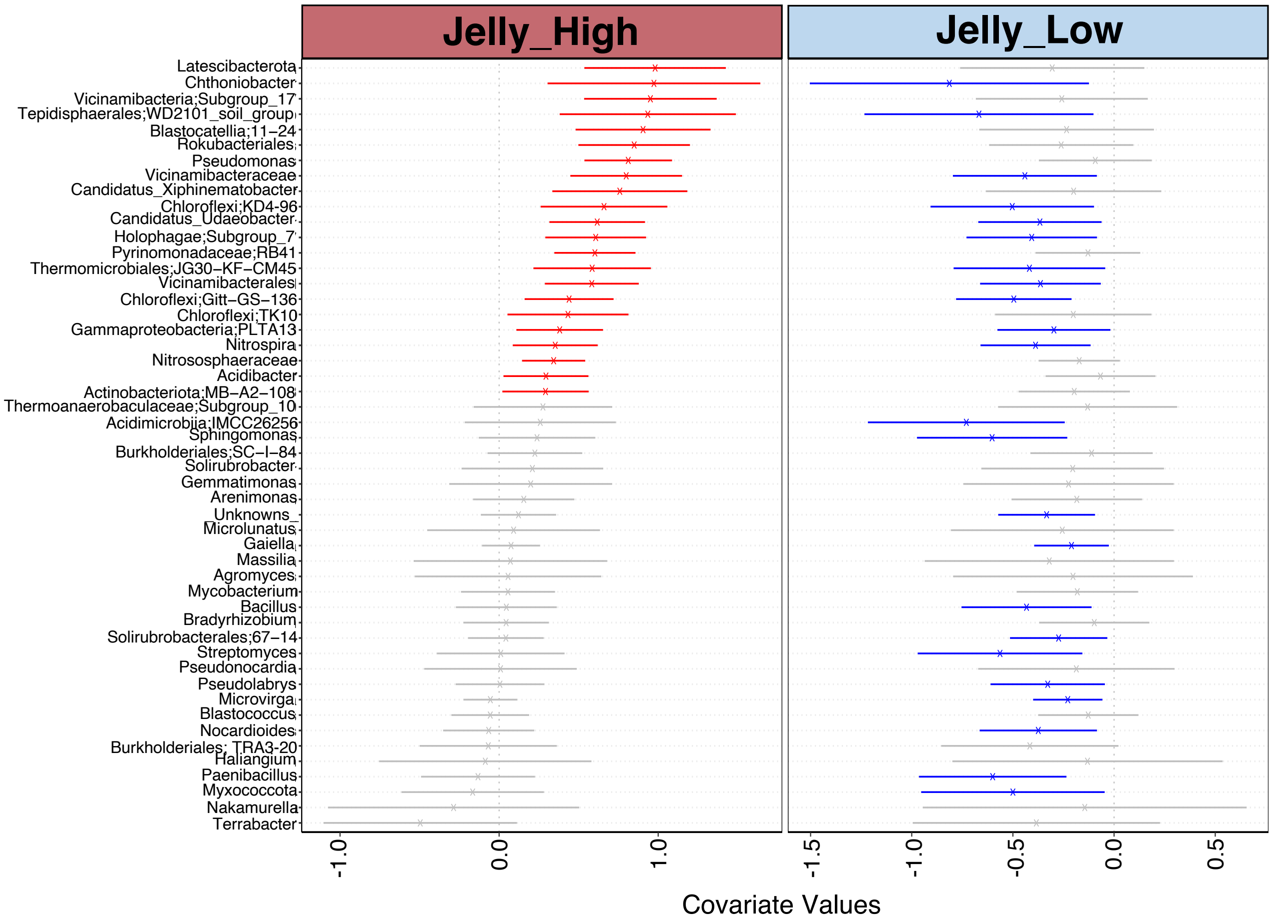


**Supplementary Figure 20. Plot highlighting the impact of 'Potato Stock' on the abundance of microbial taxa in the root samples at potato harvest (T_H), based on the GLVMM analysis.** The microbial genera names are indicated on the y-axis. The environment covariates values are indicated on the x-axis which are the coefficient values against the environmental covariates (e.g. 'Potato Stock' either JellyHigh, JellyLow as compared to the EstimaZero. The values range from positive, neutral to negative associations with the specific taxa on the y-axis. The red lines show significantly positive taxa associated with the indicated environmental covariate and the blue lines show significantly negative taxa; grey lines are not statistically significant. Note: The EstimaZero group is not displayed as the analysis uses this as the reference.

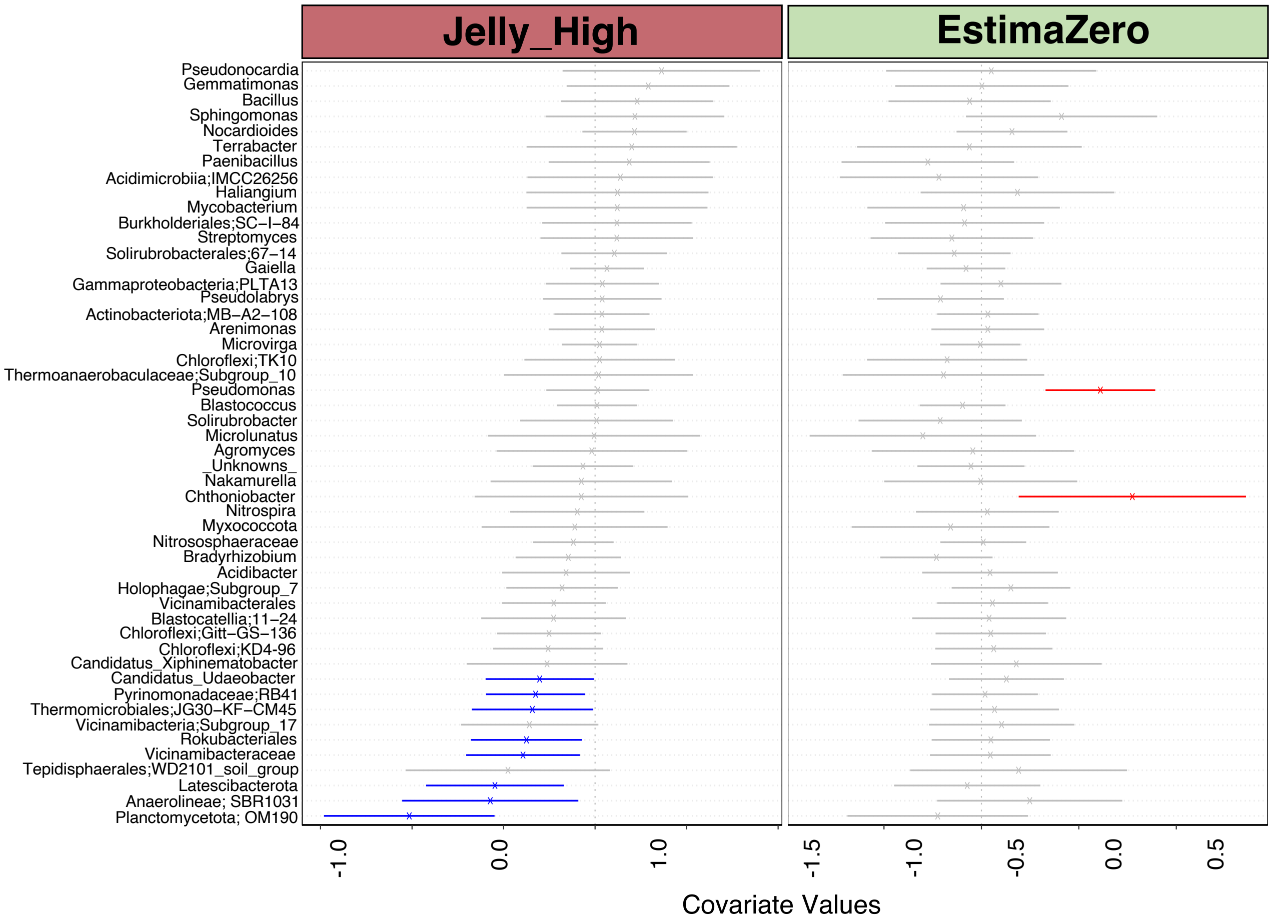


**Supplementary Figure 21. Plot highlighting the impact of 'Potato Stock' on the abundance of microbial taxa in the ridge samples at potato harvest (T_H), based on the GLVMM analysis.** The microbial genera names are indicated on the y-axis. The environment covariates values are indicated on the x-axis which are the coefficient values against the environmental covariates (e.g. 'Potato Stock' either JellyHigh, EstimaZero as compared to the JellyLow. The values range from positive, neutral to negative associations with the specific taxa on the y-axis. The red lines show significantly positive taxa associated with the indicated environmental covariate and the blue lines show significantly negative taxa; grey lines are not statistically significant. Note: The JellyLow group is not displayed as the analysis uses this as the reference.

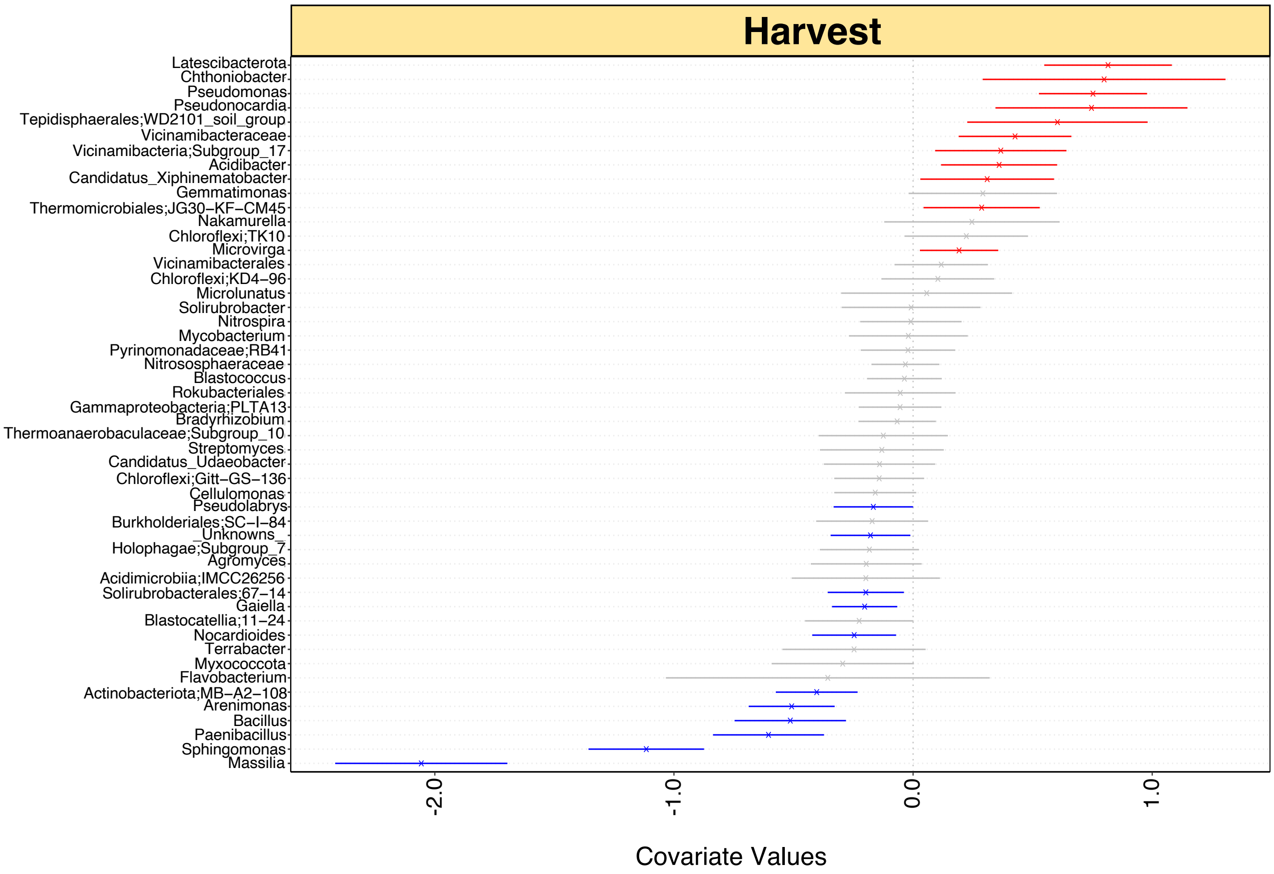


**Supplementary Figure 22. Plot highlighting the impact of time on the abundance of microbial taxa in the ridge soil samples, based on the GLVMM analysis.** The microbial genera names are indicated on the y-axis. The environment covariates values are indicated on the x-axis which are the coefficient values against the environmental covariate (e.g. Harvest [T_H] time point, as compared to the 50% plant emergence [T_E] time point. The values range from positive, neutral to negative associations with the specific taxa on the y-axis. The red lines show significantly positive taxa associated with the indicated environmental covariate and the blue lines show significantly negative taxa; g grey lines are not statistically significant. Note: The 50% emergence time point (T_E) is not displayed, as the analysis uses this as the reference.

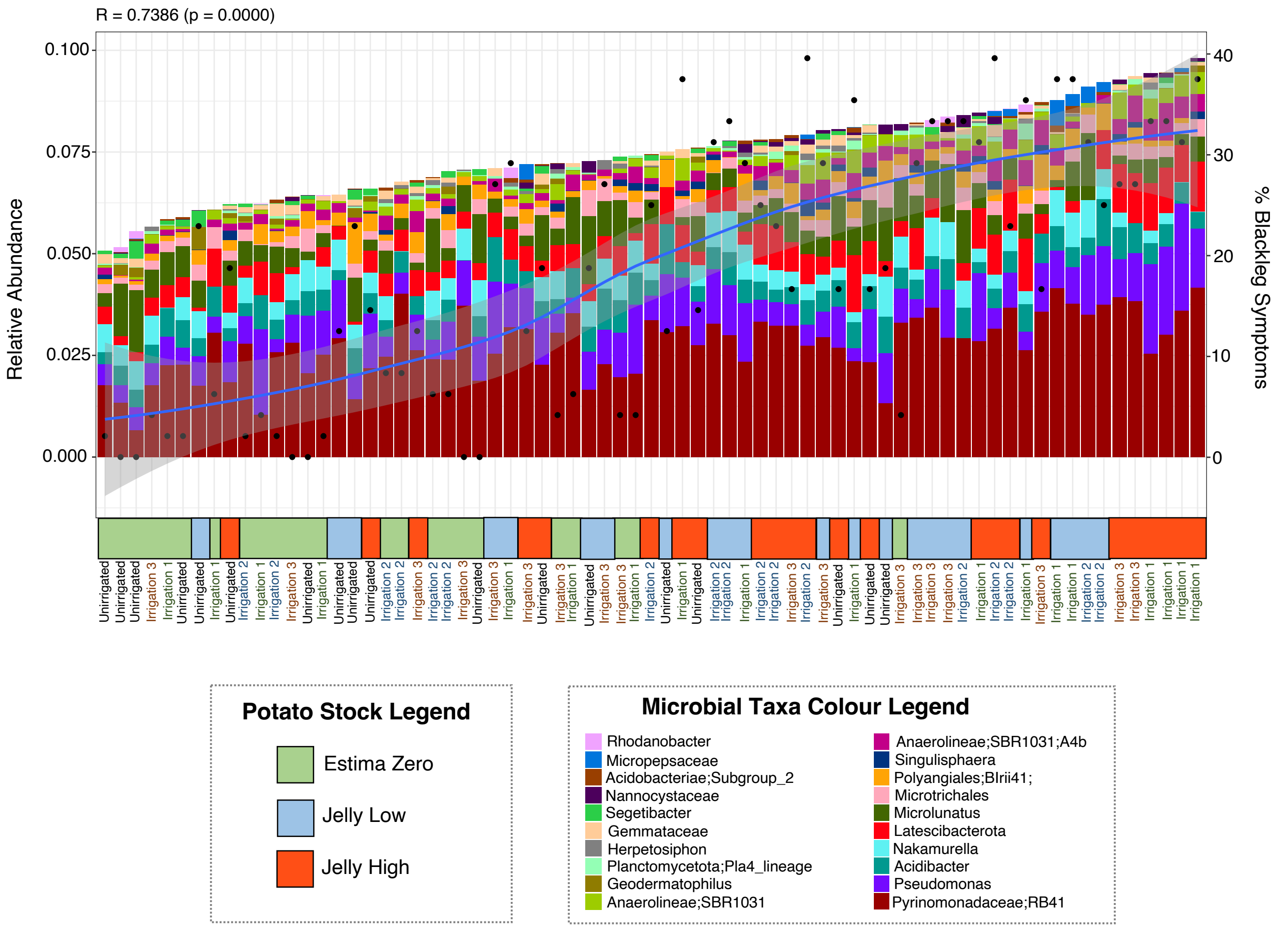


**Supplementary Figure 23. Minimum subset of amplicon sequencing variants (ASVs) associated with percentage blackleg symptoms during the field trial.** Analysis using the Ensemble Quotient Optimisation (EQO) technique considering at most 20 microbial taxa and the root samples at harvest. The left y-axis shows the relative abundance of the microbial taxa, and the right y-axis shows the percentage blackleg symptoms observed. The x-axis shows the sample identity with a colour legend for potato stock (Green EstimaZero, Blue JellyLow, and Red JellyHigh) and the irrigation regimes are shown in text (Unirrigated, Irrigation 1, Irrigation 2, and Irrigation 3).

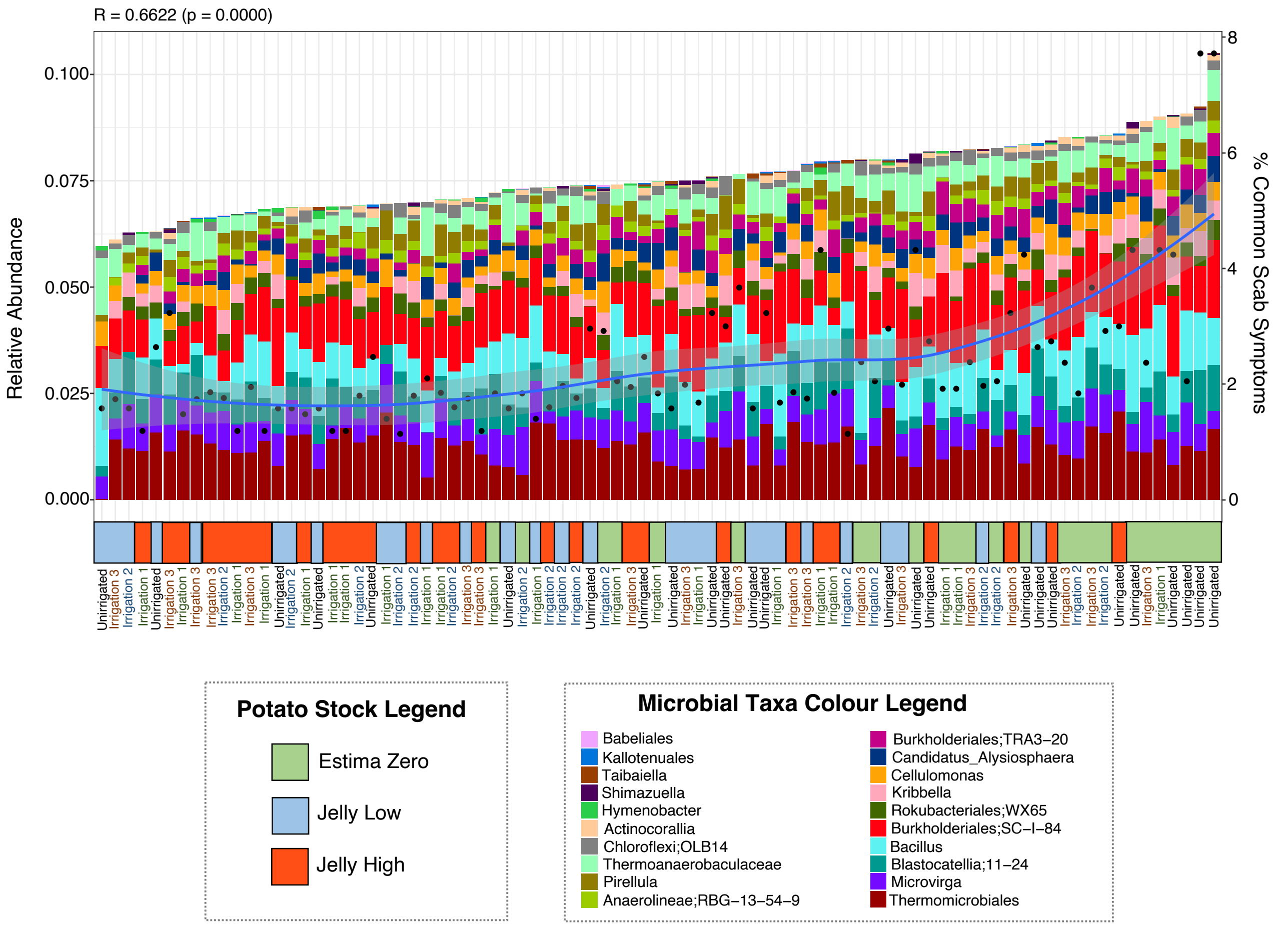


**Supplementary Figure 24. Continuous ensemble quotient analysis (EQO) of the harvest ridge samples.** Plot shows a stacked bar chart showing the minimum subset of amplicon sequencing variants (ASVs) associated with common scab symptoms. The x-axis shows the sample details including the potato stock (EstimaZero – Green, JellyLow – Blue, and JellyHigh – Red) and the irrigation regime is indicated in text. The right y-axis shows the relative abundance of the ASVs and the right y-axis shows the percentage (%) of common scab symptoms.

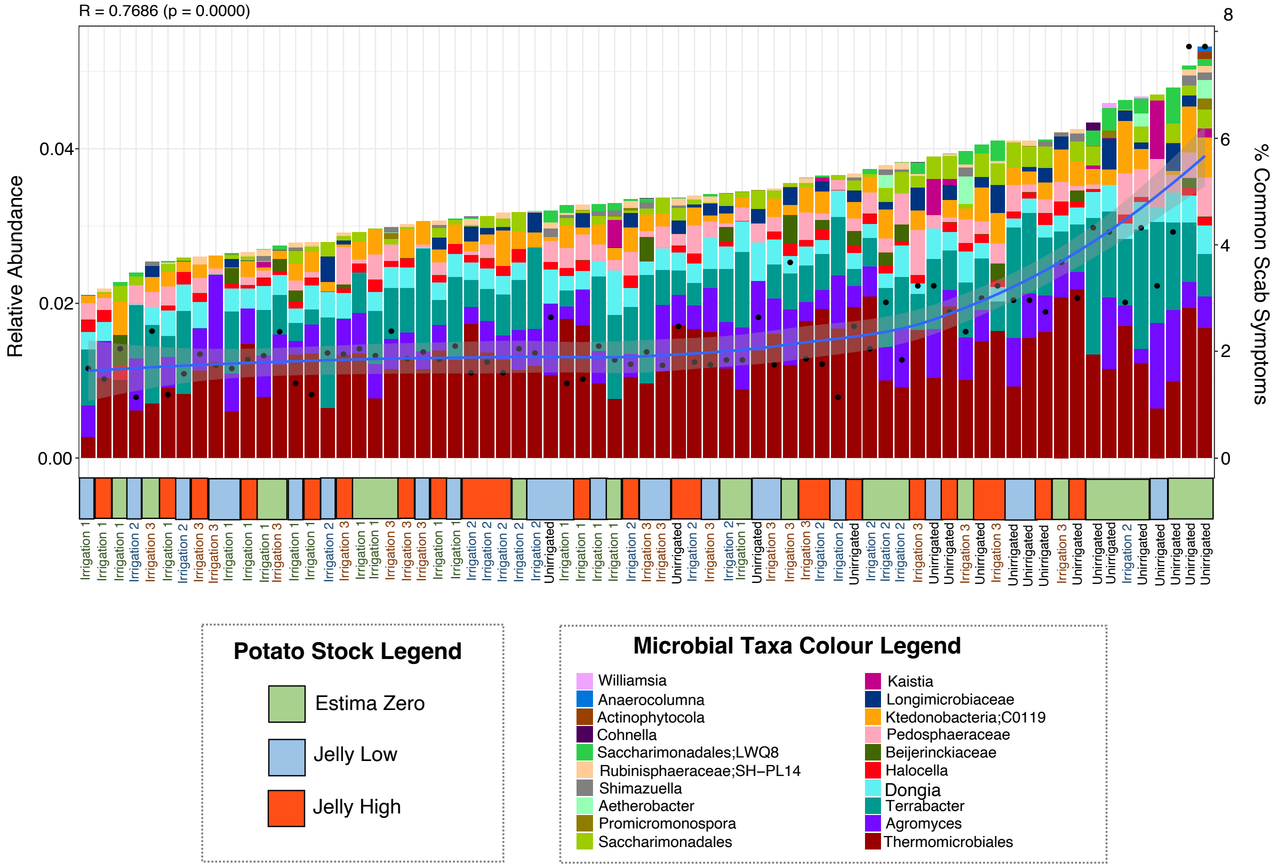


**Supplementary Figure 25. Continuous ensemble quotient analysis (EQO) of the harvest root samples.** Plot shows a stacked bar chart showing the minimum subset of amplicon sequencing variants (ASVs) associated with common scab symptoms. The x-axis shows the sample details including the potato stock (EstimaZero – Green, JellyLow – Blue, and JellyHigh – Red) and the irrigation regime is indicated in text. The right y-axis shows the relative abundance of the ASVs and the right y-axis shows the percentage (%) of common scab symptoms.

Paradis, E., Blomberg, S., Bolker [aut, B., cph, Brown, J., Claramunt, S., Claude, J., Cuong, H.S., Desper, R., Didier, G., Durand, B., Dutheil, J., Ewing, R.J., Gascuel, O., Guillerme, T., Heibl,

C., Ives, A., Jones, B., Krah [aut, F., cph, Lawson, D., Lefort, V., Legendre, P., Lemon, J., Louvel, G., Marcon [aut, E., cph, McCloskey, R., Nylander, J., Opgen-Rhein, R., Popescu, A.-A., Royer-Carenzi, M., Schliep, K., Strimmer, K., Vienne, D. de, 2023. ape: Analyses of Phylogenetics and Evolution.

Shan, X., Goyal, A., Gregor, R., Cordero, O.X., 2023. Annotation-free discovery of functional groups in microbial communities. Nat Ecol Evol 1–9. https://doi.org/10.1038/s41559-023-02021-z

Shan, X., Goyal, A., Gregor, R., Cordero, O.X., 2022. Annotation-free discovery of functional groups in microbial communities (preprint). Ecology. https://doi.org/10.1101/2022.08.02.502537

Simmel, A., 1958. Review of Non-Parametric Statistics for the Behavioral Sciences. American Journal of Sociology 63, 442–443.

Vass, M., Székely, A.J., Lindström, E.S., Langenheder, S., 2020. Using null models to compare bacterial and microeukaryotic metacommunity assembly under shifting environmental conditions. Scientific Reports 10. https://doi.org/10.1038/s41598-020-59182-1

Whitley, E., Ball, J., 2002. Statistics review 6: Nonparametric methods. Crit Care 6, 509. https://doi.org/10.1186/cc1820

Yang, S., Winkel, M., Wagner, D., Liebner, S., 2017. Community structure of rare methanogenic archaea: insight from a single functional group. FEMS Microbiology Ecology 93, fix126. https://doi.org/10.1093/femsec/fix126

Zhang, Y., Jing, G., Chen, Y., Li, J., Su, X., 2021. Hierarchical Meta-Storms enables comprehensive and rapid comparison of microbiome functional profiles on a large scale using hierarchical dissimilarity metrics and parallel computing. Bioinformatics Advances 1, vbab003. https://doi.org/10.1093/bioadv/vbab003
